# Adaptive challenges of past and future invasion of *Drosophila suzukii*: insights from novel genomic resources and statistical methods combining individual and pool sequencing data

**DOI:** 10.1101/2024.10.11.617812

**Authors:** Louise Camus, Nicolas Rode, Svitlana Serga, Anne Loiseau, Xiao Chen, Carole Iampietro, Marc Kenis, William Marande, Julián Mensch, Hugues Parinello, Marija Savić Veselinović, Sophie Valière, Jinping Zhang, Arnaud Estoup, Simon Boitard, Mathieu Gautier

## Abstract

Global change is accelerating biological invasions, making it crucial to understand how species adapt in new environments to improve management strategies. Genomic data provide valuable insights into adaptation through Genotype-Environment Association (GEA) studies, which identify genes and biological processes tied to invasion success, and through geometric Genomic Offset (gGO) statistics, which estimate genetic (mal)adaptation to new environments. Here, we investigate genetic adaptation in the invasive pest *Drosophila suzukii* using novel genomic resources and statistical methods. We use a new chromosome-level genome assembly and data from 37 populations, combining publicly available and newly generated pooled and individual sequencing data, analyzed with an enhanced version of BayPass software, tailored for such hybrid datasets. First, we identify genomic regions showing genetic differentiation between native and invasive populations. Then, using a GEA with 29 environmental covariates, we estimate the gGO between the source environments and the invaded areas, shedding light on the potential adaptive challenges *D. suzukii* faced during previous invasions. In addition, we estimate gGO for geographical areas not yet invaded to predict future invasion risks, and identify regions from which preadapted populations may originate. Our results reveal numerous genomic regions associated with the invasive status from genome scans. However, when considering broader patterns of adaptation to specific environmental variables through gGO analyses, we find that *D. suzukii* populations likely faced only limited adaptive challenges across their major invasion range, while certain uninvaded regions still remain at high risk of future invasion. Our study offers significant insights into *D. suzukii* adaptation and provides a practical population genomics framework to predict biological invasions, applicable to various species.

## Introduction

Invasive species can have significant negative impacts on biodiversity (Mollot *et al*., 2017), economy (Haubrock *et al*., 2021; Bradshaw et al., 2016), human health (Mazza *et al*., 2014), and food security (Bruce, 2010; Paini *et al*., 2016), with the total reported costs recently estimated at more than a trillion US dollars over the past few decades (Diagne *et al*., 2021). From an evolutionary perspective, invasive species raise questions about how populations can rapidly colonize new environments and the extent to which their genetic makeup is critical. One hypothesis suggests that invasive success may be due to one or more *de novo* advantageous mutations (Colautti and Barrett, 2013), although the short evolutionary timescale of biological invasions makes this hypothesis less likely (Barrett and Schluter, 2008). Alternatively, invasive success may be due to various forms of pre-adaptation, enabled by standing genetic variation in the native range that allows the exploitation of a wide range of environments, including anthropized ones (Estoup *et al*., 2016; Hufbauer et al., 2012).

With the advent of next-generation genotyping and sequencing approaches, genome-environment association (GEA) studies have proven informative in unraveling the genetic mechanisms involved in local adaptation, including to invaded environments. By identifying genomic regions whose allele frequencies covary with environmental factors (Yang *et al*., 2022; Hofmeister et al., 2021; Ma et al., 2020) or phenotypic traits (Turner *et al*., 2021; Olazcuaga *et al*., 2020; Pfenninger *et al*., 2021), they can provide insights into the biological processes that enable genetic adaptation. More generally, since GEA models summarize the relationship between the (adaptive) genomic composition of populations and their environment, they can be used to predict maladaptation to novel environments. Fitzpatrick and Keller (2015) introduced the Genomic Offset statistic to quantify the gap between the optimal genetic configuration for a new environment (as predicted by the GEA model) and the actual genetic makeup of a population. Genomic Offset can therefore be interpreted as a possible risk of maladaptation (Capblancq *et al*., 2020). Several studies have confirmed the correlation between Genomic Offset and average population fitness, using both computer simulations and common garden experiments (Gain *et al*., 2023; Láruson et al., 2020; Lotterhos, 2023; Rhoné et al., 2020; Fitzpatrick et al., 2021). However, Genomic Offset remains underutilized in invasion biology, despite its promise in predicting the success of invasive populations, as shown by a recent simulation study linking it to their probability of establishment (Camus *et al*., 2024). Moreover, the concept of Genomic Offset, which implies that populations adapt to new environments through preexisting variants, is consistent with the hypothesis of (genetic) pre-adaptation to invaded environments. Following this idea, calculating the Genomic Offset between the source and invaded environments can reveal which geographic areas have posed or would pose the greatest adaptive challenges to any source population.

Using the fruit fly *Drosophila suzukii* as a biological model, the present study will illustrate the contribution of genome scan and GEA approaches to understanding and predicting the adaptation of invasive species. Native to East Asia, *Drosophila suzukii* was first recorded in North America (California) and southern Europe (Spain and Italy) in 2008 (Asplen *et al*., 2015), and studies based on genetic data indicated independent invasion events on these continents (Fraimout *et al*., 2017; Adrion *et al*., 2014). In 2013, it had spread to South America (Deprá *et al*., 2014) and is now widespread in the Americas and Europe. It has also been detected in various parts of Africa, such as Algeria (Aouari *et al*., 2022), Morocco (Boughdad *et al*., 2021), Kenya (Kwadha *et al*., 2021), and most recently South Africa (IPCC, 2024). By laying eggs in ripening fruits, *D. suzukii* has caused significant damage to crops, particularly berry crops (e.g. cherries or strawberries) in the various areas where it has invaded, making it a major agricultural pest (Walsh *et al*., 2011).

The global spread of this species has been relatively well documented through multiple population genetic studies, which have consistently inferred similar invasion scenarios, at least in a general description(Adrion *et al*., 2014; Lewald *et al*., 2021; Feng *et al*., 2023; Fraimout *et al*., 2017; Gautier *et al*., 2022). Due to this convergence, the origin of invasive populations is now reasonably well established in *D. suzukii*. This clearly defined demographic background allows us to focus specifically on the adaptive dimension of the invasion process.

To date, few studies have comprehensively investigated the adaptation of *D. suzukii* to invaded environments. To our knowledge, only one study by Feng et al. (2024) examined *D. suzukii* environmental adaptation using GEA, identifying genes associated with environmental variables such as temperature and precipitation. Another study by Olazcuaga et al. (2020) examined the broad genetic basis of *D. suzukii* invasion success and found evidence for only a few genes associated with the invasive status of populations. Both studies were limited by the small number of sequences pooled population samples (Pool-Seq) and a fragmented genome assembly (Paris *et al*., 2020). As *D. suzukii* research expands, large-scale whole genome sequencing (WGS) efforts include sequenced individual samples (Ind-Seq; e.g. Lewald *et al*., 2021) in addition to the aforementioned Pool-Seq samples.

To deepen our understanding of the genetic basis of past *D. suzukii* invasion and to further identify geographic regions at risk of future invasion, we present a combined analysis of 27 Pool-Seq and 82 Ind-Seq samples, together representing a total of 37 populations. These data are primarily sourced from carefully curated publicly available datasets (Gautier, 2023), with the addition of newly sequenced samples from four previously unsampled areas. To enable efficient integration of these heterogeneous data for GEA and other genome scan approaches, we extend the Bayesian hierarchical models implemented in BayPass software (Gautier, 2015) for the joint analysis of hybrid data sets consisting of read counts from Pool-Seq samples and genotype likelihoods (GL) from Ind-Seq samples. In addition, to improve the precision of genomic analyzes, we rely on a newly developed chromosome-level assembly for the genome of *D. suzukii*, derived from an isofemale strain of Japanese origin (i.e., representative of the native range). Based on these new datasets, genomic resources, and methodological developments, we first search for key genes and biological processes that may have been involved in the adaptive process during invasion by extending the approach described in Olazcuaga et al. (2020). To further quantify the global contribution of genetics to *D. suzukii* invasion, we use Genomic Offset statistics based on a GEA with 29 covariables characterizing the environment of the 37 studied populations, to specifically address the three following questions: i) which parts of the invaded areas of Europe and North America may have posed the greatest adaptive challenge to *D. suzukii* ? ii) can we predict which parts of the geographic areas that have not yet been (fully) invaded (e.g. in South America and Africa) are at a higher risk of being invaded in the future? and iii) can we anticipate which region could host the highest-risk source population for a putative invasive *D. suzukii* population establishing in Australia where this pest is currently absent? The second and third applications aim at illustrating how Genomic Offset statistics may represent an additional potentially valuable tool to be considered in risk assessment and Integrated Pest Management (IPM) practice. By providing a comprehensive view of the adaptive history of *D. suzukii*, we seek to describe a generalizable and operational population genomics framework that is broadly applicable across species to study and predict biological invasions (Figure 1).

**Figure 1:**
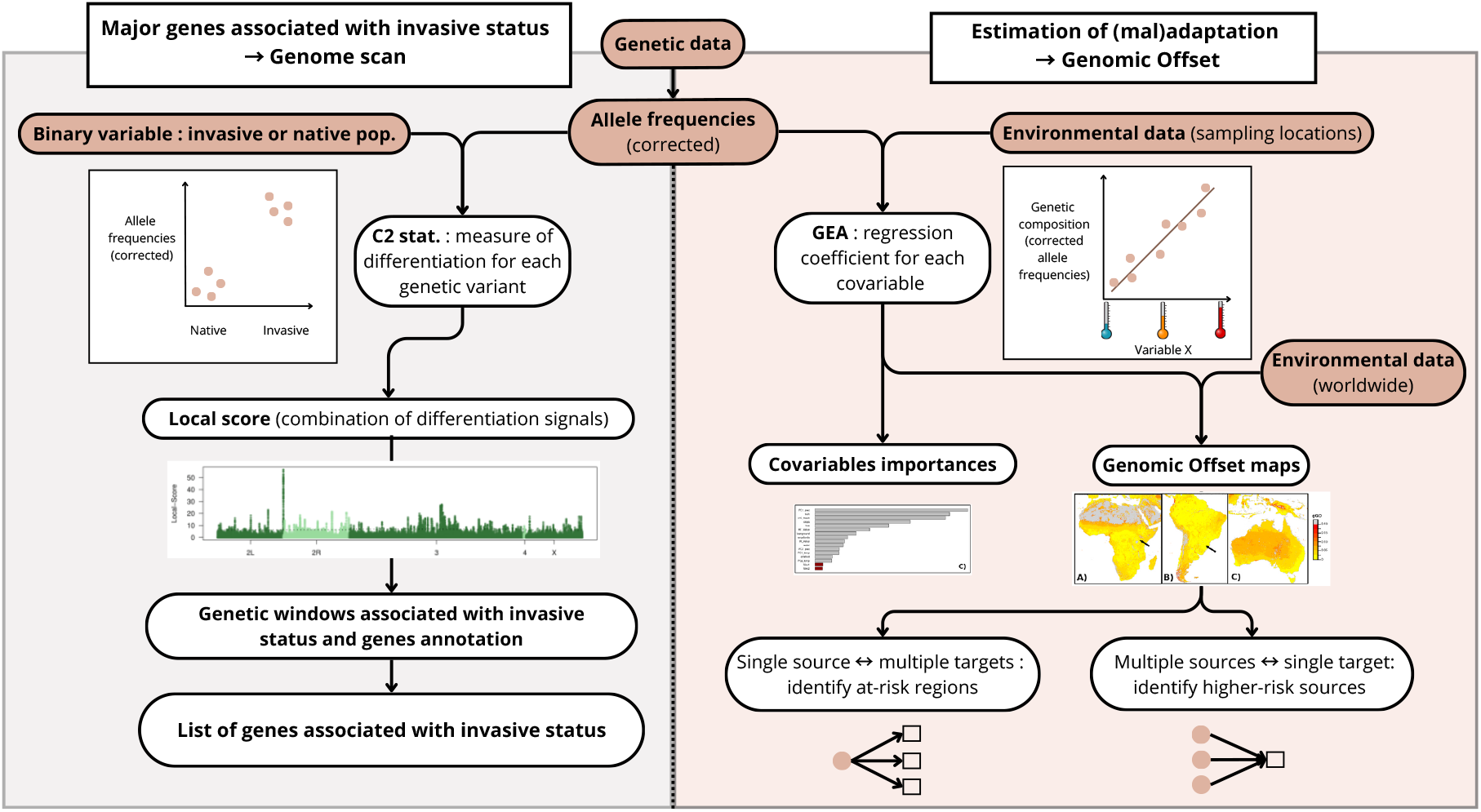
Conceptual graph illustrating the two complementary approaches used here to study adaptive challenges associated with invasion from a population genomics perspective. Genome Scan (left panel) is a relatively targeted approach that involves using a differentiation statistic (*C*_2_ here) for each variant based on a binary variable (native vs. invasive) in order to identify major genes associated with invasive status. Genomic Offset (right panel) is a broader approach based on GEA, which models the genome-wide relationship between adaptive genetic variation and environmental variables. This GEA model serves as a toolbox to assess the contribution of environmental variables to local adaptation and to compute Genomic Offset for various predictive applications. Together, these approaches thus provide both detailed insight into genes linked to invasiveness and integration of information on environmental adaptation to understand and predict invasion. Note that BayPass was used here for both approaches, but other software can also be applied within this framework.

## Materials and Methods

### Construction and annotation of a new *de novo* genome assembly

To date, all available reference assemblies for the *D. suzukii* genome were obtained from isofemale lines originating from the European (Ometto *et al*., 2013) or US (Chiu *et al*., 2013; Paris *et al*., 2020) invaded areas. They also remained suboptimal both in terms of contiguity and completeness (e.g., they only consisted of female assemblies). To circumvent these limitations, we generated PacBio HiFi and high-throughput chromosome conformation capture (HiC) sequencing data from individuals belonging to an inbred strain of Japanese origin (Ohashi *et al*., 1991) to construct a male chromosome-level assembly, hereafter referred to as *dsu_isojap1*.*0*. Briefly, the primary whole-genome assembly was generated with `hifiasm` (v0.19.8; Cheng *et al*., 2021), followed by contaminant removal with Kraken2 (v2.1.2; Wood *et al*., 2019) and self-alignment curation using `minimap2` (v2.20-r1061; Li, 2021). The final assembly was visually compared with those available for closely related species (e.g. *D. biarmipes*) using dot plots generated with `dgenies` (v1.5.0; Cabanettes and Klopp, 2018), and its general properties were summarized with snail plots using the `BlobToolKit` viewer (v4.3.5; Challis *et al*., 2020). These include the representation of contiguity and completeness which was evaluated with `BUSCO` on the `diptera odb10` dataset (v5.4.7; Manni *et al*., 2021). We finally annotated repetitive elements using `RepeatMasker` (v4.1.4; Smit *et al*., 2023) based on the curated and annotated TE database developed by (Mérel *et al*., 2021) for the *D. suzukii* genome. The assembly *dsu_isojap1*.*0* was then submitted to the NCBI repository for gene annotation with the NCBI Eukaryotic Genome Annotation Pipeline (Thibaud-Nissen *et al*., 2013). For more details on the data generation and assembly procedure, see Supplementary Note N1.

### *D. suzukii* population samples and whole genome sequencing data

To explore the genetic mechanisms underlying *D. suzukii* adaptation to invaded areas, we built a WGS dataset that covers a wide range of populations, in order to maximize geographic coverage and capture the species’ genetic diversity in its native and invasive range. WGS data representative of 32 worldwide population samples, derived from Pool-Seq (n = 24 pools; Table S1) or Ind-Seq (n=73 individuals representative of 8 populations; Table S2) were obtained from the NCBI SRA repository (https://www.ncbi.nlm.nih.gov/sra/). These data, sampled between 2014 and 2019, were the result of a careful curation of publicly available WGS data from three recent studies (Olazcuaga *et al*., 2020; Lewald *et al*., 2021; Feng *et al*., 2024), aimed at removing contaminated samples (Gautier, 2023) and excluding (or merging) geographically redundant samples (see Supplementary Note N2 for details). To further improve the worldwide representation of the samples, we newly sequenced four DNA pools representative of a single South American (AR-Gol from Argentina, n=100) and three European (CH-Del from Switzerland, n=58; DE-Jen from Germany, n=100; and RS-Zaj from Serbia, n=100) populations, as well as nine native individuals collected in Chengdu (China). These newly sequenced samples were collected between 2014 and 2023. Note that DE-Jen individuals are from the same sampling used in Olazcuaga et al. (2020) and Feng et al. (2024), but with all non-*D. suzukii* individuals discarded (Gautier, 2023). Briefly, for sequencing these new samples, we extracted DNA from whole bodies (or pools of thoraxes) using the DNeasy Blood and Tissue Kit (Qiagen, USA) according to the manufacturer’s recommendations and constructed libraries using the NEBNext Ultra II DNA Library prep kit for Illumina (New England Biolabs). These were paired-end (PE) sequenced (2×150 bp or 2×250 bp) on a NovaSeq6000 Illumina sequencer (Tables S1 and S2) on the Get Plage platform (INRAE, Toulouse).

### WGS data processing, SNP calling and genotyping

WGS data were first filtered using `fastp` (v0.23.2; Chen *et al*., 2018) run with default options to remove contaminating adapter sequences and trim poor quality bases (i.e., Q*<*15). Read pairs with either one read containing more than 40% low-quality bases or more than 5 N bases were removed. Filtered reads were then mapped onto the new *dsu_isojap1*.*0* assembly, complemented with the complete mitochondrial sequence of *D. suzukii* (GenBank Accession ID: *KU588141*) and the complete sequence for the Wolbachia endosymbiont wRI strain of *D. simulans* (NCBI ID: *GCA_000022285*), which is closely related to that of *D. suzukii* (Turelli *et al*., 2018), using `bwa-mem2` with default options (v2.2.1; Vasimuddin *et al*., 2019). PCR duplicates were then removed using the `view` and `markdup` programs of `SAMtools` (v1.14; Li *et al*., 2009). The mean realized coverage was estimated using `mosdepth` (v0.3.6; Pedersen and Quinlan, 2017) with the option *-Q 20* to ignore reads with low mapping quality. As detailed in Tables S1 and S2, on autosomes (i.e. scaffolds 2L, 2R, 3 and 4) the coverages varied from 42.94 (US-CAOx) to 105.6 (JP-Kan) with a median of 61.3 for Pool-Seq data; and from 1.47 to 13.7 with a median of 6.98 for Ind-Seq data.

Variant calling was performed using the Haplotype Caller implemented in `FreeBayes` (v1.3.6; Garrison and Marth, 2012), jointly considering all 27 Pool-Seq and 82 Ind-Seq alignment files from *D. suzukii* samples. As detailed in Supplementary Note N3, variants were called with stringent quality and depth filters per 250kb window across the genome and filtered to retain only high-quality bi-allelic variants, yielding 6,814,770 autosomal bi-allelic variants consisting of 6,732,348 SNPs, 71 MNPs (Multi-Nucleotide Polymorphisms), 52,124 indels and 8,239 so-called complex variants. Like-wise, 1,226,417 X-linked bi-allelic variants were obtained, including 1,209,454 SNPs, 22 MNPs, 15,343 indels and 1,598 complex variants. To provide a global visualization of the genetic structuring of the populations, we performed a random allele PCA using the `randomallele.pca` function of the R package `poolfstat` (v2.2.0 Gautier *et al*., 2022) on the entire *vcf* combining Pool-Seq and Ind-Seq data parsed with the `vcf2pooldata` function.

### Genome scan analyses

To identify genomic regions and biological processes potentially associated with invasion-related adaptation, we performed genome-wide scans of differentiation between native and invasive populations, following the approach of Olazcuaga et al. (2020). We used the *C*_2_ statistic (Olazcuaga *et al*., 2020), which identifies overly differentiated variants between two population groups while accounting for population structure. As *C*_2_ is non-polarized (i.e., it does not indicate in which grouop selection occurred) we use the term “associated with invasive status” throughout the manuscript to reflect association with being invasive or not. All genome-wide scans were performed using an upgraded version (v3.0) of BayPass (Gautier, 2015; Olazcuaga *et al*., 2020). As detailed in Supplementary Note N4, this new version extends all the previously described models to the analysis of individual genotyping data (coded as genotype likelihood, i.e. GL), which allows one to properly deal with low-to mid-coverage Ind-Seq data (e.g., *<*10-15X) by integrating over genotype uncertainty. In addition, it now allows the analysis of hybrid data consisting of sample allele and/or read counts and/or GL data under all the models described previously (Gautier, 2015; Olazcuaga *et al*., 2020). Following the BayPass manual recommendation (see also Gautier *et al*., 2018), the complete dataset was first split into subdata sets of 68,150 markers and 61,320 markers for autosomes and X-chromosome, respectively. Each was analyzed with the default option to estimate the scaled covariance matrix of population allelic frequencies (Ω), accounting for the neutral population structure, and three *C*_2_ statistics (Olazcuaga *et al*., 2020) contrasting native versus i) all worldwide (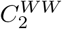, including both European and American populations); ii) European 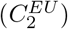; or iii) American 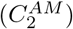 invasive populations. As observed in previous similar analyses (e.g., Gautier *et al*., 2018; Olazcuaga et al., 2020), posterior mean estimates of Ω entries over subdata sets (from the same chromosome type) were highly consistent. We thus only considered those obtained from a single randomly chosen subdata set for representation and further analysis under the `covmcmc` mode (see Estimation of the geometric Genomic Offset (gGO)). Likewise, *C*_2_ estimates were stable across three independent analyses (with different seeds) of each subdata set, and only those from the first run (`-seed 5001`) were reported. To identify and delineate significant genomic regions associated with invasive status based on the three different contrasts, i.e. with an excess of high *C*_2_ values, we relied on the local-score approach described in Fariello et al. (2017) and (re)implemented in the R function `compute.local.scores` available in the BayPass software package (v3.0). Individual SNP scores were computed as − log 10(*p*)− *ξ*, where *p* is the p-value associated with a given *C*_2_ and *ξ* is a threshold parameter (Fariello *et al*., 2017) that we set to *ξ* = 2 (i.e., *p <* 10^−2^ were cumulated and others were scored 0 in the underlying Lindley process). As recommended in the BayPass manual, to avoid penalizing the approach, we discarded low-polymorphic SNPs (with an estimated overall allele frequency 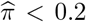 or 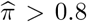) from the local–score calculation.

For each significant window, we retrieved the nearest annotated gene within 5 kb of the *C*_2_ peak, and determined *D. melanogaster* orthologs using the `datasets` NCBI command-line tool (v16.40.0). Gene Ontology (GO) enrichment analysis for Biological Process was performed on the resulting list of candidate genes (from all three *C*_2_) using the R package *clusterProfiler* (v4.12.0; Wu *et al*., 2021) with default settings. GO terms were obtained from the `org.Dm.eg.db` annotation R package.

### Estimation of the geometric Genomic Offset (gGO)

To globally assess the adaptive challenges faced by *D. suzukii* during its invasion and to anticipate its future spread, we used a GEA framework to compute geometric Genomic Offset (gGO; Gain *et al*., 2023)) based on 29 environmental covariables across its current range. We first performed a GEA under the BayPass standard covariate model (Gautier, 2015) with option `-covmcmc`, using the matrix Ω estimated above, to estimate the effect (regression coefficients) of environmental covariables on the (scaled) population allele frequencies. This allows modeling the relationship between the genomic diversity and environmental variation of (all) the studied populations. We used 29 environmental covariables that can be grouped into three data types: bioclimatic, land use, and agricultural production (see Supplementary Note N6 for more details). To address variable interdependence, which can impair Genomic Offset’s predictive performance (Camus *et al*., 2024), we performed a PCA on precipitation variables and another one on temperature variables (both bioclimatic data), and retained 2 PCs (explaining *>* 75% of the variance) for each group of variables as environmental covariables in the GEA analysis. After PCA transformation of temperature and precipitation variables, the final dataset comprised 14 environmental covariables. We used the regression coefficients estimated with BayPass for both autosomes and X-chromosome and computed the gGO using the `compute_genetic_offset` function of BayPass v3.0 (Camus *et al*., 2024; Gautier *et al*., 2024) without any pre-selection of the variants, as recommended by recent studies (Gain *et al*., 2023; Lind and Lotterhos, 2024; Camus *et al*., 2024). In accordance with the interpretation of gGO as an environmental distance of the covariables considered weighted by their contribution to the local adaptation, we also evaluated the importance of each environmental covariable in gGO computation (Gain *et al*., 2023). After performing a spectral decomposition of the covariance matrix of the estimated regression coefficients, the importance *γ*_*i*_ of each covariable *i* is then defined as 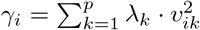, where *λ*_*k*_ is the *k*th eigenvalue and *v*_*ik*_ is the *i*th coordinate of its associated eigenvector. To empirically establish a baseline threshold for key variables, we included two simulated ‘fake’ covariables in the analysis — random variables drawn from a normal distribution and constrained to exhibit correlation coefficients between –0.1 and 0.1 with the 14 “real” variables. Finally, to produce spatial representations of genomic offset (relative to a given source environment) we computed gGO values at the scale of each grid cell using environmental rasters. Each cell was characterized by the 14 covariables retained in the GEA. Bioclimatic and land-use layers were available at 30 arc-seconds resolution (∼1 km^2^ at the equator), while agricultural layers were at 5 arc-minutes, (∼ 55 km^2^ at the equator). Following an approach similar to Lachmuth *et al*. (2023), we scaled all gGO maps by the highest gGO among the 37 studied populations (observed for the CN-Nin/US-Haw sample pair), i.e., the maximum gGO value of established populations in the current worldwide *D. suzukii* range, which was equal to 0.19.

We then defined three classes of gGO values based on the distribution observed for the 666 pairs of 37 sampled populations (Figure S4) corresponding to i) “low gGO” (0–0.05) reflecting the range of gGO values observed within the native area, where all populations are expected to be well adapted to their local environments due to their long-term presence (this range includes all native–native comparisons but some outliers related to CN–Nin population); ii) “moderate gGO” (0.05–0.15) which corresponds to the outliers gGO values found among native–native comparisons, representing increasing levels of environnemental differentiation,and thus potential local maladaptation; and iii) “high gGO” (0.15–0.19) which includes the most divergent values, observed primarily in comparisons involving at least one invasive population, and potentially reflecting stronger shifts in environmental conditions. Values above 0.19 were considered extreme.

To evaluate the strongest adaptive challenges faced by *D. suzukii* during its invasion, we computed gGO maps over Europe and North America by calculating, for each grid cell, the gGO relative to a reference environment representative of the source populations. For Europe, CN-Lia was used as the reference, reflecting a single-source invasion (Fraimout *et al*., 2017). For North America, invasion scenarios involved genetic admixture from Asia and Hawaii (Fraimout *et al*., 2017; Gautier *et al*., 2022). In this case, the interpretation of gGO as an environmental distance weighted by the adaptive genome composition was used to generate a combined (i.e. admixed) gGO map. The combined map was created using both CN-Nin and US-Haw as reference environments, with CN-Nin and US-Haw assigned a 75% and 25% weights, respectively, in agreement with previous estimated genetic contributions in North American populations (Fraimout *et al*., 2017).

## Results

### A new male chromosome-level genome assembly

The combination of PacBio HiFi long reads and HiC sequencing data allowed the generation of a highly contiguous assembly consisting of 54 scaffolds totaling 282,295,391 bp (N50=32.7 Mb) and including 99.06% (98.48% complete and single) of the 3,285 BUSCO `diptera odb10` dataset genes (Manni *et al*., 2021, see Table 1 and Figure S1A).

**Table 1:**
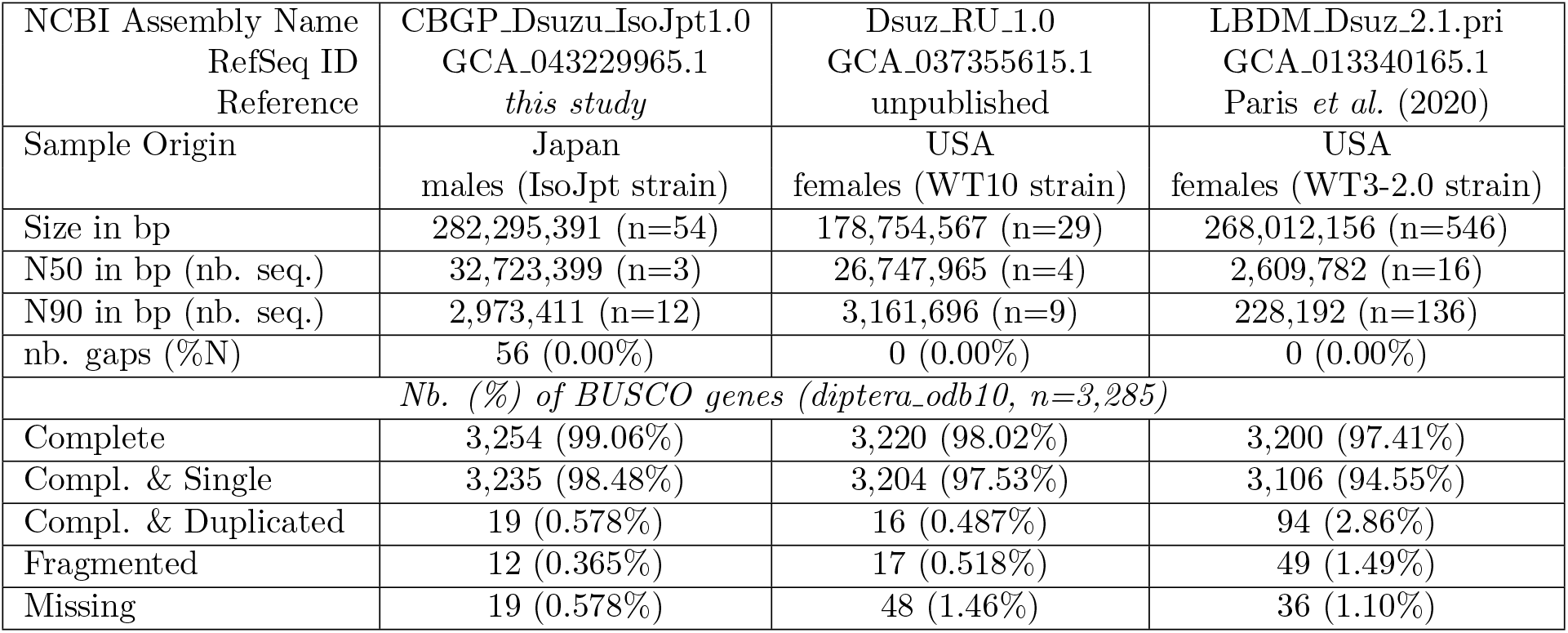
Descriptive statistics of the new *dsu_isojap1*.*0* assembly for the *D. suzukii* genome and the two other (partial) chromosome-level assemblies available from the NCBI repository as of March 2025. Aligning *dsu_isojap1*.*0* with the `RU 1.0` assembly (Figure S2) allowed the identification of 40 sequence similarity blocks *>* 1 Mb totaling 171.5 Mb (i.e. 96.1% of the `RU 1.0` assembly) with only 1.33% nucleotide divergence. Likewise, compared to the more fragmented `WT3-2.0` assembly, 35 blocks of sequence similarity *>* 1 Mb and totaling 153.6 Mb were found with a nucleotide divergence of 0.813%.

In order to validate the structure and quality of the new assembly, we compared it to genome assemblies for other closely related Drosophilids species. The alignment of the *dsu_isojap1*.*0* assembly with `Dbia2.1`, the chromosome-level reference assembly of the closely related species *Drosophila biarmipes* (Figure S1B), showed that six large scaffolds completely covered the five *D. biarmipes* autosome arms (2L, 2R, 3L, 3R and 4) and the X chromosome. Then, these *dsu_isojap1*.*0* scaffolds were named following the nomenclature of *D. biarmipes* (and *D. melanogaster*), as also justified by cytogenetic studies (Drosopoulou *et al*., 2019). Likewise, as detailed in Table S3 and Supplementary Note N1, a total of 3,197 of the 3,285 BUSCO genes from the `diptera odb10` dataset were found complete and in single copy in both the *dsu_isojap1*.*0* and the *Dmel6* assembly for the *D. melanogaster* genome (Hoskins *et al*., 2015), revealing near-complete gene overlap (*>*95.8%) across most chromosomes between these two assemblies, which aligned with the very close but not perfect conservation of Muller elements among *Drosophila* species (Bhutkar *et al*., 2008).

In agreement with previous studies on the WT3 v2.0 assembly (Paris *et al*., 2020; Mérel et al., 2021, e.g.), we found that *dsu_isojap1*.*0* was characterized by a very high TE content (53.24% of the assembly). The count distribution of the different TE orders over the scaffolds highlighted distinct chromosomal regions with some clear TE signature specifying centromeric regions similar to what was observed in *D. melanogaster* genome by Chang et al. (2019), although different TE superfamilies are involved (Figure S1C). Finally, the gene annotation pipeline identified a total of 17,936 genes (including 15,749 coding and 2,219 noncoding) and 266 nontranscribed pseudogenes. Most annotated genes were mapped to the identified autosomal scaffolds, and along the main chromosomes, the gene density was lower in the presumed centromeric regions as in the middle of chr3 (Figure S3). Overall, our new annotated chromosome-level assembly (*dsu_isojap1*.*0*) represents a clear improvement over previous assemblies (Table 1), exhibiting substantially higher contiguity and completeness, with 99.06% of BUSCO genes complete (compared to 98.02% and 97.41% in the two earlier assemblies) and fewer fragmented or missing BUSCOs. It thus provides a more consistent and comprehensive resource for the population genomics investigations conducted in the present study, as well as for future population genomics research.

### A combined Ind-Seq and Pool-Seq curated dataset representing the *D. suzukii* worldwide genetic diversity

We verified that the genetic structure of our 37 populations aligns with the known demography of *D. suzukii* and confirmed that combining Ind-Seq and Pool-Seq data did not create inconsistencies. To do so, we performed a random allele PCA focusing on autosomal variants. As shown in Figure 2B and in agreement with previous population genetics studies (e.g. Fraimout *et al*., 2017; Lewald *et al*., 2021), the first factorial plan grouped the populations into four main clusters: i) the native area; ii) European invaded area; iii) populations from the Northwest US and Hawaii; and iv) all other populations from the American invaded area (including the newly generated AR-Gol from Argentina).

**Figure 2:**
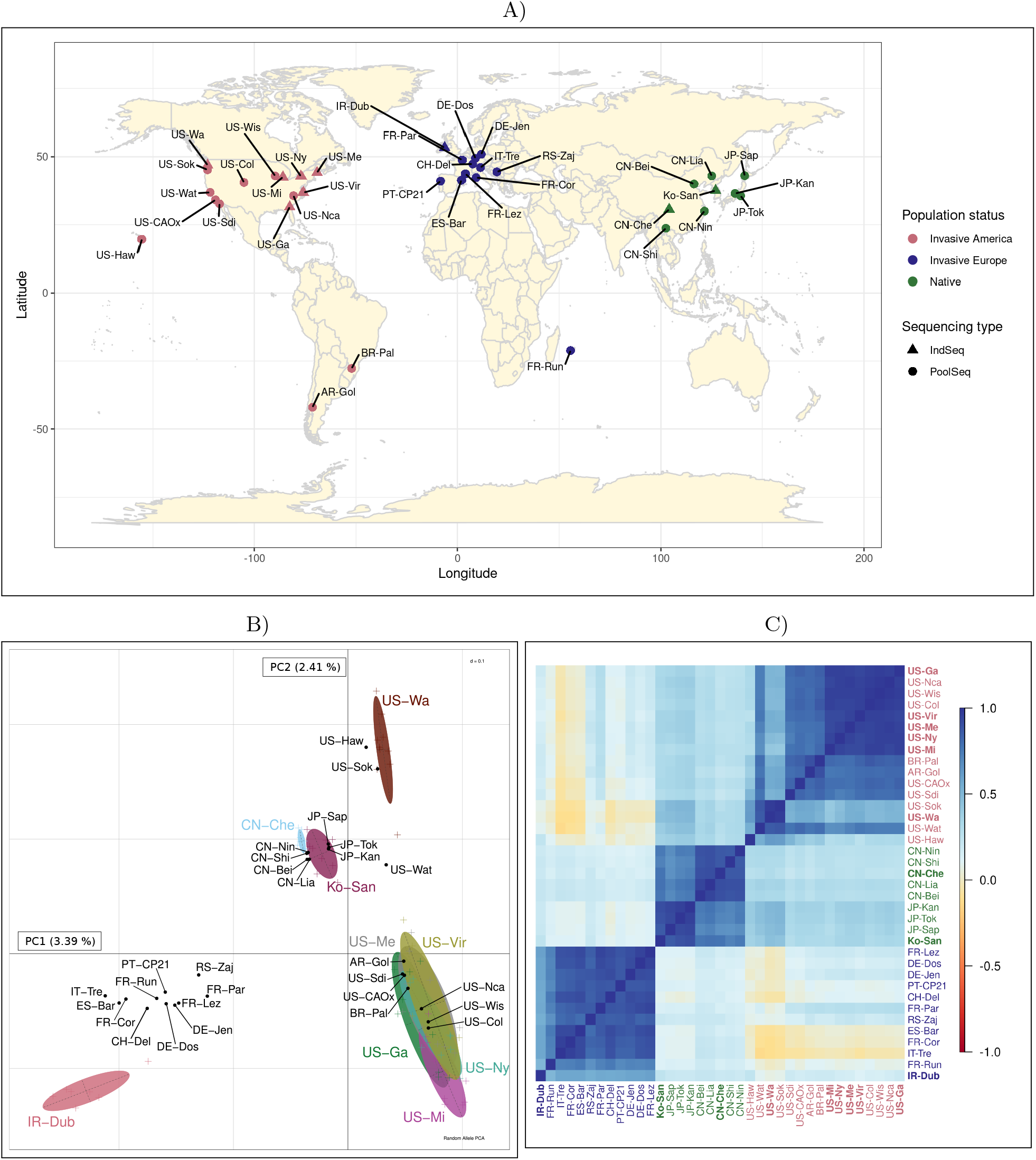
Description of the 37 studied *D. suzukii* population samples and structuring of genetic diversity on autosomal variants. A) Geographical location of the 37 *D. suzukii* populations. Populations from the native area are displayed in green, while invasive American and European populations are displayed in red and blue, respectively. B) Random allele PCA representation of the population samples. Pool-Seq samples are represented by black dots, while Ind-Seq samples are indicated by crosses colored according to their population of origin (with 95% ellipses summarizing the overall dispersion of the underlying individuals around their population center of gravity). C) Correlation plot of the scaled covariance matrix Ω estimated from the combined analysis of Pool-Seq and Ind-Seq data with the new version of BayPass software (v3.0; Gautier, 2015) on autosomal variants. The name of the populations represented by Ind-Seq samples are highlighted in bold.

The positioning of the populations of the invaded European area did not clearly reflect geographical or temporal origins, suggesting substantial migration events across Europe as well as heterogeneous introgression events of European populations by propagules from the North American continent (Fraimout *et al*., 2017). A notable exception is the IR-Dub sample (Ind-Seq), which appears clearly separated from other European populations (all Pool-Seq) in the PCA. This separation is unlikely due to sequencing type, as Lewald et al. (2021) observed a similar pattern using only Ind-Seq data. The genetic distinctiveness of IR-Dub may be explained by: i) its sampling shortly after the species’ first record in Ireland, ii) geographic isolation limiting gene flow, and iii) stronger admixture signals from Eastern US or Brazilian sources compared to other European populations (Lewald *et al*., 2021). For the American invaded area, differentiation between eastern and northwestern populations matched previous findings (Lewald *et al*., 2021), and as expected showed no apparent effect of sequencing type on population clustering.

The correlation plot of the Ω matrix estimated with BayPass showed a similar structuring into four groups (Figure 2C). The Ω representation did not cluster samples by sequencing type (Pool-Seq or Ind-Seq), indicating that the statistical models implemented in the new BayPass version perform as expected on hybrid datasets. Both the Ω matrix and random allele PCA estimations were independently repeated with X–linked variants, yielding results similar to autosomal variants (Figure S5), with only minor differences (see Figure S5 legend).

### Genome scan for association with invasive status

To identify genomic regions potentially involved in adaptation to the invaded environments, we searched for clusters of overly differentiated variants between native and invasive populations using a local-score approach based on the *C*_2_ statistic. Cumulating association signal of markers with a *C*_2_ p-value below 0.01, we could identify 231 significant windows for 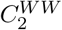, 208 windows for 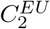, and 18 windows for 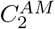 (Figure 3), predominantly located on chromosomes 3 (n=195) and 2R (n=108), followed by the X chromosome (n=74), chromosome 2L (n=69), and chromosome 4 (n=11) (see Tables S4 to S8 for the details of all the windows for each chromosome).

**Figure 3:**
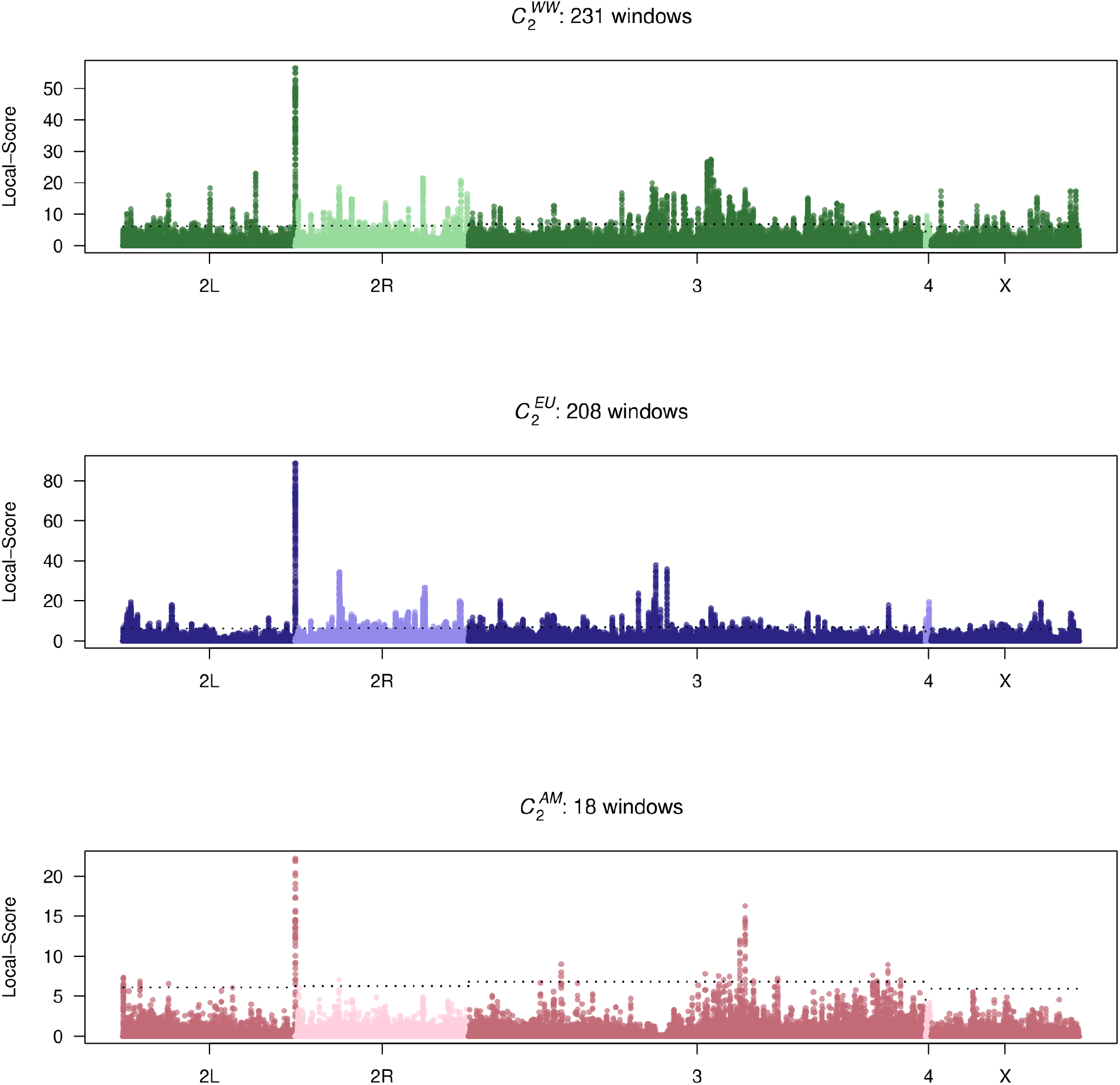
Manhattan plot of the *C*_2_ p-values based local-score contrasting native versus A) all worldwide 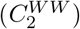; B) European 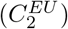; and or C) American 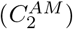 invasive populations. The dashed lines indicate the chromosome-specific 1% significance threshold for the local-score. The number of significant local-score windows is given in the main legend of each plot. Details about all the windows (separated by chromosome) can be found in Tables S4 to S8, and the top 45 genes associated with the highest local-score values are presented in Table 2.

The size of the identified windows ranged from 155 bp to 476,822 bp and included 7 to 202 SNPs (Tables S4 to S8). Merging the windows that overlapped between different *C*_2_ tests resulted in a final set of 457 candidate regions: 171 specific to 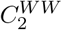, 154 to 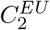, and 10 to 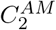. The windows detected only with 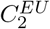 or 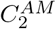 likely reflect continent-specific signatures — otherwise they would also have been detected by the 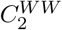 test, because of its larger sample size being expected to increase power. Conversely, our ability to assess convergence between 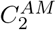 and 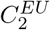 is limited by their lower power, particularly for 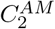, as both detected fewer significant regions than 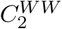. Hence, the 171 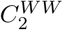 specific regions might result from a combination of signals that are consistent between continents but too weak to be detected in any of them. Overall, 52 windows were shared by 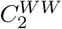 and 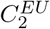; six by 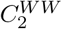 and 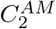; and only two by the three contrasts, which remains higher than the number (n = 0.54) expected if all tests were completely independent (see Supplementary Note N5).

**Table 2:**
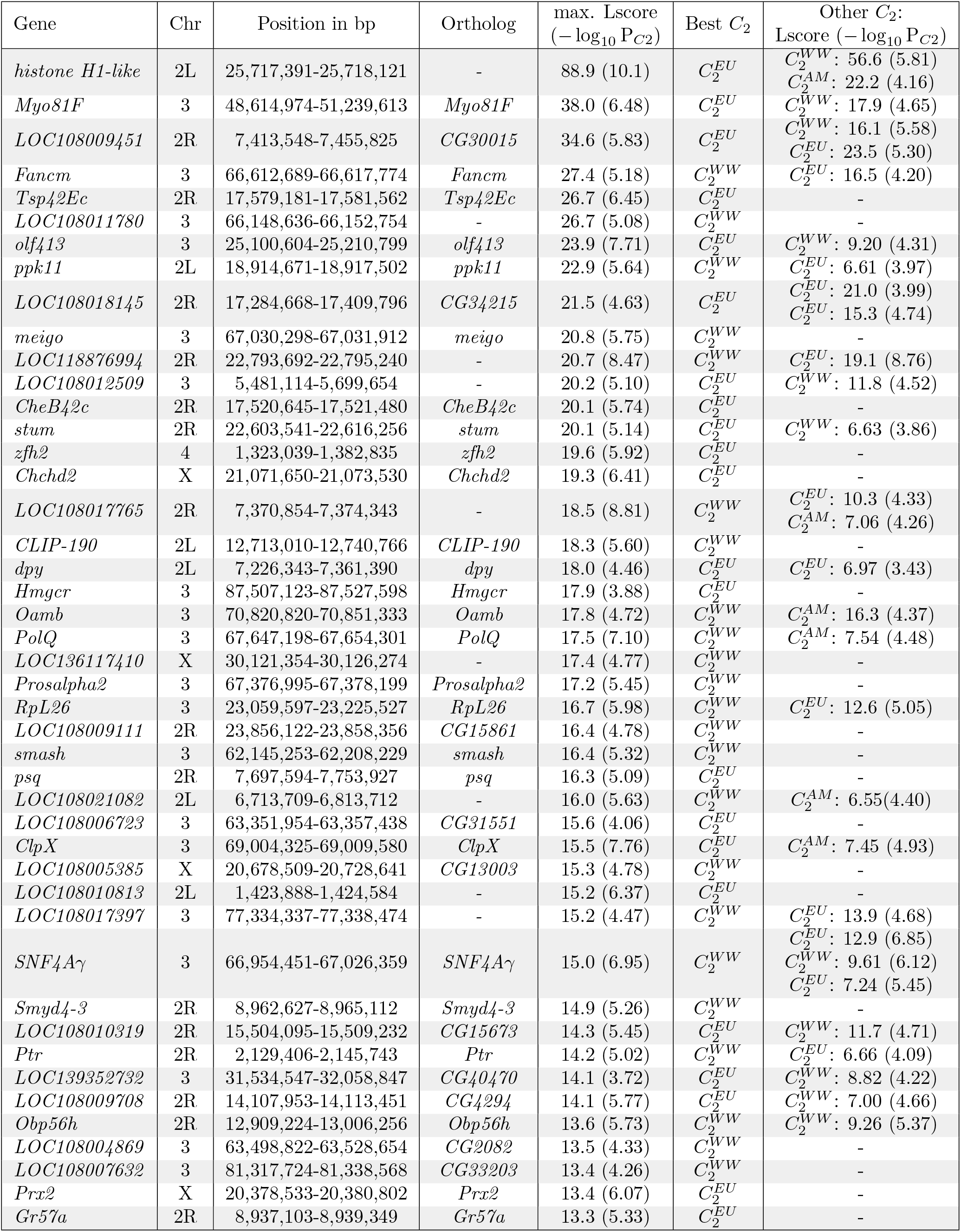
Description of the top 45 *D. suzukii* windows with the highest *C*_2_ local-score among the 396 (out of 457) with annotated positional candidate genes. The table gives for each gene i) its position in the new *dsu_isojap1*.*0* assembly of the *D. suzukii* genome; ii) its corresponding *D. melanogaster* ortholog; (iii) the highest maximum Lindley score (Lscore) and the most significant *C*_2_ P-values (in − log_10_ scale) for the associated window; iv) the *C*_2_ statistic corresponding to this highest Lscore; and v) other *C*_2_ statistics with which the gene was associated, along with the maximum Lscore and most significant *C*_2_ P-values (in − log_10_ scale) for the corresponding window.

Gene annotation of the 457 significant windows highlighted 396 positional candidate genes among which 322 could be associated with a *D. melanogaster* ortholog (Tables S4 to S8). The GO enrichment analysis for Biological Process (for all candidate genes with orthologs) showed several terms that were significantly enriched in our candidate genes dataset compared to the rest of the genome (Figure S7). The GO terms with the most significant enrichment were “cell adhesion”, “epithelial morphogenesis”, and “dicarboxylic acid transport”. Further down the list, significant terms were mostly related to morphogenesis but also included cognition, chemosensory and olfactory behavior, and memory.

More specifically, the window at the 2L centromeric end associated with the *Histone H1-like* candidate gene clearly stood out, since it was among the two windows shared by the three contrasts and displayed the highest local-score in each test. The three corresponding windows were also unusually large (476,822 bp, 440,361 bp and 356,304 bp for 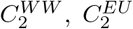 and 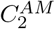 respectively), although inflated by an extended region of extremely low SNP density (4 SNPs spanning ca. 300 Mb) beyond the local-score peak (Figure S8). Nevertheless, the local-score peak was located approximately just 1kb upstream of the *Histone H1-like* gene, confirming a strong signal associated with it. Histones play a crucial role in regulating chromatin structure and gene expression, and Histone H1, the least conserved histone, is essential for heterochromatin silencing and genome stability in *D. melanogaster* (Bayona-Feliu *et al*., 2016). Although its study has gained traction recently, its full range of functions remains unclear and is likely to include multiple variants with distinct functions (Prendergast and Reinberg, 2021). It should also be noted that we here identified an “H1-like” gene, rather than a canonical Histone H1, which may thus deserve further confirmatory functional characterization in *D. suzukii*.

Among our 45 top candidates (mainly related to 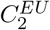 and 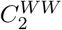, Table 2), we also identified a diverse set of genes related to key physiological and behavioral traits, including stress resistance and detoxification, dispersal and movement, chemosensation and feeding behavior, as well as immune response. These genes are involved in biological functions that may have contributed, to varying degrees, to invasive success (see Supplementary Note N7 for more details).

### Insights into past invasions from gGO

To determine whether some parts of the invaded areas of Europe and North America may have posed adaptive challenges to *D. suzukii* during its invasion, we computed gGO maps over these two continents by estimating, for each grid cell, the gGO values relative to a reference environment representative of the source populations. The gGO map over Europe (Figure 4A) generally displayed low to moderate gGO values, except in a few areas, such as the Alps and the cold regions of north-west Scotland and western Norway, with high or extreme values. The estimated gGOs were slightly higher overall across North America (Figure 4B) than in Europe, with uniformly moderate values across most of the continent. Extreme values were found mainly in the northern cold regions and the Sonoran Desert. It is worth noting that processing a combined map to reflect the admixed origin of North American invasive flies, with a weight of 75% and 25% for CN-Nin and US-Haw respectively, may have had a substantial impact on results. In agreement with this, we found that the gGO maps using CN-Nin only as environment of origin or US-Haw only (Figure S9) differed substantially from the combined map in Figure 4B, as US-Haw map displayed in particular lower values in Central America and western USA.

**Figure 4:**
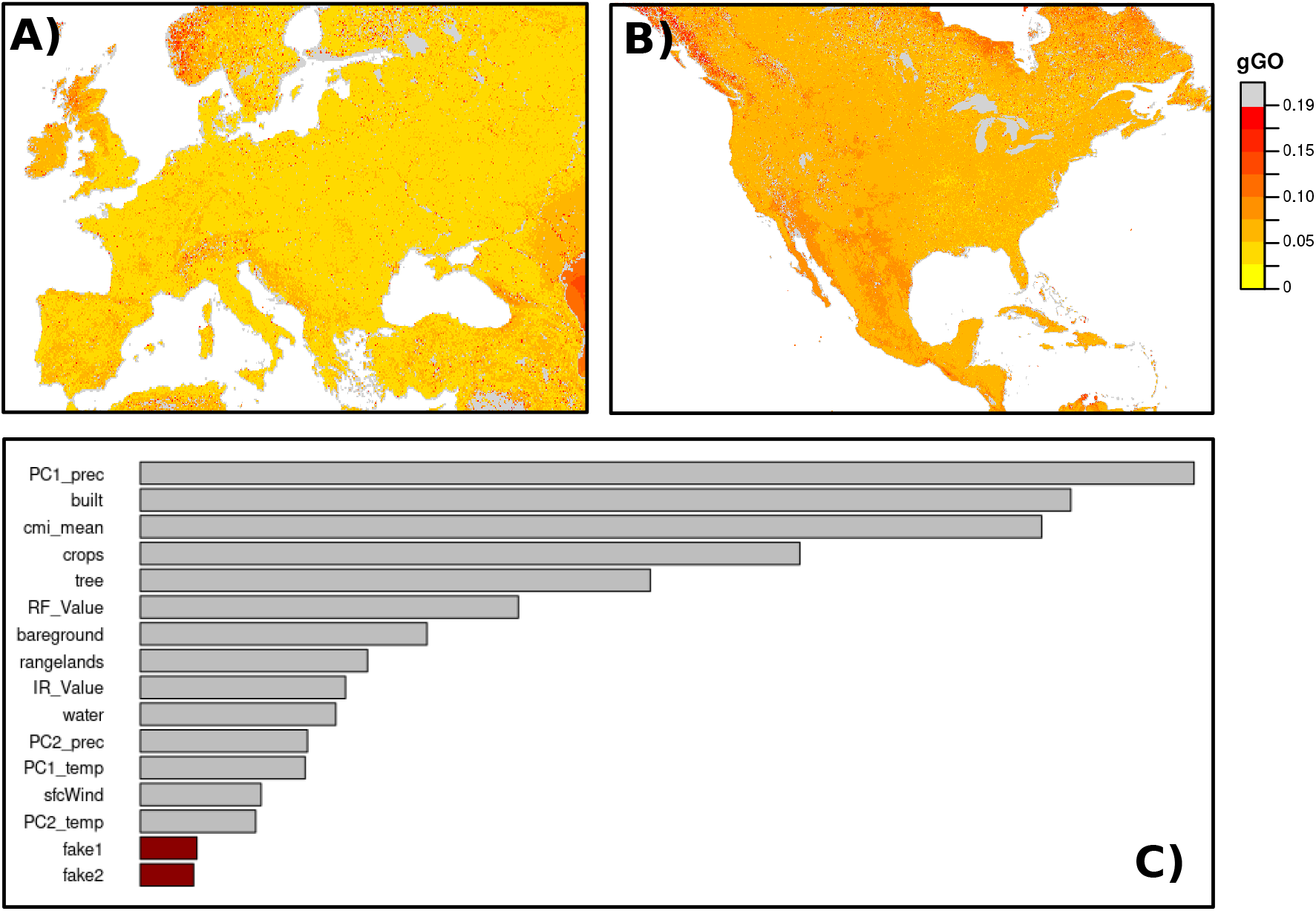
gGO values over European (A) and North American (B) invaded areas. For Europe, the reference environment corresponded to the one of the CN-Lia population, inferred as the closest proxy (Fraimout *et al*., 2017). For North America, to account for the admixed origin of the North America invasive populations (Fraimout *et al*., 2017; Gautier et al., 2022), we computed weighted gGO values by combining the two gGO maps (Figure S9) based on the environments of CN-Nin and US-Haw as references. The combined map used weights of 75% for CN-Nin and 25% for US-Haw, corresponding to their respective estimated admixture proportions in the primo-invading US population (Fraimout *et al*., 2017). Lowest (highest) gGO values correspond to lowest (strongest) adaptive challenges. Gray pixels represent outlying gGO values (as defined in Materials and Methods). (C) Ranking of the importance of the environmental covariables used for the gGO analysis. The two “fake” (control) variables are shown in red. PC1 prec and PC2 prec represent the two principal components for precipitation, while PC1 temp and PC2 temp correspond to those for temperature. Built, crops, tree, bareground, rangelands, and water refer to land-use classes. IR value and RF value indicate the annual harvested area for irrigated and rainfed perennial crops, respectively. Note that this variable importance ranking is presented here but also apply to the following gGO figures, which are based on the same regression coefficients.

The ranking of the environmental covariables confirmed the relevance of the predictors used in the gGO calculation (Figure 4C). As expected, the two control ‘fake’ covariables were of least importance, indicating that all other covariables included in the model were fairly informative with regard to local adaptation. Precipitation appeared as the most important factor, followed by built-up land use and mean monthly climate moisture (cmi mean, third). Agricultural and land-use covariables also played a prominent role, with crops, tree cover and annual harvested area for rainfed crops ranking fourth, fifth, and sixth respectively.

### Predicting future invasions from gGO

Following an approach similar to that above, we relied on gGO to predict which uninvaded geographical regions in Africa, South America, and Australia may pose stronger or weaker adaptive challenges for *D. suzukii* in the future, by deriving gGO maps over these three continents with respect to a given source environment. In this context, a lower gGO indicates a potentially more favorable adaptive landscape for establishment, whereas a higher gGO suggests a greater adaptive barrier to invasion. For Africa and South America, we selected source environments from representative sites where *D. suzukii* populations have recently established, and thus represent potential sources capable of spreading across these continents. These correspond to Longonot Farm (Nakuru county, Kenya; Kwadha *et al*., 2021) and Nova Veneza (Santa Catarina, Brazil; Deprá *et al*., 2014). For Australia, where *D. suzukii* is not established, we selected the environment of our US-Wa sample as a putative source reference, based on fresh fruit trade statistics (International Trade Center, 2024).

The gGO map of Africa displayed contrasting patterns (Figure 5A), with arid regions (e.g., the Sahara and Namib deserts) predominantly showing extreme or high values at their borders, and other regions showing moderate values (e.g., sub-Saharian area). In contrast, some parts of West Africa (e.g., Ivory Coast), Central Africa (e.g., Democratic Republic of the Congo), and East Africa (e.g., the Great Lakes region) displayed low gGO values, i.e., higher invasion risk. Interestingly, the very northern part of the continent, especially in northwestern Morocco, where *D. suzukii* infestation has recently been reported in several berry farms (Boughdad *et al*., 2021), was among the regions with the lowest values. The north-eastern region of Algeria, including the pomegranate agroecosystem where *D. suzukii* has been detected (Aouari *et al*., 2022), also showed low gGO values, but with a more heterogeneous pattern. In South America, most of the subcontinent showed low gGO values. As expected, the high or extreme outlying values were mainly located in regions with extreme climates, such as the Andes mountains or the west coast and southern parts of Patagonia (Figure 5B). It should be noted that the Chubut region in central Patagonia (Argentina) where *D. suzukii* has only recently been detected and considered established, and from which the AR-Gol sample originates, showed moderate gGO values. Finally, the gGO map of Australia displayed a moderate variability (Figure 5C). The highest gGO values were located in arid areas (i.e Simpson Desert), while the lowest gGO values were concentrated in the southern regions, particularly along the South East Coast, and in Tasmania.

**Figure 5:**
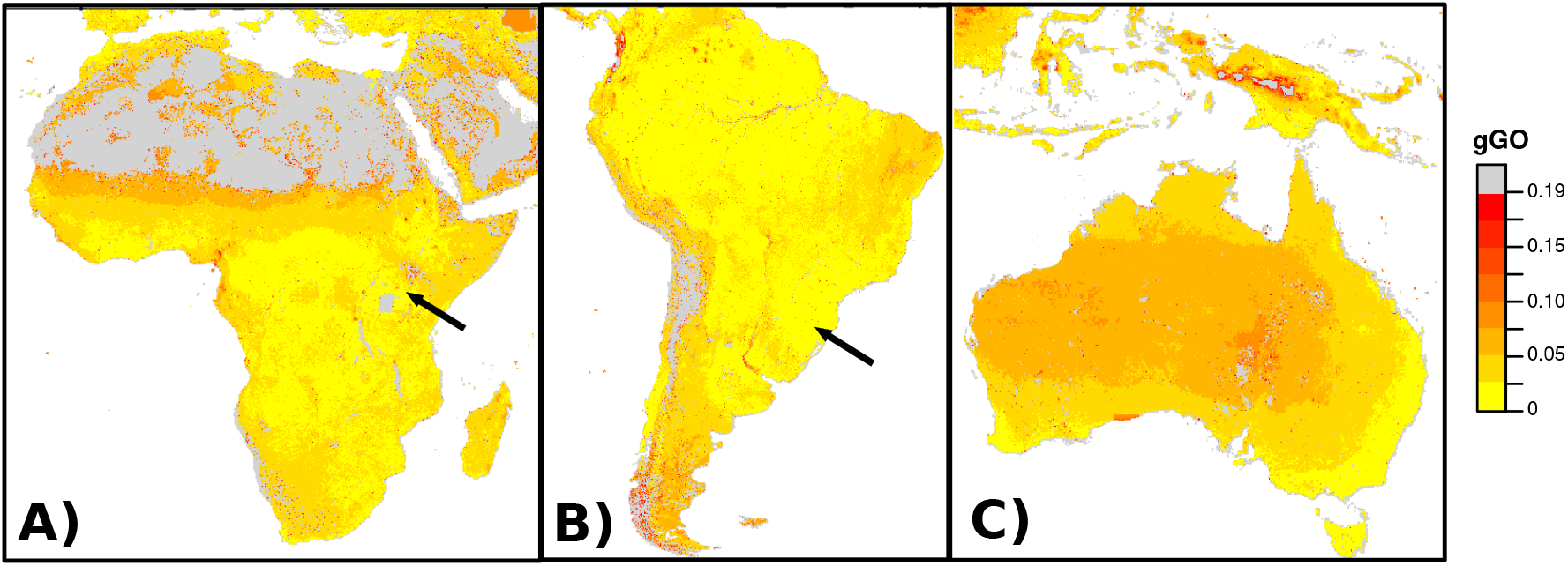
gGO estimates for the geographical regions in Africa (A), South America (B) and Australia (C) that have not been (fully) invaded to date. For Africa and South America, the black arrow indicates the (chosen) source reference environment where *D. suzukii* was first recorded and recently described as established in these continents (i.e. the environment of Longonot Farm - Nakuru county, Kenya - for Africa and of Nova Veneza - Santa Catarina, Brazil - for South America). For Australia, where *D. suzukii* has not yet been found as established, the reference environment associated with our Northwestern American sample US-Wa was used as a (putative) source reference based on trade statistics for fresh fruit exchanges (International Trade Center, 2024). Gray pixels represent outlying gGO values (as defined in Materials and Methods).

For continents that remain uninvaded, such as Australia, estimating gGO can also serve as a useful tool to identify potential source areas posing the highest risk of contributing optimally preadapted propagules of invasive *D. suzukii* individuals. To illustrate this idea, we note that the two largest exporters of fresh berries to Australia in 2023, after New Zealand and Vietnam (where *D. suzukii* is considered absent), were the USA and Thailand (International Trade Center, 2024). Using Sydney Harbor as a reference environment, since it is an obvious potential sensitive introduction point for pest species in Australia, we derived gGO maps for North America (Figure 6A) and Southeast Asia (Figure 6B). Thailand consistently showed low to moderate gGO values, with no area of very high or extreme values, and was globally characterized by lower gGO values than North America (*t* = −489.48, *p <* 2.2 *×* 10^−16^). Interestingly, in North America, the region with the lowest gGO values was mainly in East US, which is not a major producer of key *D. suzukii* host fruits such as berries.

**Figure 6:**
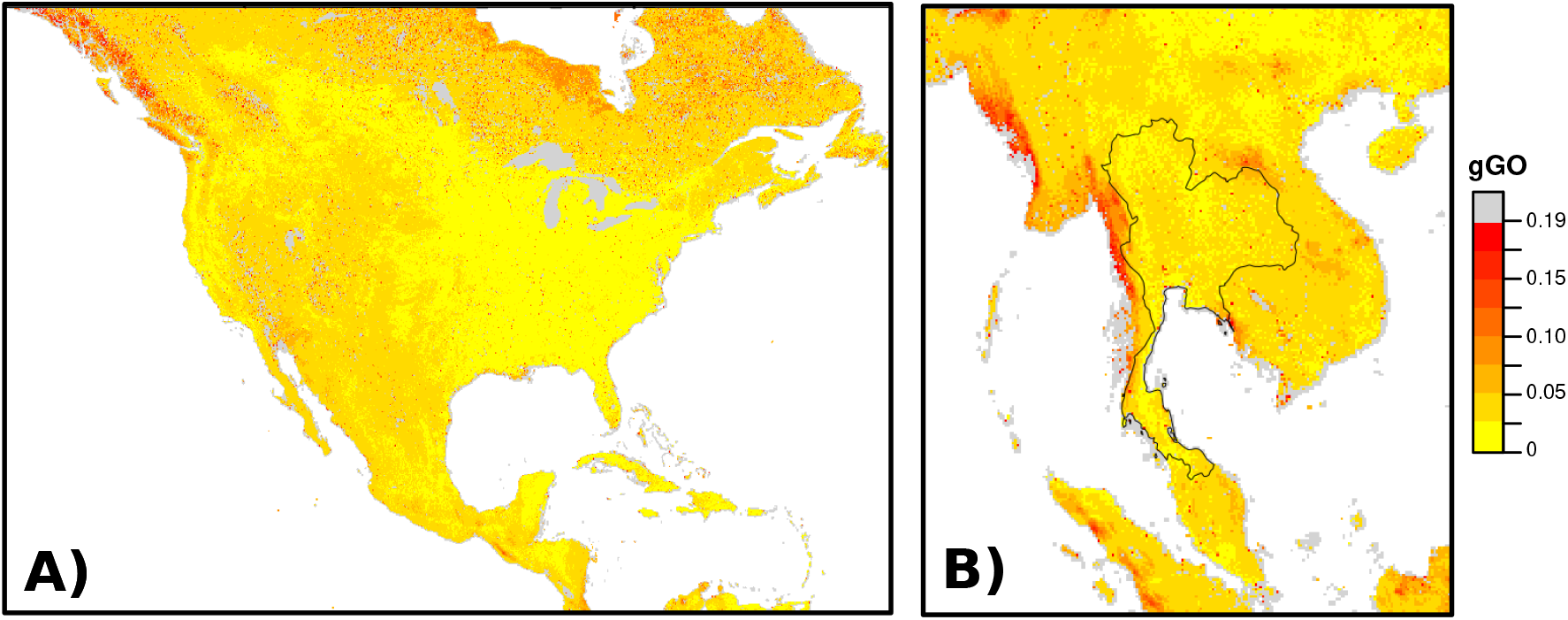
gGO map to assess the areas most at risk as sources of preadapted *D. suzukii* propagules with respect to Australia. gGO values were estimated over areas corresponding to that of the two largest exporters of fresh berries to Australia in 2023 (after New Zealand and Vietnam where *D. suzukii* is absent): North America (A) and Thailand (B, with national borders shown in black). For Australia, the reference environment is that of Sidney’s harbor, an obvious potential sensitive introduction point of pest species in Australia. Gray pixels represent outlying gGO values (as defined in M&M).

## Discussion

The primary goal of this study was to describe and empirically evaluate a population genomics framework to assess the role of genomic adaptation during biological invasions, using the invasive species *D. suzukii* as a biological model. Leveraging recent studies, we expanded the global representation of *D. suzukii* populations by incorporating publicly available Ind-Seq (Lewald *et al*., 2021) and Pool-Seq data (Olazcuaga *et al*., 2020; Feng *et al*., 2024), along with some newly sequenced samples.

### New genomic resources and combination of Ind-Seq and Pool-Seq data

Jointly analyzing heterogeneous datasets (Ind-Seq and Pool-Seq) is statistically challenging and prone to technical biases. The random allele PCA (Skoglund and Jakobsson, 2011), implemented in poolfstat (Gautier *et al*., 2022) has proven effective in accounting for sample heterogeneity when characterizing the global structuring of genetic diversity. However, its reliance on one read per sample per site makes it suboptimal for detecting local adaptive signals or environmental associations. To our knowledge, no current population genomics method fully exploits heterogeneous datasets (i.e. comprising both Ind-Seq and Pool-Seq data) which become more available across species with the growing use of sequencing technologies and the encouraging shift toward open science. We therefore extended BayPass (Gautier, 2015) to jointly analyze Ind-Seq and Pool-Seq data, and further allows modeling Ind-Seq data using either allele counts or genotype likelihoods (GLs). Although the former approach is sufficient for high-coverage data (e.g., *>*20X) where individual genotypes can be reliably called beforehand, the latter properly integrates over genotype uncertainty associated with low to medium coverage data (i.e., *<* 15*X*), which is recommended in a variety of population genomics methods (e.g., Kim *et al*., 2011; Skotte *et al*., 2013; Korneliussen *et al*., 2014; Druet and Gautier, 2017; Meisner and Albrechtsen, 2018)

In order to improve the accuracy of our genomic analyses, we generated a new chromosome-level and annotated assembly for the *D. suzukii* genome from males (to sequence the Y chromosome) of a Japanese strain established in the 1980’s (Ohashi *et al*., 1991), i.e., before the *D. suzukii* invasion outside Asia. This contrasts with previous publicly available assemblies, which were obtained from females of isofemale lines representative of the European or North American invaded areas (e.g. Ometto *et al*., 2013; Chiu et al., 2013; Paris et al., 2020). The new *dsu_isojap1*.*0* assembly can thus be considered as more representative of a *D. suzukii* “ancestral” genome for the native area (disregarding subsequent genetic changes that have occurred in the strain) to identify *de novo* mutation in invasive populations, especially regarding structural variation (including TE content), which we did not examine here as it was outside the focus of the present study. In addition, it provides a significant improvement over existing assemblies and may therefore serve as a valuable reference for other genomic studies on *D. suzukii*.

### Genome-wide scans for selection between native and invaded ranges

For our purposes, the close to optimal contiguity of the *dsu_isojap1*.*0* assembly enabled a better use of the local-score approach of Fariello et al. (2017), which accounts for LD information by combining *C*_2_ statistics at neighboring variants at a whole chromosome scale. This both increased the support and facilitated the delineation of genomic regions associated with the invasive status of populations. In total, we identified 457 significant local-score windows, which were generally small (that is, mainly *<* 20 kb), in agreement with the low level of SNP signal clustering already observed in Olazcuaga et al. (2020). Furthermore, in accordance with the notably distinct subsets of SNPs identified as associated based on 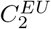 and 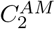 in Olazcuaga et al. (2020), the resulting continent-specific windows exhibited a negligible degree of overlap. Likewise, several highly significant 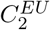 – specific windows (especially on chromosome 2R) disappeared with 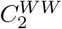, i.e., when adding American samples to the test. This suggests, at least at first sight, a low degree of convergence between the adaptation pathways observed in the two (almost) independently invaded areas by *D. suzukii* (i.e. Europe and America). However, it is important to note that a substantially smaller number of windows were detected when considering continents separately, especially for America. The power of continent-specific analyses are reduced, as illustrated by the weaker *C*_2_ signals observed, (especially for 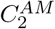), due to the lower number of populations considered for each continent, but also to the stronger neutral genetic structure in the Americas (Figure 2). All of this greatly complicates the assessment of potentially convergent adaptive signals. Also, because *C*_2_ compares the means of standardized population allele frequencies (i.e., corrected for overall structuring of genetic diversity) between two groups (here defined according to their native or invasive status), strong divergence in just a few populations from either group can disproportionately influence the statistic. As a result, a significant signal may be driven by only a subset of populations, rather than being shared across the entire contributing group.

Surprisingly, the annotation of the genomic windows identified in the present study did not reveal any candidate genes in common with those identified by Olazcuaga et al. (2020). This limited overlap appears to result primarily from three methodological differences (see Figure S10 for more details): (i) the use of the local score method, which was not applied in Olazcuaga et al. (2020) but, when retroactively used on the original dataset, recovers overlapping signals with our results; (ii) differences in variant calling pipelines, as VarScan (Koboldt *et al*., 2012), the heuristic-based caller originally used by Olazcuaga *et al*., is expected to be (far) less reliable than the FreeBayes (Garrison and Marth, 2012) haplotype caller we used here, by calling more and potentially lower-quality variants; and (iii) the larger number of populations analyzed in the present study, which increases statistical power and robustness. These differences may explain why we detected signals not captured in the earlier study or, conversely, discarded genes previously supported by only a few variants. In particular, this may explain why we were unable to confirm the strong association signal of the three neighboring SNPs upstream of the *cpo* gene, as these were actually located in a low-complexity region of the 3R chromosome that was poorly assembled and encompassed several highly complex variants in the previous reference assembly in Paris et al. (2020).

Although *cpo*, which seemed at first sight particularly attractive as a functional candidate due to its role in the control of diapause (Olazcuaga *et al*., 2020), was not confirmed, we identified several strong and even more conclusive signals encompassing genes potentially involved in other interesting functions that could be related to invasion success. Among these functions are detoxification, stress response, and pesticide resistance, which can be advantageous traits. For instance, they may enhance tolerance to pesticides, a factor to which *D. suzukii* is regularly exposed and can rapidly develop resistance, as evidenced by laboratory assessments (Haye *et al*., 2016; Deans and Hutchison, 2022). In addition, we identified genes related to gustatory and olfactory behaviors, potentially underpinning *D. suzukii* ‘s generalist feeding strategies by increasing its ability to detect and utilize a variety of host fruits in the environments it has invaded. We also discovered genes associated with immune responses that may be important, as interaction with new pathogens in newly invaded environments can impede or promote invasion success, depending on whether the allocation of resources to immune function detracts from other vital traits or improves adaptive survival in these new habitats. However, it is important to keep in mind that since *C*_2_ does not indicate in which group (native or invasive) selection occurred, we generally interpret these loci as potentially linked to invasion success, as they all show differentiation between native and invasive ranges. But in any case, *C*_2_ highlights loci experiencing shifts in selective pressure between ranges, which remains highly relevant for understanding invasion biology regardless of the selection’s direction. Finally, association analyzes such as the one performed here based on *C*_2_ are expected to provide a very partial view of the genetic basis of invasive success of a species, since the underlying traits are expected to be multiple and highly polygenic. Association analyses thus primarily aim at identifying a few important genes, such as the few candidates discussed above, that may deserve special interest for future validation studies to better characterize the traits they control.

### Genomic Offset to understand past invasion

To provide a global picture of the (possible) role of genomic adaptation in the invasive success of *D. suzukii*, we considered an alternative approach based on modeling the relationship between whole-genome genomic variation and environmental variables among populations, building on recent work on estimating population maladaptation using the Genomic Offset statistic (e.g. Capblancq *et al*., 2020), here in its gGO definition (Gain *et al*., 2023). Assuming that i) (some of) the considered environmental variables are among the main drivers of invasion success; ii) populations adapt partly genetically to these variables; and iii) genomic data are available from representative locally adapted native and invaded populations, then a low (resp. high) gGO value between two environments can be interpreted as a relative measure of (genetic) preadaptation (resp. maladaptation) for a population that would be moved from one environment to another. Accordingly, in a simulation study, Camus et al. (2024) showed that gGO correlates with the probability of invasive population establishment and can be efficiently estimated from the GEA regression model implemented in BayPass. Its new extension integrating Ind-Seq and Pool-Seq now enables gGO estimates from combined datasets, which was not possible (or highly suboptimal) with other available methods.

The worldwide invasion of *D. suzukii* provided us with a valuable case study to illustrate how gGO-based analyzes can provide insights into the level of (genomic) adaptive challenge that poses the environment of a new invaded area to a population adapted to a given region of the native area. Interestingly, despite evidence that some genes were associated with invasive status and thus likely experience different selection pressures between ranges, gGO indicated that genetic (pre)adaptation may have played an important role in *D. suzukii* ‘s invasion success since gGO values in Europe and North America remained mostly low/moderate and spatially homogeneous. This suggests that the adaptive challenge for *D. suzukii* populations to invade Europe and North America was relatively limited, and that no particular region appears to have offered a more favorable adaptive context than others to propagules originating from populations adapted to the environments of the presumed source native area. This aligns with the rapid invasion of *D. suzukii*. While not contradicting the findings of Feng et al. (2024), who identified many candidate SNPs associated with environmental variables, our results suggest that the underlying genomic regions were likely not crucial in the invasion process. Our conclusions are also in line with a previous study using species distribution modeling in *D. suzukii*, which indicated the absence of an obvious climatic niche shift between Asia and the first observation points in Europe and the US (Fraimout and Monnet, 2018).

### Genomic Offset to predict future invasions

gGO maps that we derived for Africa and South America illustrate how gGO can be used to identify the most vulnerable regions in large areas that have not yet been (fully) invaded. According to our results, relatively low gGO values (that is, high risk) were found in large parts of these continents (for example, the Amazon basin in South America or the Great Lakes region in Africa) where *D. suzukii* is currently absent, but could potentially establish rapidly. Some post hoc surveillance of these regions would be required to assess the extent to which these gGO-based predictions are informative. Indeed, while we report low gGO values in some recently invaded regions (e.g., South Africa), these were moderate in others (e.g., Patagonia, Chile). In addition, the choice of the reference source environment from which the (preadapted) invading population would originate can be critical. For simplicity, for both Africa and South America, we assumed that the reference source environment is the one where populations have been established for at least two consecutive years. Consideration of other source environments may result in slightly to substantially different gGO maps. For example, in South America, although the first record of *D. suzukii* dates back to 2013 (Deprá *et al*., 2014), it was reported just one year later, in 2014, in northern Patagonia (Lavagnino *et al*., 2018), approximately 500 km from the AR-Gol population studied here and sampled in 2023. Given the rapid expansion of *D. suzukii* in Patagonia, AR-Gol could also represent a suitable reference environment to compute a gGO map of invasion across South America. This could potentially lead to even more pessimistic predictions than those based on the Nova Veneza (Brazil) reference environment used in Figure 5B, meaning that an even greater number of regions might be predicted to have low gGO values and thus be at higher risk of invasion.

The computation of gGO maps for the uninvaded Australian continent, using a reference environment associated with the northwestern American US-Wa population (a key fresh fruit import region), also supports a high overall risk of invasion from there. Densely populated areas of eastern and southern Australia that are likely targets of significant food trade exhibited low gGO values, suggesting that such a US-Wa-like *D. suzukii* population could easily establish there and gradually spread throughout the country. Previous research, which did not incorporate genomic data, similarly identified a high risk of *D. suzukii* invasion in eastern and southern Australia, particularly through passive human-mediated pathways (Maino *et al*., 2021).

The gGO can also be used to identify the region of origin of a (locally adapted) population that would be the most (genomically) pre-adapted to a specific target area (see also Gautier *et al*., 2024, for another recent application). Such information can be useful for IPM practice to increase vigilance on imports from the highlighted high risk regions (i.e., those with low gGO values and where *D. suzukii* is established). Our gGO analysis suggests that invasions from Thailand present a greater (adaptive) risk than those from the US, implying that Australian management should also focus on monitoring imports from Thailand. In any case, this analysis serves more as an illustration of gGO-based IPM applications than as a source of actionable information for Australia, which already enforces strict control measures on most goods at risk of *D. suzukii* infestation. In a more general perspective, the sequencing of newly established populations could serve as an empirical method to validate gGO predictions in biological invasions. By comparing the most likely sources of invasion inferred from genetic data only, with those expected through gGO computations accounting for environmental variables (e.g., Gautier *et al*., 2024), we could assess the relevance of gGO-based predictions in real-world scenarios.

### Application and limitations of Genomic Offset in invasion biology

In general, approaches that rely on Genomic Offset statistics appear promising for understanding and predicting *D. suzukii* colonization in different environments. From a biological perspective, gGO estimates made sense globally, as regions with high estimated gGO values corresponded to environmental characteristics known to be physiologically challenging for *D. suzukii*. In particular, high to extreme gGO values were observed in cold regions, which are inhospitable to *D. suzukii* due to its “chill-susceptibility” (Kimura, 2004; Enriquez and Colinet, 2017; Ryan *et al*., 2016; Winkler *et al*., 2020). However, in our analysis, temperature-related variables had relatively low importance, suggesting that the reduced suitability of these regions is more likely driven by low precipitation, low humidity and unfavorable land use or agricultural production, all of which were identified as stronger predictors. Similarly, high gGO values were found in arid and semi-arid desert regions, which are also unsuitable for *D. suzukii* due to its low tolerance to drought and high temperatures (Winkler *et al*., 2020; Fanning et al., 2019; Eben et al., 2018).

It should be noted that predictions of potential future distributions based on Ecological Niche Models (ENM) show low to no suitability for some areas that were associated with moderate or low gGO values in our analyzes, namely tropical regions such as the Amazon basin or Central Africa (Nair and Peterson, 2023; Ørsted and Ørsted, 2019). This discrepancy likely arises from fundamental differences between ENM and Genomic Offset approaches, which do not rely on the same data to model the relationship between populations and their environments. On the one hand, because ENMs rely on occurrence data, they can underestimate environmental suitability in regions where sampling is less intensive (Dubos *et al*., 2022), which is likely the case of these tropical zones, overlooking the fact that the climatic characteristics of these regions are not necessarily limiting for the species, as indicated by gGO. On the other hand, the Genomic Offset based method focuses on a limited set of sequenced populations, which may limit its ability to capture the full ecological niche of the species. Moreover, it is important to keep in mind that current Genomic Offset metrics do not provide an absolute quantitative measure of invasion risk or adaptation difficulty. As a result, gGO values can only be interpreted relatively, i.e., by ranking environments according to their susceptibility to invasion by source populations that are (genetically) adapted to a given reference region. The exact conditions under which this relative interpretation is theoretically valid also need to be further explored and could, for example, be challenged if a given target environment is particularly suitable for a species, regardless of the genetic makeup of the population (Lotterhos, 2024).

Other factors not included in the models underlying gGO calculations may be critical for successful invasion and should also be considered to better explain or predict the establishment of invasive species. First, some pivotal environmental variables could have been ignored in the GEA, which may have a decisive impact on the predictive performance of gGO (Camus *et al*., 2024). For example, we could not directly incorporate specific (wild and cultivated) fruit-related variables other than land use and agricultural production due to data availability limitations on the considered continental scales. Given that small fruits availability is an important component of *D. suzukii* fitness, its inclusion in future gGO analyzes could improve the precision of invasion predictions. In the same vein, it is worth noting that many of the top significant candidate genes detected by our *C*_2_ genome scan are related to functions that may be crucial for the species invasion success (e.g., pesticide resistance, immune response, dispersal, or feeding behavior) but are not considered in the gGO analysis. Integrating environmental descriptors that are directly linked to these functions, for instance, pesticide levels, might provide a different picture of the actual adaptive challenge encountered during the invasion(s), with higher and more heterogeneous gGO values. Innate characteristics of the species, including phenotypic plasticity, can also play an important role for invasion success and may be difficult (if not impossible) to include in gGO computations. Indeed, *D. suzukii* shows several characteristics that may have facilitated its successful invasion, such as its ability to i) tolerate cold environments through mechanisms such as overwintering or diapause (Little *et al*., 2020; Jakobs *et al*., 2015; Stockton *et al*., 2019a; Rossi-Stacconi et al., 2016); ii) exploit a broad trophic niche with host plants that span several families and many fruit species (Olazcuaga *et al*., 2023); and iii) even subsist on a diet without fruits (Poyet *et al*., 2015; Stockton et al., 2019b).

Finally, contingent and extrinsic factors are also critical to successfully invade a new area. At the very least, enough individuals (of both sexes for sexual species) must first be able to reach it. In the case of *D. suzukii*, its widespread distribution is likely driven by human-mediated passive dispersal rather than active movement (Adrion *et al*., 2014; Calabria *et al*., 2012; Maino *et al*., 2021). This form of assisted migration can significantly affect the dynamics of invasion, influencing which areas are invaded first, and the spread of the species in a newly invaded area. In addition, the number of introduced individuals (propagule size) and their arrival patterns (number and spatio-temporal distribution of introduction events), which together define the so-called “propagule pressure”, have also been shown to be a crucial factor in the establishment of invasive populations (Simberloff, 2009; Estoup *et al*., 2016). In fact, high propagule pressure can buffer adaptive challenges, as shown in our recent simulation study, where the performance of Genomic Offset-based analyzes in predicting the probability of invasive populations being established was found to be reduced for scenarios involving the single introduction of larger numbers of individuals (Camus *et al*., 2024). For instance, even though some populations from Thailand may be more preadapted to Australia, populations from America may be more likely to become established in Australia due to greater fruit infestation and/or higher fruit imports and consequently greater propagule pressure.

## Conclusion

Despite these limitations and the avenue for further refinement, we believe that our findings provide valuable insights into the invasion success of *D. suzukii* and more generally illustrate and highlight the effectiveness and promise of GEA and gGO-based analyzes to elucidate the factors underpinning the dynamic of past and future biological invasions. Our framework for population genomic analysis and our recent development of methodological tools are inherently generic. As such, they can be applied to any other invasive species in order to help guide management decisions.

## Data availability

All data were deposited in the SRA repository (project PRJNA1032894). These included Pacbio HiFi raw reads (Run ID: SRR29552229) and the HiC library paired-end sequences (Run ID: SRR29552230) used for the de novo assembly of the Japanese strain, and all new Pool-Seq and Ind-Seq data (see accession details in Tables S1 and S2). The annotated *dsu_isojap1*.*0* whole genome assembly is publicly available from the NCBI repository under the ID GCA 043229665.1 (https://www.ncbi.nlm.nih.gov/datasets/genome/GCF_043229965.1/). The newly developed version (v3.0) of BayPass software is publicly available from https://forge.inrae.fr/mathieu.gautier/baypass_public

## Acknowledgments

We wish to thank Toshiro Aigaki (TMU, Tokyo, Japan), Nicolas Gompel (Bonn University, Germany) and Benjamin Prud’homme (IBDML, Université Aix-Marseille, France) for providing and having maintained individuals from the Japanese isofemale line used for the genome assembly. We are also grateful to Benjamin Prud’homme for providing *D. subpulchrella* individuals, and Dr Marina Stamenković-Radak, Dr Mihailo Jelić, and Mina Rakić for their assistance in collecting the sample from Serbia. Mathieu Gautier, Simon Boitard and Arnaud Estoup have benefited from State aid managed by the Agence Nationale de la Recherche under the France 2030 programme, reference ANR-23-EXMA-0002 (“AgroStat” project of the Programme et Equipements Prioritaires de Recherche “MathsVives”). Svitlana Serga was supported by the PAUSE-ANR Ukraine Program. This work was performed in collaboration with the GeT core facility, Toulouse, France (GeT, https://doi.org/10.15454/1.5572370921303193E12). MGX and GeT core facility acknowledge financial support from France Génomique National infrastructure, funded as part of “Investissement d’avenir” program managed by Agence Nationale pour la Recherche (contract ANR-10-INBS-09). We are also grateful to the genotoul bioinformatics platform Toulouse Occitanie (Bioinfo Genotoul, https://doi.org/10.15454/1.5572369328961167E12) for providing computing resources. We finally thank the DrosEU consortium for helpful discussions during stimulating meetings.

## Bibliography

Adrion, J. R., Kousathanas, A., Pascual, M., Burrack, H. J., Haddad, N. M., Bergland, A. O., Machado, H., Sackton, T. B., Schlenke, T. A., Watada, M., Wegmann, D., and Singh, N. D. 2014. Drosophila suzukii: The Genetic Footprint of a Recent, Worldwide Invasion. Molecular Biology and Evolution, 31(12): 3148–3163.

Aouari, I., Barech, G., and Khaldi, M. 2022. First record of the agricultural pest Drosophila suzukii (Matsumura, 1931) (Diptera: Drosophilidae) in Algeria. EPPO Bulletin, 52(2): 471–478.

Asplen, M. K., Anfora, G., Biondi, A., Choi, D. S., Chu, D., Daane, K. M., Gibert, P., Gutierrez, A. P., Hoelmer, K. A., Hutchison, W. D., Isaacs, R., Jiang, Z. L., Kárpáti, Z., Kimura, M. T., Pascual, M., Philips, C. R., Plantamp, C., Ponti, L., Vétek, G., Vogt, H., Walton, V. M., Yu, Y., Zappala, L., and Desneux, N. 2015. Invasion biology of spotted wing Drosophila (Drosophila suzukii): a global perspective and future priorities. Journal of Pest Science, 88(3): 469–494.

Barrett, R. D. H. and Schluter, D. 2008. Adaptation from standing genetic variation. Trends in Ecology & Evolution, 23(1): 38–44.

Bayona-Feliu, A., Casas-Lamesa, A., Carbonell, A., Climent-Cantó, P., Tatarski, M., Pérez-Montero, S., Azorín, F., and Bernués, J. 2016. Histone h1: lessons from drosophila. Biochimica et Biophysica Acta (BBA)-Gene Regulatory Mechanisms, 1859(3): 526–532.

Bhutkar, A., Schaeffer, S. W., Russo, S. M., Xu, M., Smith, T. F., and Gelbart, W. M. 2008. Chromosomal rearrangement inferred from comparisons of 12 Drosophila genomes. Genetics, 179(3): 1657–1680.

Boughdad, A., Haddi, K., El Bouazzati, A., Nassiri, A., Tahiri, A., El Anbri, C., Eddaya, T., Zaid, A., and Biondi, A. 2021. First record of the invasive spotted wing Drosophila infesting berry crops in Africa. Journal of Pest Science, 94(2): 261–271.

Bradshaw, C. J. A., Leroy, B., Bellard, C., Roiz, D., Albert, C., Fournier, A., Barbet-Massin, M., Salles, J.-M., Simard, F., and Courchamp, F. 2016. Massive yet grossly underestimated global costs of invasive insects. Nature Communications, 7(1): 12986.

Bruce, T. J. A. 2010. Tackling the threat to food security caused by crop pests in the new millennium. Food Security, 2(2): 133–141.

Cabanettes, F. and Klopp, C. 2018. D-GENIES: Dot plot large GENomes in an interactive, efficient and simple way. PeerJ Preprints, 6: e26567v1.

Calabria, G., Máca, J., Bächli, G., Serra, L., and Pascual, M. 2012. First records of the potential pest species drosophila suzukii (diptera: Drosophilidae) in europe. Journal of Applied entomology, 136(1-2): 139–147.

Camus, L., Gautier, M., and Boitard, S. 2024. Predicting species invasiveness with genomic data: Is genomic offset related to establishment probability? Evolutionary Applications, 17(6): e13709.

Capblancq, T., Fitzpatrick, M. C., Bay, R. A., Exposito-Alonso, M., and Keller, S. R. 2020. Genomic Prediction of (Mal)Adaptation Across Current and Future Climatic Landscapes. Annual Review of Ecology, Evolution, and Systematics, 51(1): 245–269.

Challis, R., Richards, E., Rajan, J., Cochrane, G., and Blaxter, M. 2020. BlobToolKit – Interactive Quality Assessment of Genome Assemblies. G3, 10(4): 1361–1374.

Chang, C.-H., Chavan, A., Palladino, J., Wei, X., Martins, N. M. C., Santinello, B., Chen, C.-C., Erceg, J., Beliveau, B. J., Wu, C.-T., Larracuente, A. M., and Mellone, B. G. 2019. Islands of retroelements are major components of drosophila centromeres. PLOS Biology, 17(5): 1–40.

Chen, S., Zhou, Y., Chen, Y., and Gu, J. 2018. fastp: an ultra-fast all-in-one fastq preprocessor. Bioinformatics, 34(17): i884–i890.

Cheng, H., Concepcion, G. T., Feng, X., Zhang, H., and Li, H. 2021. Haplotype-resolved de novo assembly using phased assembly graphs with hifiasm. Nature Methods, 18(2): 170–175.

Chiu, J. C., Jiang, X., Zhao, L., Hamm, C. A., Cridland, J. M., Saelao, P., Hamby, K. A., Lee, E. K., Kwok, R. S., Zhang, G., Zalom, F. G., Walton, V. M., and Begun, D. J. 2013. Genome of Drosophila suzukii, the spotted wing drosophila. G3, 3(12): 2257–71.

Colautti, R. I. and Barrett, S. C. H. 2013. Rapid Adaptation to Climate Facilitates Range Expansion of an Invasive Plant. Science, 342(6156): 364–366.

Deans, C. and Hutchison, W. D. 2022. Propensity for resistance development in the invasive berry pest, spotted-wing drosophila (drosophila suzukii), under laboratory selection. Pest management science, 78(12): 5203–5212.

Deprá, M., Poppe, J., Schmitz, H., De Toni, D., and Valente, V. 2014. The first records of the invasive pest Drosophila suzukii in the South American continent. J Pest Sci, 87.

Diagne, C., Leroy, B., Vaissiere, A.-C., Gozlan, R. E., Roiz, D., Jarić, I., Salles, J.-M., Bradshaw, C. J. A., and Courchamp, F. 2021. High and rising economic costs of biological invasions worldwide. Nature, 592(7855): 571–576.

Drosopoulou, E., Gariou-Papalexiou, A., Karamoustou, E., Gouvi, G., Augustinos, A. A., Bourtzis, K., and Zacharopoulou, A. 2019. The chromosomes of Drosophila suzukii (Diptera: Drosophilidae): detailed photographic polytene chromosomal maps and in situ hybridization data. Molecular Genetics and Genomics, 294(6): 1535–1546.

Druet, T. and Gautier, M. 2017. A model-based approach to characterize individual inbreeding at both global and local genomic scales. Molecular Ecology, 26(20): 5820–5841.

Dubos, N., Préau, C., Lenormand, M., Papuga, G., Monsarrat, S., Denelle, P., Le Louarn, M., Heremans, S., May, R., Roche, P., et al. 2022. Assessing the effect of sample bias correction in species distribution models. Ecological Indicators, 145: 109487.

Eben, A., Reifenrath, M., Briem, F., Pink, S., and Vogt, H. 2018. Response of d rosophila suzukii (d iptera: D rosophilidae) to extreme heat and dryness. Agricultural and Forest Entomology, 20(1): 113–121.

Enriquez, T. and Colinet, H. 2017. Basal tolerance to heat and cold exposure of the spotted wing drosophila, drosophila suzukii. PeerJ, 5: e3112.

Estoup, A., Ravigné, V., Hufbauer, R., Vitalis, R., Gautier, M., and Facon, B. 2016. Is There a Genetic Paradox of Biological Invasion? Annual Review of Ecology, Evolution, and Systematics, 47(Volume 47, 2016): 51–72.

Fanning, P. D., Johnson, A. E., Luttinen, B. E., Espeland, E. M., Jahn, N. T., and Isaacs, R. 2019. Behavioral and physiological resistance to desiccation in spotted wing drosophila (diptera: Drosophilidae). Environmental entomology, 48(4): 792–798.

Fariello, M. I., Boitard, S., Mercier, S., Robelin, D., Faraut, T., Arnould, C., Recoquillay, J., Bouchez, O., Salin, G., Dehais, P., Gourichon, D., Leroux, S., Pitel, F., Leterrier, C., and San-Cristobal, M. 2017. Accounting for linkage disequilibrium in genome scans for selection without individual genotypes: The local score approach. Molecular Ecology, 26(14): 3700–3714.

Feng, S., DeGrey, S. P., Guédot, C., Schoville, S. D., and Pool, J. E. 2023. Genomic Diversity Illuminates the Species History and Environmental Adaptation of Drosophila suzukii. bioRxiv, 2023.07.03.547576.

Feng, S., DeGrey, S. P., Guédot, C., Schoville, S. D., and Pool, J. E. 2024. Genomic Diversity Illuminates the Environmental Adaptation of Drosophila suzukii. Genome Biology and Evolution, 16(9): evae195.

Fitzpatrick, M. C. and Keller, S. R. 2015. Ecological genomics meets community-level modelling of biodiversity: mapping the genomic landscape of current and future environmental adaptation. Ecology Letters, 18(1): 1–16.

Fitzpatrick, M. C., Chhatre, V. E., Soolanayakanahally, R. Y., and Keller, S. R. 2021. Experimental support for genomic prediction of climate maladaptation using the machine learning approach Gradient Forests. Molecular Ecology Resources, 21(8): 2749–2765.

Fraimout, A. and Monnet, A.-C. 2018. Accounting for intraspecific variation to quantify niche dynamics along the invasion routes of drosophila suzukii. Biological Invasions, 20(10): 2963–2979.

Fraimout, A., Debat, V., Fellous, S., Hufbauer, R. A., Foucaud, J., Pudlo, P., Marin, J.-M., Price, D. K., Cattel, J., Chen, X., Deprá, M., François Duyck, P., Guedot, C., Kenis, M., Kimura, M. T., Loeb, G., Loiseau, A., Martinez-Sañudo, I., Pascual, M., Polihronakis Richmond, M., Shearer, P., Singh, N., Tamura, K., Xuéreb, A., Zhang, J., and Estoup, A. 2017. Deciphering the Routes of invasion of Drosophila suzukii by Means of ABC Random Forest. Molecular Biology and Evolution, 34(4): 980–996.

Gain, C., Rhoné, B., Cubry, P., Salazar, I., Forbes, F., Vigouroux, Y., Jay, F., and François, O. 2023. A Quantitative Theory for Genomic Offset Statistics. Molecular Biology and Evolution, 40(6): msad140.

Garrison, E. and Marth, G. 2012. Haplotype-based variant detection from short-read sequencing. arXiv, 1207.3907.

Gautier, M. 2015. Genome-Wide Scan for Adaptive Divergence and Association with Population-Specific Covariates. Genetics, 201(4): 1555–1579.

Gautier, M. 2023. Efficient k-mer based curation of raw sequence data: application in Drosophila suzukii. Peer Community Journal, 3: e79.

Gautier, M., Yamaguchi, J., Foucaud, J., Loiseau, A., Ausset, A., Facon, B., Gschloessl, B., Lagnel, J., Loire, E., Parrinello, H., Severac, D., Lopez-Roques, C., Donnadieu, C., Manno, M., Berges, H., Gharbi, K., Lawson-Handley, L., Zang, L.-S., Vogel, H., Estoup, A., and Prud’homme, B. 2018. The Genomic Basis of Color Pattern Polymorphism in the Harlequin Ladybird. Current Biology, 28(20): 3296–3302.e7.

Gautier, M., Vitalis, R., Flori, L., and Estoup, A. 2022. f-Statistics estimation and admixture graph construction with Pool-Seq or allele count data using the R package poolfstat. Molecular Ecology Resources, 22(4): 1394–1416.

Gautier, M., Micol, T., Camus, L., Moazami-Goudarzi, K., Naves, M., Guéret, E., Engelen, S., Lemainque, A., Colas, F., Flori, L., and Druet, T. 2024. Genomic Reconstruction of the Successful Establishment of a Feralized Bovine Population on the Subantarctic Island of Amsterdam. Molecular Biology and Evolution, 41(7): msae121.

Haubrock, P. J., Turbelin, A. J., Cuthbert, R. N., Novoa, A., Taylor, N. G., Angulo, E., Ballesteros-Mejia, L., Bodey, T. W., Capinha, C., Diagne, C., Essl, F., Golivets, M., Kirichenko, N., Kourantidou, M., Leroy, B., Renault, D., Verbrugge, L., and Courchamp, F. 2021. Economic costs of invasive alien species across Europe. NeoBiota, 67: 153–190.

Haye, T., Girod, P., Cuthbertson, A., Wang, X., Daane, K., Hoelmer, K., Baroffio, C., Zhang, J., and Desneux, N. 2016. Current swd ipm tactics and their practical implementation in fruit crops across different regions around the world. Journal of Pest Science, 89: 643–651.

Hofmeister, N. R., Werner, S. J., and Lovette, I. J. 2021. Environmental correlates of genetic variation in the invasive European starling in North America. Molecular Ecology, 30(5): 1251–1263.

Hoskins, R. A., Carlson, J. W., Wan, K. H., Park, S., Mendez, I., Galle, S. E., Booth, B. W., Pfeiffer, B. D., George, R. A., Svirskas, R., Krzywinski, M., Schein, J., Accardo, M. C., Damia, E., Messina, G., Méndez-Lago, M., de Pablos, B., Demakova, O. V., Andreyeva, E. N., Boldyreva, L. V., Marra, M., Carvalho, A. B., Dimitri, P., Villasante, A., Zhimulev, I. F., Rubin, G. M., Karpen, G. H., and Celniker, S. E. 2015. The release 6 reference sequence of the drosophila melanogaster genome. Genome Research, 25(3): 445–58.

Hufbauer, R. A., Facon, B., Ravigné, V., Turgeon, J., Foucaud, J., Lee, C. E., Rey, O., and Estoup, A. 2012. Anthropogenically induced adaptation to invade (AIAI): contemporary adaptation to human-altered habitats within the native range can promote invasions. Evolutionary Applications, 5(1): 89–101. International Trade Center 2024. Trade Map. Accessed: 2024-07-22.

IPCC 2024. International Plant Protection Convention country report: Notification of the detection of Drosophila suzukii, the Spotted Wing Drosophila in the Republic of South Africa. Technical Report ZAF-58/2, National Plant Protection Organization of South Africa.

Jakobs, R., Gariepy, T. D., and Sinclair, B. J. 2015. Adult plasticity of cold tolerance in a continentaltemperate population of drosophila suzukii. Journal of Insect Physiology, 79: 1–9.

Kim, S. Y., Lohmueller, K. E., Albrechtsen, A., Li, Y., Korneliussen, T., Tian, G., Grarup, N., Jiang, T., Andersen, G., Witte, D., Jorgensen, T., Hansen, T., Pedersen, O., Wang, J., and Nielsen, R. 2011. Estimation of allele frequency and association mapping using next-generation sequencing data. BMC Bioinformatics, 12(1): 231.

Kimura, M. T. 2004. Cold and heat tolerance of drosophilid flies with reference to their latitudinal distributions. Oecologia, 140: 442–449.

Koboldt, D. C., Zhang, Q., Larson, D. E., Shen, D., McLellan, M. D., Lin, L., Miller, C. A., Mardis, E. R., Ding, L., and Wilson, R. K. 2012. Varscan 2: somatic mutation and copy number alteration discovery in cancer by exome sequencing. Genome Research, 22(3): 568–76.

Korneliussen, T. S., Albrechtsen, A., and Nielsen, R. 2014. Angsd: Analysis of next generation sequencing data. BMC Bioinformatics, 15(1): 356.

Kwadha, C. A., Okwaro, L. A., Kleman, I., Rehermann, G., Revadi, S., Ndlela, S., Khamis, F. M., Nderitu, P. W., Kasina, M., George, M. K., Kithusi, G. G., Mohamed, S. A., Lattorff, H. M. G., and Becher, P. G. 2021. Detection of the spotted wing drosophila, Drosophila suzukii, in continental sub-Saharan Africa. Journal of Pest Science, 94(2): 251–259.

Lachmuth, S., Capblancq, T., Prakash, A., Keller, S. R., and Fitzpatrick, M. C. 2023. Novel genomic offset metrics integrate local adaptation into habitat suitability forecasts and inform assisted migration. Ecological Monographs, 94(1): e1593.

Láruson, A. J., Yeaman, S., and Lotterhos, K. E. 2020. The Importance of Genetic Redundancy in Evolution. Trends in Ecology & Evolution, 35(9): 809–822.

Lavagnino, N. J., Cichón, L. I., De La Vega, G. J., Garrido, S. A., Fanara, J. J., et al. 2018. New records of the invasive pest Drosophila suzukii (Matsumura)(Diptera: Drosophilidae) in the South American continent. Revista de la Sociedad Entomológica Argentina, 77(1): 27.

Lewald, K. M., Abrieux, A., Wilson, D. A., Lee, Y., Conner, W. R., Andreazza, F., Beers, E. H., Burrack, H. J., Daane, K. M., Diepenbrock, L., Drummond, F. A., Fanning, P. D., Gaffney, M. T., Hesler, S. P., Ioriatti, C., Isaacs, R., Little, B. A., Loeb, G. M., Miller, B., Nava, D. E., Rendon, D., Sial, A. A., Bezerra da Silva, C. S., Stockton, D. G., Van Timmeren, S., Wallingford, A., Walton, V. M., Wang, X., Zhao, B., Zalom, F. G., and Chiu, J. C. 2021. Population genomics of Drosophila suzukii reveal longitudinal population structure and signals of migrations in and out of the continental United States. G3, 11(12): jkab343.

Li, H. 2021. New strategies to improve minimap2 alignment accuracy. Bioinformatics, 37(23): 4572–4574.

Li, H., Handsaker, B., Wysoker, A., Fennell, T., Ruan, J., Homer, N., Marth, G., Abecasis, G., and Durbin, R. 2009. The sequence alignment/map format and samtools. Bioinformatics, 25(16): 2078–9.

Lind, B. M. and Lotterhos, K. E. 2024. The limits of predicting maladaptation to future environments with genomic data. bioRxiv, 2024.01.30.577973.

Little, C. M., Chapman, T. W., and Hillier, N. K. 2020. Plasticity is key to success of drosophila suzukii (diptera: Drosophilidae) invasion. Journal of Insect Science, 20(3): 5.

Lotterhos, K. E. 2023. The paradox of adaptive trait clines with nonclinal patterns in the underlying genes. Proceedings of the National Academy of Sciences, 120(12): e2220313120.

Lotterhos, K. E. 2024. Interpretation issues with “genomic vulnerability” arise from conceptual issues in local adaptation and maladaptation. Evolution Letters, 8(3): 331–339.

Ma, L., Cao, L.-J., Hoffmann, A. A., Gong, Y.-J., Chen, J.-C., Chen, H.-S., Wang, X.-B., Zeng, A.-P., Wei, S.-J., and Zhou, Z.-S. 2020. Rapid and strong population genetic differentiation and genomic signatures of climatic adaptation in an invasive mealybug. Diversity and Distributions, 26(5): 610–622.

Maino, J. L., Schouten, R., and Umina, P. 2021. Predicting the global invasion of drosophila suzukii to improve australian biosecurity preparedness. Journal of Applied Ecology, 58(4): 789–800.

Manni, M., Berkeley, M. R., Seppey, M., Simão, F. A., and Zdobnov, E. M. 2021. BUSCO Update: Novel and Streamlined Workflows along with Broader and Deeper Phylogenetic Coverage for Scoring of Eukaryotic, Prokaryotic, and Viral Genomes. Molecular Biology and Evolution, 38(10): 4647–4654.

Mazza, G., Tricarico, E., Genovesi, P., and Gherardi, F. 2014. Biological invaders are threats to human health: an overview. Ethology Ecology & Evolution.

Meisner, J. and Albrechtsen, A. 2018. Inferring Population Structure and Admixture Proportions in Low-Depth NGS Data. Genetics, 210(2): 719–731.

Mérel, V., Gibert, P., Buch, I., Rodriguez Rada, V., Estoup, A., Gautier, M., Fablet, M., Boulesteix, M., and Vieira, C. 2021. The Worldwide Invasion of Drosophila suzukii Is Accompanied by a Large Increase of Transposable Element Load and a Small Number of Putatively Adaptive Insertions. Molecular Biology and Evolution, 38(10): 4252–4267.

Mollot, G., Pantel, J. H., and Romanuk, T. N. 2017. Chapter Two - The Effects of Invasive Species on the Decline in Species Richness: A Global Meta-Analysis. In D. A. Bohan, A. J. Dumbrell, and F. Massol, editors, Advances in Ecological Research, volume 56 of Networks of Invasion: A Synthesis of Concepts, pages 61–83. Academic Press.

Nair, R. R. and Peterson, A. T. 2023. Mapping the global distribution of invasive pest drosophila suzukii and parasitoid leptopilina japonica: implications for biological control. PeerJ, 11: e15222.

Ohashi, Y. Y., Haino-Fukushima, K., and Fuyama, Y. 1991. Purification and charachterization of an ovulation stimulating substance from the male accessory glands of drosophila suzukii. Insect Biochemistry, 21(4): 413–419.

Olazcuaga, L., Loiseau, A., Parrinello, H., Paris, M., Fraimout, A., Guedot, C., Diepenbrock, L. M., Kenis, M., Zhang, J., Chen, X., Borowiec, N., Facon, B., Vogt, H., Price, D. K., Vogel, H., Prud’homme, B., Estoup, A., and Gautier, M. 2020. A Whole-Genome Scan for Association with Invasion Success in the Fruit Fly Drosophila suzukii Using Contrasts of Allele Frequencies Corrected for Population Structure. Molecular Biology and Evolution, 37(8): 2369–2385.

Olazcuaga, L., Baltenweck, R., Leménager, N., Maia-Grondard, A., Claudel, P., Hugueney, P., and Foucaud, J. 2023. Metabolic consequences of various fruit-based diets in a generalist insect species. eLife, 12: e84370.

Ometto, L., Cestaro, A., Ramasamy, S., Grassi, A., Revadi, S., Siozios, S., Moretto, M., Fontana, P., Varotto, C., Pisani, D., Dekker, T., Wrobel, N., Viola, R., Pertot, I., Cavalieri, D., Blaxter, M., Anfora, G., and Rota-Stabelli, O. 2013. Linking Genomics and Ecology to Investigate the Complex Evolution of an Invasive Drosophila Pest. Genome Biology and Evolution, 5(4): 745–757.

Ørsted, I. V. and Ørsted, M. 2019. Species distribution models of the spotted wing drosophila (drosophila suzukii, diptera: Drosophilidae) in its native and invasive range reveal an ecological niche shift. Journal of applied ecology, 56(2): 423–435.

Paini, D. R., Sheppard, A. W., Cook, D. C., De Barro, P. J., Worner, S. P., and Thomas, M. B. 2016. Global threat to agriculture from invasive species. Proceedings of the National Academy of Sciences, 113(27): 7575–7579.

Paris, M., Boyer, R., Jaenichen, R., Wolf, J., Karageorgi, M., Green, J., Cagnon, M., Parinello, H., Estoup, A., Gautier, M., Gompel, N., and Prud’homme, B. 2020. Near-chromosome level genome assembly of the fruit pest Drosophila suzukii using long-read sequencing. Scientific Reports, 10(1): 11227.

Pedersen, B. S. and Quinlan, A. R. 2017. Mosdepth: quick coverage calculation for genomes and exomes. Bioinformatics, 34(5): 867–868.

Pfenninger, M., Reuss, F., Kiebler, A., Schönnenbeck, P., Caliendo, C., Gerber, S., Cocchiararo, B., Reuter, S., Blüthgen, N., Mody, K., Mishra, B., Bálint, M., Thines, M., and Feldmeyer, B. 2021. Genomic basis for drought resistance in European beech forests threatened by climate change. eLife, 10: e65532.

Poyet, M., Le Roux, V., Gibert, P., Meirland, A., Prevost, G., Eslin, P., and Chabrerie, O. 2015. The wide potential trophic niche of the asiatic fruit fly drosophila suzukii: the key of its invasion success in temperate europe? PloS one, 10(11): e0142785.

Prendergast, L. and Reinberg, D. 2021. The missing linker: emerging trends for h1 variant-specific functions. Genes & development, 35(1-2): 40–58.

Rhoné, B., Defrance, D., Berthouly-Salazar, C., Mariac, C., Cubry, P., Couderc, M., Dequincey, A., Assoumanne, A., Kane, N. A., Sultan, B., Barnaud, A., and Vigouroux, Y. 2020. Pearl millet genomic vulnerability to climate change in West Africa highlights the need for regional collaboration. Nature Communications, 11(1): 5274.

Rossi-Stacconi, M. V., Kaur, R., Mazzoni, V., Ometto, L., Grassi, A., Gottardello, A., Rota-Stabelli, O., and Anfora, G. 2016. Multiple lines of evidence for reproductive winter diapause in the invasive pest drosophila suzukii: useful clues for control strategies. Journal of pest science, 89: 689–700.

Ryan, G. D., Emiljanowicz, L., Wilkinson, F., Kornya, M., and Newman, J. A. 2016. Thermal tolerances of the spotted-wing drosophila drosophila suzukii (diptera: Drosophilidae). Journal of Economic Entomology, 109(2): 746–752.

Simberloff, D. 2009. The role of propagule pressure in biological invasions. Annual review of ecology, evolution, and systematics, 40(1): 81–102.

Skoglund, P. and Jakobsson, M. 2011. Archaic human ancestry in east asia. Proceedings of the National Academy of Sciences, 108(45): 18301–18306.

Skotte, L., Korneliussen, T. S., and Albrechtsen, A. 2013. Estimating individual admixture proportions from next generation sequencing data. Genetics, 195(3): 693–702.

Smit, A., Hubley, R., and Green, P. 2023. RepeatMasker Open-4.0. Institute for Systems Biology.

Stockton, D., Wallingford, A., Rendon, D., Fanning, P., Green, C. K., Diepenbrock, L., Ballman, E., Walton, V. M., Isaacs, R., Leach, H., et al. 2019a. Interactions between biotic and abiotic factors affect survival in overwintering drosophila suzukii (diptera: Drosophilidae). Environmental Entomology, 48(2): 454–464.

Stockton, D. G., Brown, R., and Loeb, G. M. 2019b. Not berry hungry? discovering the hidden food sources of a small fruit specialist, drosophila suzukii. Ecological Entomology, 44(6): 810–822.

Thibaud-Nissen, F., Souvorov, A., Murphy, T., DiCuccio, M., and Kitts, P. 2013. Eukaryotic genome annotation pipeline. In The NCBI Handbook. National Center for Biotechnology Information (US). https://www.ncbi.nlm.nih.gov/books/NBK169439/.

Turelli, M., Cooper, B. S., Richardson, K. M., Ginsberg, P. S., Peckenpaugh, B., Antelope, C. X., Kim, K. J., May, M. R., Abrieux, A., Wilson, D. A., Bronski, M. J., Moore, B. R., Gao, J.-J., Eisen, M. B., Chiu, J. C., Conner, W. R., and Hoffmann, A. A. 2018. Rapid global spread of wri-like wolbachia across multiple drosophila. Current Biology, 28(6): 963–971.e8.

Turner, K. G., Ostevik, K. L., Grassa, C. J., and Rieseberg, L. H. 2021. Genomic Analyses of Phenotypic Differences Between Native and Invasive Populations of Diffuse Knapweed (Centaurea diffusa). Frontiers in Ecology and Evolution, 8.

Vasimuddin, M., Misra, S., Li, H., and Aluru, S. 2019. Efficient architecture-aware acceleration of bwa-mem for multicore systems. In 2019 IEEE International Parallel and Distributed Processing Symposium (IPDPS), pages 314–324.

Walsh, D. B., Bolda, M. P., Goodhue, R. E., Dreves, A. J., Lee, J., Bruck, D. J., Walton, V. M., O’Neal, S. D., and Zalom, F. G. 2011. Drosophila suzukii (Diptera: Drosophilidae): Invasive Pest of Ripening Soft Fruit Expanding its Geographic Range and Damage Potential. Journal of Integrated Pest Management, 2(1): G1–G7.

Winkler, A., Jung, J., Kleinhenz, B., and Racca, P. 2020. A review on temperature and humidity effects on Drosophila suzukii population dynamics. Agricultural and Forest Entomology, 22(3): 179–192.

Wood, D. E., Lu, J., and Langmead, B. 2019. Improved metagenomic analysis with kraken 2. Genome Biology, 20(1): 257.

Wu, T., Hu, E., Xu, S., Chen, M., Guo, P., Dai, Z., Feng, T., Zhou, L., Tang, W., Zhan, L., Fu, X., Liu, S., Bo, X., and Yu, G. 2021. clusterProfiler 4.0: A universal enrichment tool for interpreting omics data. The Innovation, 2(3): 100141.

Yang, F., Crossley, M. S., Schrader, L., Dubovskiy, I. M., Wei, S.-J., and Zhang, R. 2022. Polygenic adaptation contributes to the invasive success of the Colorado potato beetle. Molecular Ecology, 31(21): 5568–5580.

