## Supplementary Material for "Adaptive challenges of past and future invasion of *Drosophila suzukii*: insights from novel genomic resources and statistical methods combining individual and pool sequencing data"

#### Supplementary Tables

| ID | Sample Detail | | | | Sequencing | | Realized Coverage | | |
|---|---|---|---|---|---|---|---|---|---|
| | Sampling Area: location | Lat.;Long. | Sampling Date | $n_{ind}$ ($n_{♂}$) | Instrument | SRA_ID (reference) | Auto | X | Y |
| AR-Gol | AME: Argentina (Golondrinas, Chubut) | -42.02;-71.52 | 2023/02 | 100 (100) | NS6000 (PE150) | SRR29639537 (this study) | 53.17 | 27.05 | 14.96 |
| BR-Pal | AME: Brazil (Porto Alegre) | -27.72;-52.17 | 2014/07 | 50 (33) | HS2500 (PE125) | SRR10260033 (Olazcuaga *et al.*, 2020) | 62.78 | 47.37 | 6.54 |
| US-CAOx | AME: USA (Oxnard, California) | 34.15;-119.13 | 2018/11 | 100 (0) | NS6000 (PE150) | SRR24598919 (Feng *et al.*, 2024) | 42.94 | 43.71 | 0.43 |
| US-Col | AME: USA (Fort Collins, Colorado) | 40.57;-105.09 | 2015/09 | 50 (26) | HS2500 (PE125) | SRR10260032 (Olazcuaga *et al.*, 2020) | 66.83 | 52.55 | 6.55 |
| US-Haw | AME: USA (Hilo, Hawaii) | 19.67;-155.47 | 2016/06 | 50 (25) | HS2500 (PE125) | SRR10260031 (Olazcuaga *et al.*, 2020) | 79.46 | 62.12 | 6.27 |
| US-Nca | AME: USA (Raleigh, North Carolina) | 35.70;-80.62 | 2016/10 | 100 (50) | HS2500 (PE125) | SRR10260030 (Olazcuaga *et al.*, 2020) | 61.05 | 48.01 | 5.38 |
| US-Sdi | AME: USA (San-Diego, California) | 32.72;-117.16 | 2014/05 | 50 (32) | HS2500 (PE125) | SRR10260029 (Olazcuaga *et al.*, 2020) | 73.79 | 54.28 | 7.61 |
| US-Sok | AME: USA (Dayton, Oregon) | 45.22;-123.07 | 2014/10 | 75 (55) | HS2500 (PE125) | SRR10260028 (Olazcuaga *et al.*, 2020) | 54.75 | 34.18 | 10.76 |
| US-Wat | AME: USA (Watsonville, California) | 36.90;-121.75 | 2014/10 | 50 (46) | HS2500 (PE125) | SRR10260026 (Olazcuaga *et al.*, 2020) | 59.69 | 33.56 | 10.64 |
| US-Wis | AME: USA (Barneveld, Wisconsin) | 42.97;-89.69 | 2016/11 | 75 (30) | HS2500 (PE125) | SRR10260025 (Olazcuaga *et al.*, 2020) | 65.67 | 52.06 | 6.33 |
| CH-Del | EUR: Switzerland (Delemont) | 47.37;7.35 | 2014/10 | 58 (30) | NS6000 (PE250) | SRR26544765 (this study) | 61.58 | 49.23 | 8.54 |
| DE-Dos | EUR: Germany (Dossenheim) | 49.45;8.66 | 2015/08 | 50 (42) | HS2500 (PE125) | SRR10260021 (Olazcuaga *et al.*, 2020) | 70.37 | 44.90 | 13.44 |
| DE-Jen | EUR: Germany (Jena) | 50.93;11.56 | 2016/09 | 100 (0) | NS6000 (PE250) | SRR29639538 (this study) | 50.26 | 49.60 | 0.92 |
| ES-Bar | EUR: Spain (Barcelona) | 41.36;1.96 | 2014/07 | 50 (50) | HS2500 (PE125) | SRR10260019 (Olazcuaga *et al.*, 2020) | 66.56 | 34.11 | 17.24 |
| FR-Cor | EUR: France (Lozzi, Corsica) | 42.35;9.00 | 2016/08 | 50 (25) | HS2500 (PE125) | SRR10260018 (Olazcuaga *et al.*, 2020) | 60.89 | 51.92 | 4.77 |
| FR-Lez | EUR: France (Montpellier) | 43.70;3.83 | 2014/07 | 100 (50) | HS2500 (PE125) | SRR10260015 (Olazcuaga *et al.*, 2020) | 74.03 | 58.43 | 6.95 |
| FR-Par | EUR: France (Paris) | 48.84;2.36 | 2016/11 | 100 (50) | HS2500 (PE125) | SRR10260014 (Olazcuaga *et al.*, 2020) | 61.33 | 49.24 | 7.45 |
| FR-Run | EUR: France (La Réunion Island) | -21.15;55.64 | 2016/09 | 100 (50) | HS2500 (PE125) | SRR10260013 (Olazcuaga *et al.*, 2020) | 78.15 | 60.83 | 7.02 |
| IT-Tre | EUR: Italy (Trento) | 46.04;11.15 | 2014/09 | 100 (60) | HS2500 (PE125) | SRR10260034 (Olazcuaga *et al.*, 2020) | 60.34 | 44.51 | 9.13 |
| PT-CP21 | EUR: Portugal (Castelo de Paiva) | 41.0;-8.28 | 2021/09 | 80 (40) | NS6000 (PE150) | SRR28145941 (Sario et al., 2024) | 81.90 | 65.06 | 9.14 |
| RS-Zaj | EUR: Serbia (Zajača) | 44.45;19.25 | 2020/09 | 100 (100) | NS6000 (PE250) | SRR26544764 (this study) | 45.80 | 23.31 | 13.32 |
| CN-Bei | NAT: China (Beijing) | 40.00;116.35 | 2014/06 | 50 (11) | HS2500 (PE125) | SRR10260017 (Olazcuaga *et al.*, 2020) | 82.65 | 76.69 | 4.23 |
| CN-Lia | NAT: China (Liaoyuan city, Jilin) | 42.96;125.09 | 2014/08 | 50 (40) | HS2500 (PE125) | SRR10260016 (Olazcuaga *et al.*, 2020) | 59.20 | 37.67 | 9.24 |
| CN-Nin | NAT: China (Ningbo city, Zhejiang) | 30.02;121.5 | 2014/07;09 | 50 (14) | HS2500 (PE125) | SRR10260027 (Olazcuaga *et al.*, 2020) | 56.95 | 52.84 | 2.40 |
| CN-Shi | NAT: China (Shiping county, Yunnan) | 23.70;102.48 | 2014/06;08 | 50 (47) | HS2500 (PE125) | SRR10260024 (Olazcuaga *et al.*, 2020) | 58.56 | 31.76 | 12.08 |
| JP-Kan | NAT: Japan (Kanazawa) | 36.50;136.60 | 2019/10 | 212 (183) | NS6000 (PE150) | SRR24598921 (Feng *et al.*, 2024) | 105.55 | 71.61 | 19.66 |
| JP-Sap | NAT: Japan (Sapporo) | 43.05;141.35 | 2014/07;2016/09 | 50 (46) | HS2500 (PE125) | SRR10260023 (Olazcuaga *et al.*, 2020) | 73.23 | 37.49 | 16.78 |
| JP-Tok | NAT: Japan (Tokyo) | 35.64;139.37 | 2016/06 | 50 (10) | HS2500 (PE125) | SRR10260022 (Olazcuaga *et al.*, 2020) | 57.75 | 51.23 | 3.18 |

Table S1: Description of the Pool-Seq data consisting of 28 *D. suzukii* population. Sampling areas corresponding to the European and American invasion routes are denoted EUR and AME, respectively, and those corresponding to the native area are denoted NAT. Sequencing data consisted of whole-genome shotgun paired-end short reads (e.g., PE150 for pairs of 150 nt reads) generated on Illumina sequencers (NS=NovaSeq and HS=HiSeq). Realized coverages were estimated for autosomes (Auto.) over chr2L, chr2R, chr3 and chr4; the chrX (X) and the chrY (Y) scaffolds of the novel *dsu_isojap1.0* assembly from the read alignment *bam* files using `mosdepth` (v0.3.6, (Pedersen and Quinlan, 2017)) with the option *-Q 20* to ignore reads with low mapping quality. Note that the coverages over Y–linked contigs always remained significantly lower than over X–linked contigs in pools consisting of males only (AR-Gol, ES-Bar and RS-Zaj), which may be mainly due to the high repeat content of the Y combined with the filtering on read mapping quality to estimate coverages (see also Table S2).

| ID | reference | Pop ID | Sample Origin | | | | Realized Coverage | | |
|---|---|---|---|---|---|---|---|---|---|
| | | | Area: Sampling Location | Lat;Lon | Sampling Date | Sex | Auto | X | Y |
| SRR26544762 | this study | CN-Che | NAT: China (Chengdu) | 30.66;104.06 | 2014/08 | F | 13.93 | 13.89 | 0.15 |
| SRR26544761 | | | | | | F | 14.61 | 14.86 | 0.13 |
| SRR26544760 | | | | | | F | 6.71 | 6.77 | 2.39 |
| SRR26544759 | | | | | | F | 12.11 | 11.91 | 0.34 |
| SRR26544758 | | | | | | F | 11.09 | 11.20 | 0.13 |
| SRR26544757 | | | | | | F | 5.83 | 6.01 | 0.08 |
| SRR26544756 | | | | | | F | 11.22 | 11.23 | 0.19 |
| SRR26544755 | | | | | | F | 8.02 | 8.05 | 0.10 |
| SRR26544763 | | | | | | F | 6.57 | 6.22 | 0.36 |
| SRR13964056 | Lewald *et al.* (2021) | IR-Dub | EUR: Ireland (Dublin) | 53.35;-6.27 | 2016/06 | F | 5.39 | 5.49 | 0.05 |
| SRR13964055 | | | | | | F | 8.01 | 8.10 | 0.07 |
| SRR13964054 | | | | | | F | 5.53 | 5.52 | 0.05 |
| SRR13964052 | | | | | | F | 5.35 | 5.44 | 0.06 |
| SRR13964051 | | | | | | F | 6.37 | 6.45 | 0.06 |
| SRR13964050 | | | | | | F | 7.27 | 7.22 | 0.07 |
| SRR13964049 | | | | | | F | 4.43 | 4.40 | 0.04 |
| SRR13964175 | | Ko-San | NAT: South Korea (Sancheong) | 37.57;127.00 | 2016/06 | F | 6.38 | 6.44 | 0.06 |
| SRR13964173 | | | | | | F | 9.30 | 9.25 | 0.09 |
| SRR13964172 | | | | | | F | 8.47 | 8.40 | 0.07 |
| SRR13964171 | | | | | | F | 9.15 | 9.20 | 0.08 |
| SRR13964170 | | | | | | F | 9.23 | 9.16 | 0.08 |
| SRR13964167 | | | | | | F | 1.69 | 1.64 | 0.02 |
| SRR13964166 | | | | | | F | 2.08 | 2.04 | 0.02 |
| SRR13964165 | | | | | | F | 1.79 | 1.79 | 0.03 |
| SRR13964174 | | | | | | F | 3.01 | 2.98 | 0.04 |
| SRR13964157 | | US-Ga | AME: USA (Appling,Georgia) | 31.78;-82.49 | 2016/07;08 | F | 4.90 | 4.76 | 0.05 |
| SRR13964146 | | | | | | F | 2.65 | 2.60 | 0.03 |
| SRR13964135 | | | | | | F | 4.35 | 4.28 | 0.04 |
| SRR13964124 | | | | | | F | 4.13 | 4.05 | 0.04 |
| SRR13964088 | | | AME: USA (Bacon,Georgia) | 31.54;-82.46 | 2016/07;08 | F | 1.47 | 1.48 | 0.01 |
| SRR13964075 | | | | | | M | 1.63 | 1.53 | 0.06 |
| SRR13964064 | | | | | | F | 2.81 | 2.83 | 0.03 |
| SRR13964053 | | | | | | F | 3.28 | 3.28 | 0.03 |
| SRR13964042 | | | | | | F | 2.53 | 2.54 | 0.03 |
| SRR13964249 | | US-Me | AME: USA (Union, Maine) | 44.20;-69.25 | 2016/08 | F | 8.86 | 8.85 | 0.07 |
| SRR13964248 | | | | | | F | 5.15 | 5.08 | 0.07 |
| SRR13964247 | | | | | | F | 8.17 | 8.06 | 0.08 |
| SRR13964245 | | | | | | F | 3.21 | 3.22 | 0.03 |
| SRR13964244 | | | | | | F | 6.29 | 6.15 | 0.06 |
| SRR13964159 | | | | | | F | 3.42 | 3.43 | 0.04 |
| SRR13964156 | | | | | | F | 3.86 | 3.88 | 0.04 |
| SRR13964155 | | | | | | F | 3.89 | 3.78 | 0.05 |
| SRR13964154 | | | | | | F | 3.60 | 3.57 | 0.05 |
| SRR13964153 | | | | | | F | 3.74 | 3.76 | 0.05 |
| SRR13964079 | | US-Mi | AME: USA (Allegan, Michigan) | 42.56;-85.86 | 2016/08 | F | 5.00 | 5.00 | 0.05 |
| SRR13964078 | | | | | | F | 4.71 | 4.70 | 0.04 |
| SRR13964158 | | | | | | F | 5.25 | 5.32 | 0.06 |
| SRR13964110 | | | | | | F | 5.19 | 5.22 | 0.05 |
| SRR13964099 | | | | | | F | 4.57 | 4.63 | 0.05 |
| SRR13964213 | | | | | 2016/07 | F | 5.33 | 5.32 | 0.05 |
| SRR13964202 | | | | | | F | 6.10 | 6.08 | 0.05 |
| SRR13964191 | | | | | | F | 6.94 | 6.89 | 0.07 |
| SRR13964180 | | | | | | M | 8.62 | 4.34 | 1.74 |
| SRR13964169 | | | | | | F | 7.50 | 7.41 | 0.08 |
| SRR13964189 | | US-Ny | AME: USA (Ontario, New York) | 42.86;-77.02 | 2016/08 | F | 4.88 | 4.77 | 0.06 |
| SRR13964188 | | | | | | F | 5.50 | 5.42 | 0.05 |
| SRR13964187 | | | | | | F | 5.33 | 5.27 | 0.05 |
| SRR13964186 | | | | | | F | 4.65 | 4.62 | 0.05 |
| SRR13964185 | | | | | | F | 5.57 | 5.50 | 0.06 |
| SRR13964164 | | | AME: USA (Schuyler, New York) | 42.49;-76.73 | 2016/08 | F | 4.97 | 4.90 | 0.05 |
| SRR13964163 | | | | | | F | 6.03 | 5.92 | 0.06 |
| SRR13964162 | | | | | | F | 7.19 | 7.05 | 0.07 |
| SRR13964161 | | | | | | F | 5.35 | 5.28 | 0.05 |
| SRR13964160 | | | | | | F | 4.70 | 4.62 | 0.05 |
| SRR13964147 | | US-Vir | AME: USA (Virginia Beach, Virginia) | 36.85;-75.98 | 2016/05 | F | 10.69 | 10.48 | 0.09 |
| SRR13964145 | | | | | | F | 7.21 | 7.03 | 0.06 |
| SRR13964144 | | | | | | F | 11.62 | 11.46 | 0.11 |
| SRR13964143 | | | | | | F | 11.80 | 11.44 | 0.11 |
| SRR13964142 | | | | | | F | 12.21 | 11.97 | 0.14 |

| | | | | | | | | | |
|---|---|---|---|---|---|---|---|---|---|
| SRR13964141 | | | | | | M | 12.10 | 6.16 | 2.72 |
| SRR13964140 | | | | | | F | 11.26 | 11.08 | 0.09 |
| SRR13964139 | | | | | | F | 13.66 | 13.29 | 0.11 |
| SRR13964263 | | US-Wa | AME: USA (Little Rock, Washington) | 46.90;-123.02 | 2016/08 | F | 7.75 | 7.76 | 0.10 |
| SRR13964262 | | | | | | M | 6.78 | 3.18 | 2.45 |
| SRR13964261 | | | | | | F | 5.77 | 5.74 | 0.06 |
| SRR13964260 | | | | | | F | 5.88 | 5.84 | 1.36 |
| SRR13964259 | | | | | | M | 6.72 | 3.29 | 2.01 |
| SRR13964116 | | | AME: USA (Olympia, Washington) | 47.03;-122.95 | 2016/08 | F | 9.60 | 9.75 | 0.09 |
| SRR13964115 | | | | | | F | 4.86 | 4.80 | 0.05 |
| SRR13964114 | | | | | | F | 9.12 | 9.08 | 0.07 |
| SRR13964113 | | | | | | F | 8.27 | 8.31 | 0.07 |
| SRR13964112 | | | | | | F | 7.63 | 7.64 | 0.08 |

Table S2: Description of Ind-Seq data consisting of 82 *D. suzukii* individuals sequenced individually and representing nine different populations. Sampling sites corresponding to the European and American invasion routes are denoted EUR and AME, respectively, and those corresponding to the native range are denoted NAT. Note that US-Ga, US-Ny and US-Wa individuals each originated from two adjacent sampling sites (27 km, 45 km and 15 km apart, respectively) that were collected at the same time. Sequencing data consisted of whole-genome shotgun paired-end short reads generated on an Illumina NovaSeq6000 sequencer in PE250 for the 9 newly sequenced CN-Che individuals; and on a HiSeq4000 sequencer in PE150 for the 73 other individuals from (Lewald *et al.*, 2021). Realized coverages were estimated for autosomes (Auto.) over chr2L, chr2R, chr3 and chr4; the chrX (X) and the chrY (Y) scaffolds of the novel *dsu_isojap1.0* assembly from the read alignment *bam* files using `mosdepth` (v0.3.6; Pedersen and Quinlan, 2017) with the option *-Q 20* to ignore reads with low mapping quality. Note that the coverages over Y–linked contigs always remained significantly lower than over X–linked contigs in male individuals, which may be mainly due to the high repeat content of the Y combined with the filtering on read mapping quality to estimate coverages (see also Table S1).

| *Dmel6*<br>*dsu_isojap1.0* | X<br>(23.54 Mb) | 2L<br>(23.51 Mb) | 2R<br>(25.29 Mb) | 3L<br>(28.11 Mb) | 3R<br>(32.08 Mb) | 4<br>(1.348 Mb) | Total |
|---|---|---|---|---|---|---|---|
| X (32.72 Mb) | 506 | 1 | - | 1 | - | - | 508 |
| 2L (26.65 Mb) | - | 541 | 1 | - | - | - | 542 |
| 2R (24.72 Mb) | - | - | 622 | - | - | - | 622 |
| 3 (93.26 Mb) | 1 | - | - | 660 | 804 | - | 1465 |
| 4 (2.560 Mb) | - | - | - | - | - | 13 | 13 |
| Y (13.21 Mb) | - | - | - | - | - | - | - |
| *2c* (40.07 Mb) | - | 8 | 11 | - | - | - | 19 |
| *scf_Xc* (6.316 Mb) | - | - | - | - | - | - | - |
| *scf_Y1* (3.338 Mb) | - | - | - | - | - | - | - |
| other (n=12) | - | 14 | 14 | - | - | - | 28 |
| Total | 507 | 564 | 648 | 661 | 804 | 13 | 3197 |

Table S3: Comparison of BUSCO gene content between the *dsu_isojap1.0* assembly and the *Dmel6* assembly for the *D. melanogaster* genome (Hoskins *et al.*, 2015). In total, 3,197 genes from the `diptera_odb10` dataset that were found to be complete and in single copy in both assemblies. The table provides the number of these genes co-localizing on the chromosomes (or scaffolds) of the two assemblies. Overall, a 99.4% overlap was found for the X chromosome (out of the 509 BUSCO genes mapping to the X chromosomes of either species, 506 were mapped to the X chromosome in both assemblies). Likewise, the overlap was equal to 99.9% and 100% for chromosomes 3 and 4 respectively. The overlap was a bit lower for chromosome arms 2L and 2R (95.8% and 96.0%) respectively, suggesting a less complete sequence. However, an additional large scaffold (40.07 Mb) containing only 19 BUSCO genes, 8 mapping to the 2L and 11 to the 2R *D. melanogaster* chromosome arms, likely corresponded to the (extended) centromeric regions of chromosome 2 (as confirmed by their repeat content, see below). This scaffold was thus named *2c*. Yet, out of the 1,212 (2.31%) BUSCO genes mapped to both *dsu_isojap1.0* and *Dmel6* assemblies and located on the *D. melanogaster* 2L and 2R, 28 were mapped to other unscaffolded contigs of *dsu_isojap1.0*. In contrast, no BUSCO genes from *D. melanogaster* chromosomal arms other than 2L and 2R mapped to unscaffolded contigs of *dsu_isojap1.0*. Conversely, no BUSCO genes were found on the Y (and `Xc`) chromosomes of *dsu_isojap1.0*.

| Chr. | Window position (bp) | Size (bp) | nsnps | Lscore (-log10) | C2 | Gene | Ortholog |
|---|---|---|---|---|---|---|---|
| 2L | 28,410-33,266 | 4856 | 9 | 6.66 (6.24) | $C_2^{EU}$ | *LOC118876810* | - |
| 2L | 95,009-101,846 | 6837 | 54 | 7.33 (3.53) | $C_2^{AM}$ | *dbr* | *dbr* |
| 2L | 608,199-608,947 | 748 | 11 | 8.28 (5.88) | $C_2^{EU}$ | *ush* | *ush* |
| 2L | 814,774-816,200 | 1426 | 13 | 10.12 (5.35) | $C_2^{WW}$ | *ds* | *ds* |
| 2L | 814,774-817,928 | 3154 | 16 | 13.28 (5.41) | $C_2^{EU}$ | *ds* | *ds* |
| 2L | 873,333-873,500 | 167 | 10 | 6.95 (4.92) | $C_2^{EU}$ | *Eaat2* | *Eaat2* |
| 2L | 875,000-875,703 | 703 | 19 | 9.55 (4.69) | $C_2^{EU}$ | *Eaat2* | *Eaat2* |
| 2L | 1,050,346-1,050,912 | 566 | 20 | 15.91 (5.99) | $C_2^{EU}$ | - | - |
| 2L | 1,335,669-1,335,906 | 237 | 10 | 7.3 (4.34) | $C_2^{EU}$ | *Naprt* | *Naprt* |
| 2L | 1,359,257-1,360,428 | 1171 | 14 | 10.21 (6.6) | $C_2^{EU}$ | *LOC108021058* | - |
| 2L | 1,380,287-1,382,356 | 2069 | 25 | 11.61 (4.69) | $C_2^{WW}$ | - | - |
| 2L | 1,380,287-1,382,841 | 2554 | 31 | 19.52 (5.51) | $C_2^{EU}$ | - | - |
| 2L | 1,412,207-1,413,679 | 1472 | 14 | 6.99 (4.11) | $C_2^{EU}$ | - | - |
| 2L | 1,423,404-1,425,307 | 1903 | 37 | 15.23 (6.37) | $C_2^{EU}$ | *LOC108010813* | - |
| 2L | 1,563,301-1,563,788 | 487 | 14 | 8.16 (5.82) | $C_2^{EU}$ | *LOC108011088* | *CG43774* |
| 2L | 1,679,596-1,681,056 | 1460 | 12 | 6.2 (4.07) | $C_2^{EU}$ | *fred* | *fred* |
| 2L | 1,837,663-1,838,181 | 518 | 24 | 6.41 (3.23) | $C_2^{EU}$ | *LOC108011626* | *CG31960* |
| 2L | 1,844,510-1,845,536 | 1026 | 13 | 7.23 (5.41) | $C_2^{EU}$ | *bowl* | *bowl* |
| 2L | 1,979,926-1,980,321 | 395 | 13 | 8.95 (5.93) | $C_2^{EU}$ | *Drgx* | *Drgx* |
| 2L | 2,188,311-2,190,084 | 1773 | 19 | 10.24 (5.08) | $C_2^{EU}$ | *LOC108009123* | *CG4701* |
| 2L | 2,207,327-2,208,491 | 1164 | 32 | 9.36 (3.73) | $C_2^{EU}$ | *LOC108012551* | *CG42586* |
| 2L | 2,237,879-2,239,433 | 1554 | 27 | 11.2 (4.19) | $C_2^{EU}$ | *LOC108012551* | *CG42586* |
| 2L | 2,316,618-2,317,872 | 1254 | 18 | 13.11 (5.45) | $C_2^{EU}$ | *ProtB* | *ProtB* |
| 2L | 2,369,627-2,370,360 | 733 | 9 | 6.37 (5.91) | $C_2^{EU}$ | *side-II* | *side-II* |
| 2L | 2,687,634-2,688,523 | 889 | 14 | 6.88 (4.57) | $C_2^{AM}$ | *LOC108010143* | - |
| 2L | 3,824,831-3,825,054 | 223 | 12 | 6.54 (4.34) | $C_2^{WW}$ | *Slh* | *Slh* |
| 2L | 3,951,649-3,953,187 | 1538 | 13 | 6.2 (4.44) | $C_2^{WW}$ | *Eogt* | *Eogt* |
| 2L | 3,960,124-3,960,797 | 673 | 11 | 7.63 (4.79) | $C_2^{EU}$ | *LOC108021575* | *CG3609* |
| 2L | 4,069,098-4,069,579 | 481 | 11 | 6.19 (4.4) | $C_2^{WW}$ | *LOC108012956* | *CG42658* |
| 2L | 4,178,325-4,179,336 | 1011 | 13 | 8.79 (5.06) | $C_2^{EU}$ | - | - |
| 2L | 4,419,286-4,419,904 | 618 | 10 | 6.58 (5.1) | $C_2^{WW}$ | *witty* | *witty* |
| 2L | 4,688,107-4,689,479 | 1372 | 11 | 8.85 (5.64) | $C_2^{WW}$ | *RpL13* | *RpL13* |
| 2L | 5,394,032-5,394,396 | 364 | 10 | 7.64 (5.85) | $C_2^{WW}$ | *LOC108015628* | *CG31815* |
| 2L | 5,511,310-5,511,703 | 393 | 13 | 9.19 (4.56) | $C_2^{EU}$ | *LOC108020156* | - |
| 2L | 5,688,109-5,689,253 | 1144 | 10 | 8.58 (5.04) | $C_2^{WW}$ | *LOC108021241* | - |
| 2L | 5,706,348-5,707,136 | 788 | 15 | 7.36 (3.96) | $C_2^{WW}$ | *LOC108021190* | *CG34313* |
| 2L | 6,263,978-6,265,153 | 1175 | 11 | 6.37 (4.88) | $C_2^{WW}$ | *Ccdc85* | *Ccdc85* |
| 2L | 6,495,666-6,496,876 | 1210 | 13 | 8.79 (4.9) | $C_2^{WW}$ | *G6P* | *G6P* |
| 2L | 6,813,735-6,815,435 | 1700 | 24 | 16.02 (5.63) | $C_2^{WW}$ | *LOC108021082* | - |
| 2L | 6,813,735-6,814,092 | 357 | 14 | 6.55 (4.4) | $C_2^{AM}$ | *LOC108021082* | - |
| 2L | 6,827,010-6,827,922 | 912 | 16 | 8.01 (4.63) | $C_2^{EU}$ | *LOC139352197* | - |
| 2L | 7,309,565-7,311,164 | 1599 | 42 | 18 (4.46) | $C_2^{EU}$ | *dpy* | *dpy* |
| 2L | 7,314,064-7,314,933 | 869 | 19 | 6.97 (3.43) | $C_2^{EU}$ | *dpy* | *dpy* |
| 2L | 7,742,787-7,743,265 | 478 | 10 | 6.73 (4.36) | $C_2^{WW}$ | *Charon* | *Charon* |
| 2L | 7,742,787-7,743,288 | 501 | 11 | 7.39 (5) | $C_2^{EU}$ | *Charon* | *Charon* |
| 2L | 7,926,376-7,927,135 | 759 | 14 | 8.02 (4.31) | $C_2^{EU}$ | *a5* | *a5* |
| 2L | 10,159,887-10,160,257 | 370 | 14 | 6.39 (3.43) | $C_2^{WW}$ | *Sr-CIV* | *Sr-CIV* |
| 2L | 10,172,886-10,173,768 | 882 | 18 | 8.54 (4.82) | $C_2^{EU}$ | *lace* | *lace* |
| 2L | 10,175,972-10,176,456 | 484 | 10 | 6.9 (4.14) | $C_2^{WW}$ | *lace* | *lace* |
| 2L | 10,576,776-10,577,864 | 1088 | 11 | 7.23 (5.88) | $C_2^{WW}$ | *LOC108010798* | *CG31735* |
| 2L | 11,121,404-11,122,040 | 636 | 10 | 6.64 (4.4) | $C_2^{WW}$ | *LOC108020912* | *CG43861* |
| 2L | 12,073,003-12,073,312 | 309 | 15 | 7.41 (3.69) | $C_2^{EU}$ | *bur* | *bur* |
| 2L | 12,199,279-12,200,397 | 1118 | 28 | 6.58 (3.12) | $C_2^{EU}$ | - | - |
| 2L | 12,564,341-12,565,169 | 828 | 12 | 7.8 (4.88) | $C_2^{WW}$ | *beat-IIIc* | *beat-IIIc* |
| 2L | 12,712,579-12,714,060 | 1481 | 26 | 18.29 (5.6) | $C_2^{WW}$ | *CLIP-190* | *CLIP-190* |
| 2L | 15,723,038-15,723,748 | 710 | 14 | 11.49 (5.22) | $C_2^{WW}$ | *nompB* | *nompB* |
| 2L | 16,048,808-16,050,273 | 1465 | 13 | 6.94 (4.25) | $C_2^{WW}$ | *LOC136116976* | - |
| 2L | 17,791,181-17,791,720 | 539 | 15 | 9.82 (3.98) | $C_2^{WW}$ | *Myo28B1* | *Myo28B1* |
| 2L | 18,390,030-18,391,738 | 1708 | 14 | 7.65 (6.04) | $C_2^{WW}$ | *Eaat1* | *Eaat1* |

| | | | | | | | |
|---|---|---|---|---|---|---|---|
| 2L | 18,916,420-18,917,305 | 885 | 35 | 22.94 (5.64) | $C_2^{WW}$ | *ppk11* | *ppk11* |
| 2L | 18,916,544-18,916,699 | 155 | 16 | 6.61 (3.97) | $C_2^{EU}$ | *ppk11* | *ppk11* |
| 2L | 20,732,806-20,734,671 | 1865 | 20 | 11.32 (4.26) | $C_2^{EU}$ | *LOC108021564* | - |
| 2L | 23,649,472-23,651,201 | 1729 | 16 | 7.58 (3.7) | $C_2^{EU}$ | - | - |
| 2L | 23,710,750-23,711,683 | 933 | 11 | 8.68 (5.46) | $C_2^{EU}$ | *LOC108011978* | - |
| 2L | 25,715,620-26,155,981 | 440361 | 120 | 88.89 (10.12) | $C_2^{EU}$ | *LOC139352211* | - |
| 2L | 25,716,108-26,192,930 | 476822 | 139 | 56.62 (5.81) | $C_2^{WW}$ | *LOC139352211* | - |
| 2L | 25,716,215-26,072,519 | 356304 | 49 | 22.24 (4.16) | $C_2^{AM}$ | *LOC139352211* | - |
| 2L | 26,294,113-26,344,502 | 50389 | 20 | 7.28 (3.86) | $C_2^{WW}$ | - | - |
| 2L | 26,294,113-26,320,695 | 26582 | 13 | 7.77 (3.71) | $C_2^{EU}$ | - | - |

Table S4: Description of all the significant windows for the chromosome 2R (n=69), with annotated positional candidate genes. The table gives for each significant window (i) its position in the new *dsu_isojap1.0* assembly of the *D. suzukii* genome; (ii) its size in bp; (iii) its number of SNPs; (iv) the window highest maximum Lindley score (Lscore) and its most significant $C_2$ P-values (in $-\log_{10}$ scale); (v) the $C_2$ statistic corresponding to this highest Lscore; (vi) the positional candidate gene associated and (vii) its corresponding *D. melanogaster* ortholog.

| Chr. | Position in bp | Size in bp | nsnps | Lscore ($-\log_{10}$ P$_{C2}$) | $C_2$ | Gene | Ortholog |
|---|---|---|---|---|---|---|---|
| 2R | 2,107,984-2,109,383 | 1399 | 28 | 13.29 (4.72) | $C_2^{WW}$ | *Cyp6w1* | *Cyp6w1* |
| 2R | 2,142,612-2,143,598 | 986 | 24 | 14.27 (5.02) | $C_2^{WW}$ | *Ptr* | *Ptr* |
| 2R | 2,142,653-2,143,238 | 585 | 13 | 6.66 (4.09) | $C_2^{EU}$ | *Ptr* | *Ptr* |
| 2R | 3,798,668-3,799,613 | 945 | 22 | 9.79 (4.07) | $C_2^{WW}$ | *Dic3* | *Dic3* |
| 2R | 3,798,693-3,799,603 | 910 | 18 | 9.99 (4.14) | $C_2^{EU}$ | *Dic3* | *Dic3* |
| 2R | 5,500,268-5,509,155 | 8887 | 18 | 10.61 (4.56) | $C_2^{WW}$ | *LOC108018112* | *CG15107* |
| 2R | 6,120,038-6,120,440 | 402 | 13 | 8.45 (4.42) | $C_2^{EU}$ | *Strn-Mlck* | *Strn-Mlck* |
| 2R | 6,120,090-6,120,379 | 289 | 11 | 8.63 (4.86) | $C_2^{WW}$ | *Strn-Mlck* | *Strn-Mlck* |
| 2R | 6,288,199-6,290,157 | 1958 | 20 | 10.44 (3.96) | $C_2^{WW}$ | *sbb* | *sbb* |
| 2R | 6,626,895-6,628,109 | 1214 | 15 | 11.76 (5.52) | $C_2^{EU}$ | *GEFmeso* | *GEFmeso* |
| 2R | 6,774,982-6,775,936 | 954 | 21 | 6.46 (4.17) | $C_2^{WW}$ | *Vps13* | *Vps13* |
| 2R | 6,776,794-6,777,618 | 824 | 23 | 8.16 (3.38) | $C_2^{WW}$ | *Vps13* | *Vps13* |
| 2R | 6,832,900-6,833,533 | 633 | 11 | 7.39 (4.54) | $C_2^{WW}$ | *LOC108018122* | *CG1598* |
| 2R | 7,365,581-7,366,417 | 836 | 23 | 12.1 (4.14) | $C_2^{WW}$ | *Tsp42El* | *Tsp42El* |
| 2R | 7,370,703-7,372,947 | 2244 | 38 | 18.53 (5.81) | $C_2^{WW}$ | *LOC108017765* | - |
| 2R | 7,371,039-7,372,614 | 1575 | 27 | 10.34 (4.33) | $C_2^{EU}$ | *LOC108017765* | - |
| 2R | 7,371,789-7,372,280 | 491 | 13 | 7.06 (4.26) | $C_2^{AM}$ | *LOC108017765* | - |
| 2R | 7,412,815-7,413,168 | 353 | 11 | 7.77 (4.68) | $C_2^{EU}$ | *LOC108008867* | *CG30016* |
| 2R | 7,414,353-7,423,226 | 8873 | 112 | 34.6 (5.84) | $C_2^{EU}$ | *LOC108009451* | *CG30015* |
| 2R | 7,431,356-7,435,038 | 3682 | 51 | 15.16 (4.26) | $C_2^{EU}$ | *LOC108009451* | *CG30015* |
| 2R | 7,436,051-7,437,830 | 1779 | 18 | 8.95 (4.89) | $C_2^{EU}$ | *LOC108009451* | *CG30015* |
| 2R | 7,441,162-7,451,978 | 10816 | 145 | 23.53 (5.3) | $C_2^{EU}$ | *LOC108009451* | *CG30015* |
| 2R | 7,441,385-7,448,210 | 6825 | 84 | 16.14 (5.59) | $C_2^{WW}$ | *LOC108009451* | *CG30015* |
| 2R | 7,453,238-7,453,527 | 289 | 9 | 6.41 (4.56) | $C_2^{EU}$ | *LOC108009451* | *CG30015* |
| 2R | 7,696,603-7,701,667 | 5064 | 31 | 16.32 (5.09) | $C_2^{EU}$ | *psq* | *psq* |
| 2R | 7,696,603-7,701,667 | 5064 | 31 | 16.32 (5.09) | $C_2^{EU}$ | *LOC136116670* | - |
| 2R | 7,917,791-7,918,880 | 1089 | 12 | 7.08 (4.71) | $C_2^{WW}$ | *Prx6c* | *Prx6c* |
| 2R | 7,917,791-7,918,985 | 1194 | 13 | 8.15 (5.32) | $C_2^{EU}$ | *LOC108009870* | *CG11825* |
| 2R | 8,106,820-8,108,418 | 1598 | 17 | 7.09 (3.79) | $C_2^{WW}$ | *LOC108009011* | *CG12910* |
| 2R | 8,219,851-8,223,669 | 3818 | 30 | 7.72 (5.06) | $C_2^{WW}$ | *KCNQ* | *KCNQ* |
| 2R | 8,219,851-8,222,053 | 2202 | 21 | 9.15 (4.41) | $C_2^{EU}$ | *KCNQ* | *KCNQ* |
| 2R | 8,293,842-8,295,314 | 1472 | 18 | 7.41 (4.76) | $C_2^{EU}$ | *LOC108009803* | *CG2269* |
| 2R | 8,297,632-8,301,183 | 3551 | 29 | 6.96 (4.77) | $C_2^{WW}$ | *LOC108009803* | *CG2269* |
| 2R | 8,921,852-8,924,115 | 2263 | 34 | 11.73 (5.25) | $C_2^{EU}$ | *LOC108009002* | *CG13438* |
| 2R | 8,937,103-8,939,349 | 2246 | 28 | 13.33 (5.33) | $C_2^{EU}$ | *Gr57a* | *Gr57a* |
| 2R | 8,958,439-8,959,264 | 825 | 34 | 8.09 (4.16) | $C_2^{WW}$ | *LOC108008181* | - |
| 2R | 8,961,800-8,964,052 | 2252 | 53 | 14.88 (5.26) | $C_2^{WW}$ | *Smyd4-3* | *Smyd4-3* |
| 2R | 9,233,752-9,235,457 | 1705 | 19 | 11.79 (5.63) | $C_2^{EU}$ | *LOC108010005* | *CG13229* |
| 2R | 9,691,283-9,692,013 | 730 | 9 | 7.07 (4.53) | $C_2^{EU}$ | *Trnah-gug* | - |
| 2R | 9,821,111-9,822,208 | 1097 | 17 | 7.51 (3.91) | $C_2^{WW}$ | *Sr-CII* | *Sr-CII* |
| 2R | 9,839,236-9,840,436 | 1200 | 21 | 6.46 (4.69) | $C_2^{EU}$ | *LOC108008443* | - |
| 2R | 9,875,018-9,875,624 | 606 | 10 | 7.35 (5.67) | $C_2^{EU}$ | *Pcf11* | *Pcf11* |
| 2R | 10,307,367-10,309,308 | 1941 | 25 | 6.93 (4.22) | $C_2^{EU}$ | *Cpsf160* | *Cpsf160* |
| 2R | 10,312,263-10,314,478 | 2215 | 29 | 7.69 (4.29) | $C_2^{EU}$ | *PRAS40* | *PRAS40* |
| 2R | 10,324,833-10,329,512 | 4679 | 30 | 8.69 (4.09) | $C_2^{EU}$ | *PRAS40* | *PRAS40* |
| 2R | 10,378,629-10,378,953 | 324 | 9 | 7.3 (5.77) | $C_2^{EU}$ | *Achl* | *Achl* |
| 2R | 10,402,741-10,406,402 | 3661 | 21 | 7.06 (5.59) | $C_2^{WW}$ | - | - |
| 2R | 10,522,521-10,523,243 | 722 | 15 | 7.32 (6.53) | $C_2^{WW}$ | *Ih* | *Ih* |
| 2R | 10,782,486-10,783,728 | 1242 | 14 | 9.61 (5.97) | $C_2^{EU}$ | *mam* | *mam* |
| 2R | 11,308,192-11,309,132 | 940 | 13 | 7.55 (5.56) | $C_2^{EU}$ | *LOC139352563* | - |
| 2R | 11,547,397-11,548,573 | 1176 | 8 | 6.47 (5.38) | $C_2^{WW}$ | *reb* | *reb* |
| 2R | 11,708,384-11,710,108 | 1724 | 15 | 8.47 (5.71) | $C_2^{EU}$ | *LOC108008906* | *CG30039* |
| 2R | 12,136,254-12,136,902 | 648 | 12 | 9.48 (4.93) | $C_2^{EU}$ | *LOC108008703* | *CG13423* |
| 2R | 12,293,409-12,295,176 | 1767 | 13 | 6.52 (3.69) | $C_2^{WW}$ | - | - |
| 2R | 12,819,188-12,821,486 | 2298 | 19 | 8 (4.81) | $C_2^{EU}$ | *LOC108018783* | *CG8654* |
| 2R | 12,859,018-12,859,321 | 303 | 16 | 8.55 (3.66) | $C_2^{WW}$ | *LOC108018780* | *CG43195* |
| 2R | 12,928,537-12,929,306 | 769 | 16 | 9.26 (5.37) | $C_2^{WW}$ | *Obp56h* | *Obp56h* |
| 2R | 12,931,873-12,933,825 | 1952 | 27 | 13.6 (5.73) | $C_2^{WW}$ | *Obp56h* | *Obp56h* |

| | | | | | | | |
|---|---|---|---|---|---|---|---|
| 2R | 13,225,212-13,225,626 | 414 | 8 | 7.03 (5.24) | $C_2^{WW}$ | *LOC108008886* | *CG15124* |
| 2R | 13,380,549-13,382,736 | 2187 | 30 | 6.34 (5.38) | $C_2^{WW}$ | *LOC108008462* | *CG16926* |
| 2R | 14,110,849-14,111,974 | 1125 | 18 | 7 (4.66) | $C_2^{WW}$ | *LOC108009708* | *CG4294* |
| 2R | 14,110,849-14,112,472 | 1623 | 25 | 14.09 (5.77) | $C_2^{EU}$ | *LOC108009708* | *CG4294* |
| 2R | 14,117,322-14,117,602 | 280 | 12 | 8.06 (4.17) | $C_2^{EU}$ | *Vps20* | *Vps20* |
| 2R | 14,268,941-14,270,639 | 1698 | 32 | 7.48 (4.19) | $C_2^{EU}$ | *LOC108010233* | *CG30268* |
| 2R | 14,505,916-14,506,937 | 1021 | 22 | 9.23 (3.8) | $C_2^{EU}$ | *Alg3* | *Alg3* |
| 2R | 14,540,951-14,544,110 | 3159 | 27 | 10.37 (4.24) | $C_2^{EU}$ | *egl* | *egl* |
| 2R | 14,559,329-14,559,961 | 632 | 12 | 7.2 (5.48) | $C_2^{WW}$ | *LOC108008125* | - |
| 2R | 14,562,141-14,562,931 | 790 | 19 | 6.82 (4.55) | $C_2^{WW}$ | *LOC118876909* | - |
| 2R | 14,602,876-14,604,202 | 1326 | 12 | 7.61 (5.62) | $C_2^{WW}$ | *sona* | *sona* |
| 2R | 14,817,419-14,817,773 | 354 | 21 | 6.78 (3.46) | $C_2^{WW}$ | *LOC108010306* | *CG8089* |
| 2R | 14,844,223-14,845,039 | 816 | 15 | 10.28 (5.62) | $C_2^{EU}$ | *chn* | *chn* |
| 2R | 14,935,288-14,937,792 | 2504 | 14 | 7.76 (5) | $C_2^{EU}$ | *hbs* | *hbs* |
| 2R | 15,341,641-15,342,441 | 800 | 22 | 12.86 (5.68) | $C_2^{EU}$ | *Sara* | *Sara* |
| 2R | 15,505,848-15,506,652 | 804 | 16 | 11.69 (4.71) | $C_2^{WW}$ | *LOC108010319* | *CG15673* |
| 2R | 15,505,848-15,507,003 | 1155 | 20 | 14.33 (5.45) | $C_2^{EU}$ | *LOC108010319* | *CG15673* |
| 2R | 16,254,443-16,255,885 | 1442 | 24 | 7.57 (3.43) | $C_2^{EU}$ | *Pms2* | *Pms2* |
| 2R | 16,279,591-16,282,779 | 3188 | 36 | 14.47 (6.55) | $C_2^{EU}$ | - | - |
| 2R | 16,285,979-16,291,308 | 5329 | 33 | 7.15 (4.35) | $C_2^{EU}$ | - | - |
| 2R | 17,385,602-17,395,255 | 9653 | 85 | 15.29 (4.74) | $C_2^{EU}$ | *LOC108018145* | *CG34215* |
| 2R | 17,405,228-17,409,398 | 4170 | 55 | 20.99 (3.99) | $C_2^{EU}$ | *LOC108018145* | *CG34215* |
| 2R | 17,407,367-17,409,807 | 2440 | 56 | 21.49 (4.63) | $C_2^{WW}$ | *LOC108018145* | *CG34215* |
| 2R | 17,422,192-17,422,654 | 462 | 12 | 6.33 (4.65) | $C_2^{EU}$ | *LOC108017707* | *CG10474* |
| 2R | 17,429,292-17,430,223 | 931 | 19 | 6.94 (4.42) | $C_2^{EU}$ | *coro* | *coro* |
| 2R | 17,520,068-17,524,082 | 4014 | 80 | 20.09 (5.74) | $C_2^{EU}$ | *CheB42c* | *CheB42c* |
| 2R | 17,560,199-17,562,822 | 2623 | 32 | 9.68 (4.98) | $C_2^{EU}$ | *LOC108017744* | *CG30156* |
| 2R | 17,572,794-17,598,204 | 25410 | 55 | 11.11 (4.97) | $C_2^{EU}$ | *Taf5* | *Taf5* |
| 2R | 17,572,968-17,574,278 | 1310 | 16 | 6.38 (4.41) | $C_2^{WW}$ | *Tsp42Ea* | *Tsp42Ea* |
| 2R | 17,572,968-17,596,298 | 23330 | 202 | 26.67 (6.46) | $C_2^{EU}$ | *Tsp42Ec* | *Tsp42Ec* |
| 2R | 18,811,937-18,813,450 | 1513 | 12 | 7.15 (5.15) | $C_2^{EU}$ | *LOC108010050* | *CG13579* |
| 2R | 19,513,287-19,513,599 | 312 | 15 | 6.57 (3.74) | $C_2^{EU}$ | *fab1* | *fab1* |
| 2R | 20,199,358-20,200,926 | 1568 | 11 | 7.95 (4.79) | $C_2^{EU}$ | *SP2353* | *SP2353* |
| 2R | 20,370,968-20,372,060 | 1092 | 14 | 7.65 (4.02) | $C_2^{WW}$ | *LOC108010310* | *CG6262* |
| 2R | 20,372,624-20,376,078 | 3454 | 31 | 7.77 (5.47) | $C_2^{EU}$ | *LOC139352496* | - |
| 2R | 20,515,990-20,517,824 | 1834 | 23 | 9.61 (4.39) | $C_2^{EU}$ | - | - |
| 2R | 20,804,646-20,805,565 | 919 | 18 | 11.8 (5.6) | $C_2^{WW}$ | *mute* | *mute* |
| 2R | 20,804,646-20,805,376 | 730 | 12 | 6.9 (4.15) | $C_2^{EU}$ | *mute* | *mute* |
| 2R | 21,658,573-21,661,462 | 2889 | 18 | 8.91 (4.14) | $C_2^{WW}$ | *LOC108009354* | *CG13954* |
| 2R | 21,913,039-21,914,886 | 1847 | 19 | 9.06 (3.8) | $C_2^{EU}$ | *Mmp2* | *Mmp2* |
| 2R | 22,219,828-22,221,051 | 1223 | 17 | 7.77 (4.65) | $C_2^{WW}$ | *Gr59c* | *Gr59c* |
| 2R | 22,296,061-22,297,755 | 1694 | 23 | 11.47 (4.92) | $C_2^{WW}$ | *LOC108008935* | *CG9877* |
| 2R | 22,296,061-22,297,645 | 1584 | 21 | 7.94 (4.75) | $C_2^{EU}$ | *LOC108008935* | *CG9877* |
| 2R | 22,611,155-22,612,754 | 1599 | 15 | 6.63 (3.86) | $C_2^{WW}$ | *stum* | *stum* |
| 2R | 22,611,155-22,614,150 | 2995 | 26 | 20.06 (5.14) | $C_2^{EU}$ | *stum* | *stum* |
| 2R | 22,788,361-22,789,678 | 1317 | 10 | 7.03 (4.94) | $C_2^{EU}$ | *LOC108009650* | - |
| 2R | 22,792,797-22,807,161 | 14364 | 21 | 20.68 (8.47) | $C_2^{WW}$ | *LOC118876994* | - |
| 2R | 22,792,797-22,807,211 | 14414 | 22 | 19.08 (8.76) | $C_2^{EU}$ | *LOC118876994* | - |
| 2R | 23,835,642-23,836,806 | 1164 | 27 | 7.41 (3.42) | $C_2^{WW}$ | *pain* | *pain* |
| 2R | 23,835,642-23,836,548 | 906 | 22 | 6.43 (3.34) | $C_2^{EU}$ | *pain* | *pain* |
| 2R | 23,855,924-23,858,526 | 2602 | 37 | 16.43 (4.78) | $C_2^{WW}$ | *LOC108009111* | *CG15861* |

Table S5: Description of all the significant windows for the chromosome 2R (n=108), with annotated positional candidate genes. The table gives for each significant window (i) its position in the new *dsu_isojap1.0* assembly of the *D. suzukii* genome; (ii) its size in bp; (iii) its number of SNPs; (iv) the window highest maximum Lindley score (Lscore) and its most significant $C_2$ P-values (in $-\log_{10}$ scale); (v) the $C_2$ statistic corresponding to this highest Lscore; (vi) the positional candidate gene associated and (vii) its corresponding *D. melanogaster* ortholog.

| Chr. | Position in bp | Size in bp | nsnps | Lscore ($-\log_{10} \mathrm{P}_{C2}$) | $C_2$ | Gene | Ortholog |
|---|---|---|---|---|---|---|---|
| 3 | 360,977-365,403 | 4426 | 14 | 9.62 (5.38) | $C_2^{EU}$ | *Lsp1gamma* | *Lsp1gamma* |
| 3 | 886,223-886,898 | 675 | 10 | 7.53 (6.13) | $C_2^{WW}$ | - | - |
| 3 | 1,010,570-1,012,828 | 2258 | 16 | 11.68 (5.95) | $C_2^{EU}$ | *LOC136116930* | - |
| 3 | 1,010,935-1,012,141 | 1206 | 13 | 10.33 (5.86) | $C_2^{WW}$ | *LOC136116930* | - |
| 3 | 1,430,701-1,432,307 | 1606 | 13 | 7.18 (4.03) | $C_2^{EU}$ | *mwh* | *mwh* |
| 3 | 1,971,582-1,972,835 | 1253 | 13 | 7.68 (5.51) | $C_2^{EU}$ | *LOC108013501* | *CG7991* |
| 3 | 2,233,648-2,234,068 | 420 | 19 | 9.81 (4.66) | $C_2^{WW}$ | *Mfap1* | *Mfap1* |
| 3 | 2,233,648-2,234,068 | 420 | 19 | 11.73 (5.43) | $C_2^{EU}$ | *Mfap1* | *Mfap1* |
| 3 | 2,689,981-2,691,928 | 1947 | 12 | 10.76 (5.78) | $C_2^{EU}$ | *MsR1* | *MsR1* |
| 3 | 4,143,342-4,146,450 | 3108 | 14 | 7.11 (4.95) | $C_2^{EU}$ | *siz* | *siz* |
| 3 | 4,486,627-4,489,879 | 3252 | 19 | 12.54 (4.8) | $C_2^{WW}$ | *rgn* | *rgn* |
| 3 | 5,302,737-5,304,268 | 1531 | 14 | 7.82 (4.4) | $C_2^{EU}$ | *Or65a* | *Or65a* |
| 3 | 5,433,657-5,433,922 | 265 | 13 | 8.08 (4.28) | $C_2^{EU}$ | *LOC108012428* | *CG7386* |
| 3 | 5,495,947-5,499,268 | 3321 | 33 | 20.19 (5.1) | $C_2^{EU}$ | *LOC108012509* | - |
| 3 | 5,496,000-5,498,772 | 2772 | 22 | 11.82 (4.52) | $C_2^{WW}$ | *LOC108012509* | - |
| 3 | 5,540,913-5,542,743 | 1830 | 13 | 7.27 (4.72) | $C_2^{EU}$ | *LOC108013595* | - |
| 3 | 5,820,390-5,821,845 | 1455 | 14 | 7.22 (4.39) | $C_2^{EU}$ | *Ak1* | *Ak1* |
| 3 | 5,864,972-5,866,752 | 1780 | 14 | 7.32 (4.28) | $C_2^{EU}$ | *LOC108012919* | - |
| 3 | 5,961,387-5,965,296 | 3909 | 22 | 9.09 (4.91) | $C_2^{EU}$ | *Lmx1a* | *Lmx1a* |
| 3 | 6,072,355-6,073,855 | 1500 | 18 | 7.4 (5.14) | $C_2^{EU}$ | *toe* | *toe* |
| 3 | 6,211,391-6,211,838 | 447 | 13 | 7.15 (6.01) | $C_2^{EU}$ | *Sytbeta* | *Sytbeta* |
| 3 | 8,013,117-8,013,768 | 651 | 15 | 7.92 (5.68) | $C_2^{EU}$ | *Rbp6* | *Rbp6* |
| 3 | 8,088,552-8,089,617 | 1065 | 14 | 8.46 (4.22) | $C_2^{WW}$ | - | - |
| 3 | 8,757,572-8,758,900 | 1328 | 34 | 10.62 (5.75) | $C_2^{EU}$ | *Rbfox1* | *Rbfox1* |
| 3 | 11,266,222-11,266,636 | 414 | 10 | 8.11 (5.36) | $C_2^{WW}$ | *Atg18a* | *Atg18a* |
| 3 | 11,532,016-11,532,415 | 399 | 14 | 10.48 (5.61) | $C_2^{EU}$ | *stv* | *stv* |
| 3 | 12,156,547-12,159,913 | 3366 | 29 | 9.89 (4.54) | $C_2^{EU}$ | - | - |
| 3 | 12,209,850-12,213,634 | 3784 | 28 | 13.23 (5.47) | $C_2^{EU}$ | *Con* | *Con* |
| 3 | 12,211,541-12,213,099 | 1558 | 10 | 7.47 (4.62) | $C_2^{WW}$ | *Con* | *Con* |
| 3 | 12,772,552-12,773,767 | 1215 | 25 | 9.77 (5.2) | $C_2^{EU}$ | *promL* | *promL* |
| 3 | 12,975,254-12,976,245 | 991 | 35 | 12.71 (4.86) | $C_2^{EU}$ | *LOC108007079* | *CG12009* |
| 3 | 13,064,257-13,065,521 | 1264 | 22 | 11.53 (4.55) | $C_2^{EU}$ | *kst* | *kst* |
| 3 | 13,064,318-13,065,353 | 1035 | 16 | 12.78 (5.17) | $C_2^{WW}$ | *kst* | *kst* |
| 3 | 13,747,593-13,748,047 | 454 | 10 | 7.54 (5.07) | $C_2^{WW}$ | *LOC108011838* | *CG12766* |
| 3 | 13,747,593-13,748,131 | 538 | 12 | 7.43 (4.81) | $C_2^{EU}$ | *LOC108011838* | *CG12766* |
| 3 | 14,160,044-14,163,040 | 2996 | 17 | 8.99 (4.49) | $C_2^{AM}$ | *LOC108020039* | - |
| 3 | 16,193,344-16,194,399 | 1055 | 11 | 7.58 (5.46) | $C_2^{WW}$ | *nudC* | *nudC* |
| 3 | 17,183,154-17,184,657 | 1503 | 14 | 7.59 (4.97) | $C_2^{WW}$ | *Grip163* | *Grip163* |
| 3 | 17,190,751-17,191,644 | 893 | 15 | 8.29 (3.78) | $C_2^{WW}$ | *Pop2* | *Pop2* |
| 3 | 17,195,783-17,196,370 | 587 | 17 | 12.02 (6.03) | $C_2^{EU}$ | *LOC108014665* | *CG6928* |
| 3 | 17,827,007-17,827,998 | 991 | 16 | 9.98 (4.11) | $C_2^{EU}$ | *LOC108006195* | - |
| 3 | 21,381,090-21,381,978 | 888 | 18 | 7.59 (4.34) | $C_2^{WW}$ | *Pi3K68D* | *Pi3K68D* |
| 3 | 21,580,711-21,581,596 | 885 | 23 | 11.63 (3.68) | $C_2^{EU}$ | *Brr2* | *Brr2* |
| 3 | 22,120,425-22,123,142 | 2717 | 9 | 7.14 (5.13) | $C_2^{WW}$ | *Ran-like* | *Ran-like* |
| 3 | 23,130,492-23,132,612 | 2120 | 20 | 16.68 (5.98) | $C_2^{WW}$ | *RpL26* | *RpL26* |
| 3 | 23,130,492-23,132,229 | 1737 | 17 | 12.63 (5.05) | $C_2^{EU}$ | *RpL26* | *RpL26* |
| 3 | 24,025,960-24,026,367 | 407 | 18 | 7.98 (4.23) | $C_2^{WW}$ | *Cpr76Bc* | *Cpr76Bc* |
| 3 | 24,154,424-24,155,648 | 1224 | 29 | 9.74 (3.63) | $C_2^{WW}$ | *wnd* | *wnd* |
| 3 | 25,214,793-25,215,059 | 266 | 14 | 9.2 (4.32) | $C_2^{WW}$ | *olf413* | *olf413* |
| 3 | 25,214,793-25,215,283 | 490 | 26 | 23.9 (7.71) | $C_2^{EU}$ | *olf413* | *olf413* |
| 3 | 25,290,062-25,292,344 | 2282 | 21 | 8.23 (4.86) | $C_2^{EU}$ | *Ten-m* | *Ten-m* |
| 3 | 31,670,526-31,680,129 | 9603 | 28 | 8.82 (4.22) | $C_2^{WW}$ | *LOC139352732* | *CG40470* |
| 3 | 31,916,881-31,947,817 | 30936 | 22 | 14.13 (3.72) | $C_2^{EU}$ | *LOC139352732* | *CG40470* |
| 3 | 35,403,601-35,413,200 | 9599 | 14 | 9.74 (4.96) | $C_2^{WW}$ | *nvd* | *nvd* |
| 3 | 35,403,609-35,412,503 | 8894 | 10 | 8.13 (4.99) | $C_2^{EU}$ | *nvd* | *nvd* |
| 3 | 35,840,733-35,854,980 | 14247 | 20 | 9.48 (4.33) | $C_2^{WW}$ | - | - |
| 3 | 38,412,467-38,422,123 | 9656 | 13 | 11.85 (5.62) | $C_2^{WW}$ | *LOC108020539* | - |
| 3 | 38,412,467-38,420,880 | 8413 | 12 | 10.23 (5.31) | $C_2^{EU}$ | *LOC108020539* | - |

| | | | | | | | |
|---|---|---|---|---|---|---|---|
| 3 | 39,644,847-39,673,166 | 28319 | 18 | 9.43 (3.88) | $C_2^{WW}$ | *MED21* | *MED21* |
| 3 | 41,170,546-41,195,093 | 24547 | 31 | 19.93 (5.14) | $C_2^{WW}$ | - | - |
| 3 | 43,836,675-43,842,731 | 6056 | 11 | 7.19 (7.82) | $C_2^{WW}$ | - | - |
| 3 | 46,847,315-46,877,930 | 30615 | 29 | 13.71 (5.54) | $C_2^{EU}$ | - | - |
| 3 | 47,044,857-47,046,392 | 1535 | 16 | 16.21 (5.96) | $C_2^{WW}$ | - | - |
| 3 | 47,044,857-47,062,008 | 17151 | 41 | 20.82 (7.19) | $C_2^{EU}$ | - | - |
| 3 | 47,069,550-47,072,304 | 2754 | 21 | 17.32 (7.05) | $C_2^{WW}$ | - | - |
| 3 | 47,069,550-47,072,569 | 3019 | 22 | 18.82 (7.62) | $C_2^{EU}$ | - | - |
| 3 | 47,152,278-47,155,183 | 2905 | 17 | 9.24 (4.57) | $C_2^{EU}$ | *LOC139352805* | - |
| 3 | 47,197,814-47,199,600 | 1786 | 15 | 7.53 (4.43) | $C_2^{WW}$ | *LOC139352805* | - |
| 3 | 48,966,347-48,989,605 | 23258 | 25 | 17.97 (4.65) | $C_2^{WW}$ | *Myo81F* | *Myo81F* |
| 3 | 48,966,347-48,992,700 | 26353 | 37 | 38 (6.48) | $C_2^{EU}$ | *Myo81F* | *Myo81F* |
| 3 | 50,334,828-50,346,059 | 11231 | 19 | 11.01 (4.22) | $C_2^{WW}$ | *Myo81F* | *Myo81F* |
| 3 | 50,866,233-50,868,278 | 2045 | 10 | 7.03 (5.57) | $C_2^{WW}$ | *Myo81F* | *Myo81F* |
| 3 | 50,866,233-50,868,416 | 2183 | 13 | 11.49 (6.88) | $C_2^{EU}$ | *Myo81F* | *Myo81F* |
| 3 | 52,686,029-52,687,628 | 1599 | 14 | 9.47 (4.12) | $C_2^{WW}$ | - | - |
| 3 | 53,137,144-53,138,589 | 1445 | 11 | 9.46 (4.95) | $C_2^{WW}$ | - | - |
| 3 | 53,137,144-53,138,631 | 1487 | 14 | 15.67 (7.25) | $C_2^{EU}$ | - | - |
| 3 | 54,485,924-54,491,774 | 5850 | 19 | 14.45 (4.11) | $C_2^{WW}$ | - | - |
| 3 | 56,797,825-56,798,212 | 387 | 13 | 10.18 (5.34) | $C_2^{WW}$ | - | - |
| 3 | 58,688,930-58,699,535 | 10605 | 41 | 35.87 (5.4) | $C_2^{EU}$ | - | - |
| 3 | 58,689,109-58,696,252 | 7143 | 26 | 15.82 (4.42) | $C_2^{WW}$ | - | - |
| 3 | 61,495,131-61,495,984 | 853 | 11 | 8.09 (5.21) | $C_2^{EU}$ | - | - |
| 3 | 62,039,104-62,045,236 | 6132 | 8 | 7.81 (6.81) | $C_2^{WW}$ | - | - |
| 3 | 62,196,558-62,197,745 | 1187 | 26 | 16.4 (5.32) | $C_2^{WW}$ | *smash* | *smash* |
| 3 | 63,351,158-63,353,657 | 2499 | 46 | 15.62 (4.06) | $C_2^{WW}$ | *LOC108006723* | *CG31551* |
| 3 | 63,400,566-63,401,196 | 630 | 13 | 7.32 (5.31) | $C_2^{WW}$ | *Wdr33* | *Wdr33* |
| 3 | 63,504,772-63,505,527 | 755 | 17 | 13.48 (4.33) | $C_2^{WW}$ | *LOC108004869* | *CG2082* |
| 3 | 63,653,625-63,654,722 | 1097 | 10 | 8.04 (5.82) | $C_2^{EU}$ | *side-III* | *side-III* |
| 3 | 63,808,355-63,808,828 | 473 | 13 | 8 (4.54) | $C_2^{WW}$ | *LOC108014344* | *CG10280* |
| 3 | 63,879,134-63,879,769 | 635 | 19 | 7.6 (3.98) | $C_2^{WW}$ | *Or83c* | *Or83c* |
| 3 | 64,144,317-64,145,939 | 1622 | 10 | 8.07 (6.9) | $C_2^{WW}$ | *LOC118878028* | *CG46026* |
| 3 | 65,102,132-65,103,932 | 1800 | 15 | 11.64 (5.82) | $C_2^{WW}$ | *dj* | *dj* |
| 3 | 65,561,438-65,563,715 | 2277 | 24 | 9.85 (4.63) | $C_2^{EU}$ | - | - |
| 3 | 65,874,336-65,875,145 | 809 | 12 | 7.8 (5.87) | $C_2^{AM}$ | *LOC108019334* | *CG2616* |
| 3 | 65,982,513-65,983,089 | 576 | 13 | 10.55 (6.78) | $C_2^{WW}$ | *LOC136117009* | - |
| 3 | 65,990,559-65,991,700 | 1141 | 12 | 7.49 (6.86) | $C_2^{WW}$ | *LOC108007940* | - |
| 3 | 66,146,262-66,153,885 | 7623 | 107 | 26.75 (5.09) | $C_2^{WW}$ | *LOC108011780* | - |
| 3 | 66,618,563-66,622,771 | 4208 | 51 | 27.41 (5.19) | $C_2^{WW}$ | *Fancm* | *Fancm* |
| 3 | 66,618,563-66,620,103 | 1540 | 35 | 16.48 (4.2) | $C_2^{EU}$ | *Fancm* | *Fancm* |
| 3 | 66,988,086-66,988,323 | 237 | 11 | 9.61 (6.12) | $C_2^{WW}$ | *SNF4Agamma* | *SNF4Agamma* |
| 3 | 66,988,086-66,988,313 | 227 | 9 | 7.24 (5.45) | $C_2^{EU}$ | *SNF4Agamma* | *SNF4Agamma* |
| 3 | 67,003,379-67,004,586 | 1207 | 16 | 15.01 (6.95) | $C_2^{WW}$ | *SNF4Agamma* | *SNF4Agamma* |
| 3 | 67,003,379-67,004,559 | 1180 | 15 | 12.87 (6.85) | $C_2^{EU}$ | *SNF4Agamma* | *SNF4Agamma* |
| 3 | 67,027,806-67,030,204 | 2398 | 28 | 20.79 (5.76) | $C_2^{WW}$ | *meigo* | *meigo* |
| 3 | 67,209,525-67,210,814 | 1289 | 24 | 13.13 (5.52) | $C_2^{EU}$ | *Calx* | *Calx* |
| 3 | 67,270,458-67,271,482 | 1024 | 13 | 8.77 (5.4) | $C_2^{WW}$ | *LOC108018355* | - |
| 3 | 67,376,351-67,378,540 | 2189 | 34 | 17.17 (5.45) | $C_2^{WW}$ | *Prosalpha2* | *Prosalpha2* |
| 3 | 67,451,854-67,453,404 | 1550 | 18 | 8.28 (5.03) | $C_2^{WW}$ | *Octbeta3R* | *Octbeta3R* |
| 3 | 67,493,763-67,495,087 | 1324 | 26 | 7.44 (5.17) | $C_2^{WW}$ | *Octbeta2R* | *Octbeta2R* |
| 3 | 67,648,297-67,650,377 | 2080 | 68 | 17.51 (7.1) | $C_2^{WW}$ | *PolQ* | *PolQ* |
| 3 | 67,648,297-67,648,572 | 275 | 13 | 7.54 (4.48) | $C_2^{AM}$ | *PolQ* | *PolQ* |
| 3 | 67,671,030-67,671,192 | 162 | 12 | 10.06 (6.36) | $C_2^{WW}$ | *Muc91C* | *Muc91C* |
| 3 | 67,717,124-67,718,063 | 939 | 14 | 9.24 (5.11) | $C_2^{WW}$ | *Mekk1* | *Mekk1* |
| 3 | 68,120,579-68,122,202 | 1623 | 14 | 9.57 (4.43) | $C_2^{WW}$ | *LOC108014141* | *CG6040* |
| 3 | 68,157,835-68,158,816 | 981 | 12 | 8.67 (6.13) | $C_2^{WW}$ | *LOC108014141* | *CG6040* |
| 3 | 68,159,410-68,159,762 | 352 | 10 | 8.38 (5.75) | $C_2^{EU}$ | *LOC108014141* | *CG6040* |
| 3 | 68,219,493-68,219,895 | 402 | 18 | 6.94 (4.69) | $C_2^{WW}$ | *LOC108013822* | *CG5555* |
| 3 | 68,326,018-68,326,660 | 642 | 15 | 7.06 (4.34) | $C_2^{AM}$ | - | - |
| 3 | 68,838,360-68,844,193 | 5833 | 18 | 8.28 (5) | $C_2^{WW}$ | *GluClalpha* | *GluClalpha* |

| | | | | | | | |
|---|---|---|---|---|---|---|---|
| 3 | 68,941,829-68,942,505 | 676 | 21 | 7.67 (6.25) | $C_2^{WW}$ | *LOC108015572* | *CG31459* |
| 3 | 68,959,740-68,960,368 | 628 | 18 | 9.53 (4.52) | $C_2^{WW}$ | *Ire1* | *Ire1* |
| 3 | 68,982,662-68,983,991 | 1329 | 15 | 8.22 (3.92) | $C_2^{WW}$ | *Arc42* | *Arc42* |
| 3 | 69,005,900-69,007,886 | 1986 | 33 | 15.47 (7.76) | $C_2^{WW}$ | *ClpX* | *ClpX* |
| 3 | 69,005,900-69,006,486 | 586 | 11 | 7.46 (4.93) | $C_2^{AM}$ | *ClpX* | *ClpX* |
| 3 | 69,125,617-69,127,275 | 1658 | 17 | 11.15 (5.64) | $C_2^{WW}$ | *Hs6st* | *Hs6st* |
| 3 | 69,181,252-69,181,837 | 585 | 13 | 7.13 (6.73) | $C_2^{WW}$ | *LOC108005061* | *CG17193* |
| 3 | 69,186,108-69,189,219 | 3111 | 34 | 12.03 (4.31) | $C_2^{WW}$ | *LOC108005030* | *CG4390* |
| 3 | 69,311,747-69,312,585 | 838 | 22 | 12.78 (6.2) | $C_2^{WW}$ | - | - |
| 3 | 69,311,747-69,312,527 | 780 | 19 | 9.46 (4.96) | $C_2^{EU}$ | - | - |
| 3 | 69,405,460-69,407,339 | 1879 | 28 | 10.24 (5.54) | $C_2^{WW}$ | *Nlg4* | *Nlg4* |
| 3 | 69,527,454-69,528,261 | 807 | 14 | 7.35 (4.6) | $C_2^{WW}$ | *yrt* | *yrt* |
| 3 | 69,593,440-69,596,212 | 2772 | 28 | 10.17 (5.61) | $C_2^{WW}$ | *LOC108013900* | *CG12538* |
| 3 | 69,849,527-69,850,634 | 1107 | 13 | 6.91 (4.18) | $C_2^{EU}$ | *kmr* | *kmr* |
| 3 | 70,288,421-70,290,432 | 2011 | 39 | 10.39 (4.6) | $C_2^{WW}$ | *Npc2b* | *Npc2b* |
| 3 | 70,288,421-70,290,463 | 2042 | 40 | 11.99 (4.65) | $C_2^{AM}$ | *Npc2b* | *Npc2b* |
| 3 | 70,298,211-70,299,321 | 1110 | 28 | 8.72 (4.2) | $C_2^{WW}$ | *CCHa1* | *CCHa1* |
| 3 | 70,445,425-70,446,878 | 1453 | 16 | 9.86 (5.61) | $C_2^{WW}$ | - | - |
| 3 | 70,460,631-70,461,930 | 1299 | 19 | 8.35 (4.7) | $C_2^{WW}$ | *MetRS-m* | *MetRS-m* |
| 3 | 70,585,934-70,586,298 | 364 | 17 | 10.77 (5.38) | $C_2^{WW}$ | *LOC108017200* | *CG9444* |
| 3 | 70,684,544-70,685,541 | 997 | 25 | 12.51 (6.28) | $C_2^{WW}$ | *MtnA* | *MtnA* |
| 3 | 70,700,870-70,703,685 | 2815 | 26 | 10.72 (4.75) | $C_2^{EU}$ | *LOC108019431* | *CG17271* |
| 3 | 70,836,557-70,839,384 | 2827 | 32 | 17.81 (4.72) | $C_2^{WW}$ | *Oamb* | *Oamb* |
| 3 | 70,836,557-70,839,294 | 2737 | 29 | 16.27 (4.37) | $C_2^{AM}$ | *Oamb* | *Oamb* |
| 3 | 70,859,288-70,860,185 | 897 | 9 | 8.48 (6.53) | $C_2^{EU}$ | *LOC108010544* | *CG4000* |
| 3 | 70,928,020-70,929,468 | 1448 | 22 | 8.96 (3.58) | $C_2^{WW}$ | *H* | *H* |
| 3 | 70,976,520-70,979,227 | 2707 | 32 | 8.37 (3.94) | $C_2^{WW}$ | $TFC_2^{AM}$ | $TFC_2^{AM}$ |
| 3 | 71,076,152-71,077,697 | 1545 | 30 | 13.33 (5.25) | $C_2^{WW}$ | *Gfrl* | *Gfrl* |
| 3 | 71,076,419-71,077,117 | 698 | 26 | 10.95 (5.02) | $C_2^{EU}$ | *Gfrl* | *Gfrl* |
| 3 | 71,357,978-71,358,786 | 808 | 14 | 7.27 (5.3) | $C_2^{WW}$ | *PK2-R2* | *PK2-R2* |
| 3 | 71,366,848-71,367,505 | 657 | 20 | 8.06 (3.79) | $C_2^{WW}$ | *LOC108007725* | *CG4009* |
| 3 | 71,409,454-71,410,248 | 794 | 19 | 7.34 (4.06) | $C_2^{WW}$ | *Ns1* | *Ns1* |
| 3 | 71,448,238-71,448,648 | 410 | 15 | 9.58 (4.77) | $C_2^{WW}$ | *Cad86C* | *Cad86C* |
| 3 | 71,559,480-71,560,746 | 1266 | 32 | 11.59 (4.09) | $C_2^{WW}$ | - | - |
| 3 | 71,559,480-71,560,624 | 1144 | 24 | 9.24 (3.9) | $C_2^{EU}$ | - | - |
| 3 | 71,579,643-71,580,518 | 875 | 24 | 6.92 (3.14) | $C_2^{AM}$ | *LOC118877970* | - |
| 3 | 72,051,051-72,052,410 | 1359 | 20 | 7.51 (4.72) | $C_2^{EU}$ | *Rfx* | *Rfx* |
| 3 | 72,385,787-72,386,908 | 1121 | 17 | 10.62 (4.44) | $C_2^{EU}$ | *FER* | *FER* |
| 3 | 73,871,866-73,873,257 | 1391 | 41 | 8.47 (4.83) | $C_2^{WW}$ | *Gr93c* | *Gr93c* |
| 3 | 74,162,047-74,163,088 | 1041 | 13 | 7.2 (4.15) | $C_2^{AM}$ | *BomT3* | *BomT3* |
| 3 | 75,584,393-75,584,991 | 598 | 11 | 7.18 (5.01) | $C_2^{EU}$ | *LOC108011504* | *CG13640* |
| 3 | 75,806,245-75,809,289 | 3044 | 33 | 7.32 (4.02) | $C_2^{WW}$ | *LOC108011536* | *CG31125* |
| 3 | 75,806,263-75,807,974 | 1711 | 29 | 8.73 (4.52) | $C_2^{WW}$ | *LOC108011524* | *CG6695* |
| 3 | 77,240,194-77,241,076 | 882 | 16 | 9.19 (5.38) | $C_2^{WW}$ | *Trnat-ugu* | - |
| 3 | 77,240,194-77,241,113 | 919 | 17 | 11.2 (4.81) | $C_2^{EU}$ | *Trnat-ugu* | - |
| 3 | 77,337,303-77,338,189 | 886 | 56 | 15.19 (4.48) | $C_2^{WW}$ | *LOC108017397* | - |
| 3 | 77,337,499-77,338,008 | 509 | 32 | 13.92 (4.68) | $C_2^{EU}$ | *LOC108017397* | - |
| 3 | 77,462,962-77,463,477 | 515 | 18 | 7.01 (4.21) | $C_2^{WW}$ | *LOC108019257* | *CG4702* |
| 3 | 77,724,188-77,724,766 | 578 | 13 | 8.93 (5.17) | $C_2^{WW}$ | *Sulf1* | *Sulf1* |
| 3 | 77,894,990-77,895,640 | 650 | 12 | 9.52 (5.22) | $C_2^{WW}$ | *Cad89D* | *Cad89D* |
| 3 | 78,499,920-78,500,536 | 616 | 17 | 11.78 (4.11) | $C_2^{EU}$ | *Osi22* | *Osi22* |
| 3 | 79,047,294-79,047,924 | 630 | 15 | 8.93 (4.62) | $C_2^{WW}$ | *LOC108011585* | *CG10909* |
| 3 | 79,148,789-79,150,224 | 1435 | 12 | 7.91 (6.17) | $C_2^{WW}$ | *LOC108005702* | *CG31345* |
| 3 | 79,525,967-79,526,692 | 725 | 16 | 11.38 (5.48) | $C_2^{WW}$ | *fru* | *fru* |
| 3 | 79,525,967-79,526,606 | 639 | 15 | 9.49 (5.22) | $C_2^{EU}$ | *fru* | *fru* |
| 3 | 79,917,736-79,918,318 | 582 | 16 | 7.16 (3.62) | $C_2^{WW}$ | *sr* | *sr* |
| 3 | 80,166,803-80,167,826 | 1023 | 12 | 8.62 (5.56) | $C_2^{WW}$ | *sra* | *sra* |
| 3 | 80,353,395-80,354,569 | 1174 | 15 | 10.64 (6.91) | $C_2^{WW}$ | - | - |
| 3 | 81,321,704-81,324,105 | 2401 | 32 | 13.45 (4.26) | $C_2^{WW}$ | *LOC108007632* | *CG33203* |
| 3 | 81,357,569-81,358,490 | 921 | 22 | 7.15 (4.3) | $C_2^{WW}$ | *Doa* | *Doa* |

| | | | | | | | |
|---|---|---|---|---|---|---|---|
| 3 | 81,930,891-81,931,887 | 996 | 19 | 12.81 (4.65) | $C_2^{WW}$ | *pins* | *pins* |
| 3 | 82,710,180-82,710,501 | 321 | 10 | 7.07 (4.3) | $C_2^{WW}$ | *BRWD3* | *BRWD3* |
| 3 | 84,494,796-84,495,243 | 447 | 15 | 6.86 (3.95) | $C_2^{WW}$ | *ymp* | *ymp* |
| 3 | 85,617,115-85,618,318 | 1203 | 29 | 9.46 (3.84) | $C_2^{WW}$ | *rdog* | *rdog* |
| 3 | 85,617,115-85,618,129 | 1014 | 23 | 7.64 (4.17) | $C_2^{AM}$ | *rdog* | *rdog* |
| 3 | 86,125,148-86,126,005 | 857 | 18 | 9.89 (5.54) | $C_2^{WW}$ | *LOC108010729* | *CG7567* |
| 3 | 86,162,308-86,164,606 | 2298 | 23 | 6.92 (4.46) | $C_2^{AM}$ | *LOC108019458* | *CG1907* |
| 3 | 86,313,451-86,314,841 | 1390 | 15 | 10.46 (5.39) | $C_2^{WW}$ | *Acam* | *Acam* |
| 3 | 86,839,458-86,840,165 | 707 | 26 | 8.38 (4.46) | $C_2^{WW}$ | *RpS3* | *RpS3* |
| 3 | 87,453,952-87,454,469 | 517 | 17 | 8.94 (3.22) | $C_2^{AM}$ | *LOC108007848* | - |
| 3 | 87,506,586-87,509,876 | 3290 | 51 | 17.93 (3.88) | $C_2^{EU}$ | *Hmgcr* | *Hmgcr* |
| 3 | 88,572,321-88,573,780 | 1459 | 22 | 10.29 (4.28) | $C_2^{WW}$ | *LOC108007385* | - |
| 3 | 89,139,363-89,140,135 | 772 | 20 | 7.03 (3.32) | $C_2^{AM}$ | *LOC108016129* | *CG5880* |
| 3 | 89,791,461-89,792,811 | 1350 | 24 | 7.1 (4.01) | $C_2^{WW}$ | *scrib* | *scrib* |
| 3 | 89,799,315-89,799,897 | 582 | 8 | 6.94 (6.9) | $C_2^{EU}$ | *LOC118878096* | - |
| 3 | 90,542,111-90,542,926 | 815 | 13 | 9.02 (6.33) | $C_2^{WW}$ | *SppL* | *SppL* |
| 3 | 91,603,781-91,604,031 | 250 | 23 | 8.8 (4) | $C_2^{WW}$ | *LOC108008038* | - |

Table S6: Description of all the significant windows for the chromosome 3 (n=195), with annotated positional candidate genes. The table gives for each significant window (i) its position in the new *dsu_isojap1.0* assembly of the *D. suzukii* genome; (ii) its size in bp; (iii) its number of SNPs; (iv) the window highest maximum Lindley score (Lscore) and its most significant $C_2$ P-values (in $-\log_{10}$ scale); (v) the $C_2$ statistic corresponding to this highest Lscore; (vi) the positional candidate gene associated and (vii) its corresponding *D. melanogaster* ortholog.

| Chr. | Position in bp | Size in bp | nsnps | Lscore ($-\log_{10}$ P$_{C2}$) | $C_2$ | Gene | Ortholog |
|---|---|---|---|---|---|---|---|
| 4 | 19,963-21,589 | 1626 | 16 | 10.93 (4.78) | $C_2^{EU}$ | *LOC118878296* | - |
| 4 | 28,989-37,445 | 8456 | 23 | 5.91 (3.84) | $C_2^{EU}$ | - | - |
| 4 | 590,198-591,974 | 1776 | 18 | 9.52 (4.53) | $C_2^{WW}$ | *fuss* | *fuss* |
| 4 | 872,114-872,717 | 603 | 10 | 6.01 (4.04) | $C_2^{WW}$ | *Sox102F* | *Sox102F* |
| 4 | 876,191-876,980 | 789 | 8 | 5.49 (4.74) | $C_2^{WW}$ | *Sox102F* | *Sox102F* |
| 4 | 1,304,042-1,309,833 | 5791 | 11 | 6.89 (4.39) | $C_2^{EU}$ | *Tdg* | *Tdg* |
| 4 | 1,320,563-1,321,698 | 1135 | 13 | 7 (4.08) | $C_2^{EU}$ | *LOC139353324* | - |
| 4 | 1,385,246-1,386,591 | 1345 | 23 | 19.58 (5.92) | $C_2^{EU}$ | *zfh2* | *zfh2* |
| 4 | 1,482,729-1,486,912 | 4183 | 14 | 5.84 (3.32) | $C_2^{EU}$ | *lgs* | *lgs* |
| 4 | 1,521,119-1,525,419 | 4300 | 14 | 6.77 (6.29) | $C_2^{WW}$ | *LOC139353319* | - |
| 4 | 1,998,487-1,999,638 | 1151 | 8 | 5.72 (5.88) | $C_2^{EU}$ | *Crk* | *Crk* |

Table S7: Description of all the significant windows for the chromosome 4 (n=11), with annotated positional candidate genes. The table gives for each significant window (i) its position in the new *dsu_isojap1.0* assembly of the *D. suzukii* genome; (ii) its size in bp; (iii) its number of SNPs; (iv) the window highest maximum Lindley score (Lscore) and its most significant $C_2$ P-values (in $-\log_{10}$ scale); (v) the $C_2$ statistic corresponding to this highest Lscore; (vi) the positional candidate gene associated and (vii) its corresponding *D. melanogaster* ortholog.

| Chr. | Position in bp | Size in bp | nsnps | Lscore ($-\log_{10}$ P$_{C2}$) | $C_2$ | Gene | Ortholog |
|---|---|---|---|---|---|---|---|
| X | 3,406,420-3,413,303 | 6883 | 26 | 17.39 (5.81) | $C_2^{WW}$ | - | - |
| X | 4,515,264-4,518,631 | 3367 | 9 | 6.23 (3.87) | $C_2^{WW}$ | *Ogg1* | *Ogg1* |
| X | 8,840,140-8,843,873 | 3733 | 8 | 7.29 (5.73) | $C_2^{WW}$ | *LOC108020178* | *CG5254* |
| X | 10,238,523-10,239,089 | 566 | 14 | 7.53 (5.63) | $C_2^{EU}$ | *LOC108016570* | - |
| X | 11,220,921-11,222,068 | 1147 | 9 | 7.38 (6.09) | $C_2^{WW}$ | *mamo* | *mamo* |
| X | 11,323,503-11,325,439 | 1936 | 17 | 12.66 (5.96) | $C_2^{WW}$ | *Trnas-aga* | - |
| X | 12,412,613-12,412,846 | 233 | 13 | 7.06 (5.1) | $C_2^{EU}$ | *LOC108006288* | *CG9981* |
| X | 12,417,807-12,418,092 | 285 | 12 | 6.93 (4.71) | $C_2^{EU}$ | *mei-41* | *mei-41* |
| X | 12,417,885-12,418,097 | 212 | 10 | 6.07 (5.09) | $C_2^{WW}$ | *mei-41* | *mei-41* |
| X | 12,431,127-12,431,344 | 217 | 9 | 6.39 (5.22) | $C_2^{WW}$ | *LOC108006273* | *CG9992* |
| X | 12,821,875-12,831,789 | 9914 | 13 | 9.17 (5.9) | $C_2^{WW}$ | *LOC139353334* | - |
| X | 13,072,527-13,073,689 | 1162 | 12 | 6.34 (4.43) | $C_2^{WW}$ | *shi* | *shi* |
| X | 13,696,115-13,697,427 | 1312 | 10 | 9.1 (7.15) | $C_2^{EU}$ | *sdt* | *sdt* |
| X | 13,718,304-13,718,721 | 417 | 8 | 7.05 (6.66) | $C_2^{EU}$ | *sdt* | *sdt* |
| X | 14,219,391-14,220,467 | 1076 | 11 | 8.84 (5.58) | $C_2^{WW}$ | *LOC108018858* | *CG1628* |
| X | 14,583,384-14,584,611 | 1227 | 10 | 7.02 (4.8) | $C_2^{WW}$ | *Obp19a* | *Obp19a* |
| X | 15,093,018-15,093,439 | 421 | 11 | 7.35 (5.57) | $C_2^{EU}$ | - | - |
| X | 15,503,913-15,505,271 | 1358 | 10 | 8.29 (6.92) | $C_2^{EU}$ | - | - |
| X | 15,527,257-15,529,110 | 1853 | 8 | 6.73 (8.59) | $C_2^{EU}$ | - | - |
| X | 15,589,507-15,591,066 | 1559 | 20 | 8.25 (4.29) | $C_2^{EU}$ | *LOC118878157* | *CG14265* |
| X | 15,878,099-15,880,687 | 2588 | 23 | 10.53 (4.92) | $C_2^{WW}$ | *LOC118878171* | - |
| X | 16,415,048-16,415,610 | 562 | 11 | 7.94 (6.25) | $C_2^{EU}$ | *ec* | *ec* |
| X | 16,608,496-16,609,985 | 1489 | 11 | 6.58 (4.69) | $C_2^{WW}$ | - | - |
| X | 16,723,698-16,725,318 | 1620 | 23 | 6.8 (4.82) | $C_2^{EU}$ | *Cmtr1* | *Cmtr1* |
| X | 16,786,001-16,787,289 | 1288 | 12 | 6.54 (7.65) | $C_2^{EU}$ | *Hers* | *Hers* |
| X | 18,316,644-18,317,737 | 1093 | 9 | 7.5 (6.89) | $C_2^{EU}$ | *LOC108011160* | *CG15765* |
| X | 19,809,392-19,810,859 | 1467 | 16 | 11 (7.07) | $C_2^{WW}$ | *Flo2* | *Flo2* |
| X | 19,809,392-19,810,859 | 1467 | 16 | 7.04 (4.31) | $C_2^{EU}$ | *Flo2* | *Flo2* |
| X | 19,913,834-19,914,358 | 524 | 19 | 9.07 (4.73) | $C_2^{EU}$ | *Nrd1* | *Nrd1* |
| X | 19,937,004-19,938,447 | 1443 | 8 | 7.72 (8.59) | $C_2^{WW}$ | *LOC139353364* | - |
| X | 19,998,780-20,000,289 | 1509 | 11 | 6.55 (5.37) | $C_2^{WW}$ | *LOC108005118* | *CG1578* |
| X | 20,080,285-20,080,866 | 581 | 18 | 7.9 (4.36) | $C_2^{WW}$ | *inaF-A* | *inaF-A* |
| X | 20,382,372-20,388,371 | 5999 | 15 | 13.38 (6.07) | $C_2^{EU}$ | *Prx2* | *Prx2* |
| X | 20,479,390-20,480,318 | 928 | 14 | 9.17 (4.94) | $C_2^{EU}$ | *mew* | *mew* |
| X | 20,683,444-20,683,816 | 372 | 26 | 15.34 (4.78) | $C_2^{WW}$ | *LOC108005385* | *CG13003* |
| X | 20,782,296-20,783,873 | 1577 | 12 | 8.13 (6.13) | $C_2^{EU}$ | *LOC108005458* | *CG4928* |
| X | 21,067,868-21,068,116 | 248 | 12 | 6.94 (5.84) | $C_2^{EU}$ | *RpS5a* | *RpS5a* |
| X | 21,070,167-21,070,999 | 832 | 27 | 10.05 (3.77) | $C_2^{EU}$ | *RpS5a* | *RpS5a* |
| X | 21,074,130-21,075,885 | 1755 | 39 | 19.31 (6.41) | $C_2^{EU}$ | *Chchd2* | *Chchd2* |
| X | 21,127,624-21,128,242 | 618 | 11 | 6.8 (4.85) | $C_2^{EU}$ | *Chpf* | *Chpf* |
| X | 21,195,982-21,196,272 | 290 | 14 | 7.38 (7.14) | $C_2^{EU}$ | *rsh* | *rsh* |
| X | 21,252,695-21,253,654 | 959 | 15 | 10.51 (4.95) | $C_2^{WW}$ | *rsh* | *rsh* |
| X | 21,252,695-21,253,598 | 903 | 13 | 7.51 (4.05) | $C_2^{EU}$ | *rsh* | *rsh* |
| X | 21,591,603-21,591,883 | 280 | 12 | 6.41 (5.82) | $C_2^{EU}$ | *LOC108005411* | *CG32647* |
| X | 21,915,844-21,916,289 | 445 | 14 | 9.84 (5.05) | $C_2^{WW}$ | *LOC108004576* | *CG32751* |
| X | 22,063,281-22,064,078 | 797 | 12 | 9.58 (6.4) | $C_2^{EU}$ | *Tsp5D* | *Tsp5D* |
| X | 22,139,022-22,139,368 | 346 | 14 | 8.89 (5.79) | $C_2^{EU}$ | *Es2* | *Es2* |
| X | 22,193,820-22,195,746 | 1926 | 9 | 6.32 (4.81) | $C_2^{WW}$ | *LOC136117265* | - |
| X | 22,193,820-22,195,746 | 1926 | 9 | 6.24 (5) | $C_2^{EU}$ | *LOC136117265* | - |
| X | 22,280,846-22,281,809 | 963 | 19 | 6.72 (3.44) | $C_2^{WW}$ | *LOC108004780* | - |
| X | 23,913,583-23,915,081 | 1498 | 11 | 8.26 (6.18) | $C_2^{WW}$ | - | - |
| X | 24,109,619-24,110,178 | 559 | 14 | 6.56 (3.74) | $C_2^{WW}$ | *LOC108004773* | *CG32719* |
| X | 25,060,448-25,060,704 | 256 | 16 | 6.79 (4.07) | $C_2^{WW}$ | *CrebB* | *CrebB* |
| X | 25,178,700-25,179,490 | 790 | 20 | 6.26 (5.14) | $C_2^{WW}$ | *LOC108010457* | *CG32547* |
| X | 26,156,841-26,161,080 | 4239 | 9 | 6.56 (5.75) | $C_2^{WW}$ | - | - |
| X | 26,360,462-26,384,075 | 23613 | 13 | 6.45 (6.09) | $C_2^{EU}$ | - | - |
| X | 26,543,420-26,547,062 | 3642 | 13 | 12.31 (5.24) | $C_2^{WW}$ | - | - |
| X | 26,642,716-26,649,899 | 7183 | 23 | 17.41 (8.48) | $C_2^{WW}$ | - | - |

| | | | | | | | |
|---|---|---|---|---|---|---|---|
| X | 26,646,089-26,649,892 | 3803 | 15 | 13.86 (7.01) | $C_2^{EU}$ | - | - |
| X | 26,869,170-26,870,151 | 981 | 9 | 6.52 (4.33) | $C_2^{WW}$ | - | - |
| X | 27,437,759-27,441,512 | 3753 | 16 | 7.77 (4.43) | $C_2^{EU}$ | - | - |
| X | 27,712,894-27,715,778 | 2884 | 16 | 13.5 (7.68) | $C_2^{EU}$ | - | - |
| X | 27,881,396-27,883,080 | 1684 | 20 | 7.77 (3.45) | $C_2^{WW}$ | - | - |
| X | 28,446,142-28,448,353 | 2211 | 10 | 6.01 (4.9) | $C_2^{EU}$ | *LOC108020464* | *CG12061* |
| X | 28,577,275-28,581,889 | 4614 | 10 | 6.98 (4.98) | $C_2^{WW}$ | *zyd* | *zyd* |
| X | 28,614,879-28,617,976 | 3097 | 33 | 9.19 (3.74) | $C_2^{EU}$ | *FucTC* | *FucTC* |
| X | 28,724,256-28,726,568 | 2312 | 15 | 9.6 (4.16) | $C_2^{EU}$ | - | - |
| X | 28,724,643-28,726,568 | 1925 | 13 | 10.45 (4.72) | $C_2^{WW}$ | - | - |
| X | 28,737,148-28,743,175 | 6027 | 11 | 8.02 (8) | $C_2^{EU}$ | *LOC118878189* | |
| X | 28,828,471-28,830,076 | 1605 | 13 | 8.38 (4.59) | $C_2^{WW}$ | - | - |
| X | 28,930,581-28,943,082 | 12501 | 16 | 8.5 (4.17) | $C_2^{WW}$ | - | - |
| X | 30,128,029-30,139,270 | 11241 | 27 | 17.38 (4.78) | $C_2^{WW}$ | *LOC136117410* | - |
| X | 30,767,370-30,799,878 | 32508 | 10 | 8.65 (6.1) | $C_2^{WW}$ | - | - |
| X | 31,518,750-31,519,341 | 591 | 7 | 6.65 (6.07) | $C_2^{WW}$ | - | - |

Table S8: Description of all the significant windows for the chromosome X (n=74), with annotated positional candidate genes. The table gives for each significant window (i) its position in the new *dsu_isojap1.0* assembly of the *D. suzukii* genome; (ii) its size in bp; (iii) its number of SNPs; (iv) the window highest maximum Lindley score (Lscore) and its most significant $C_2$ P-values (in $-\log_{10}$ scale); (v) the $C_2$ statistic corresponding to this highest Lscore; (vi) the positional candidate gene associated and (vii) its corresponding *D. melanogaster* ortholog.

#### Supplementary Figures

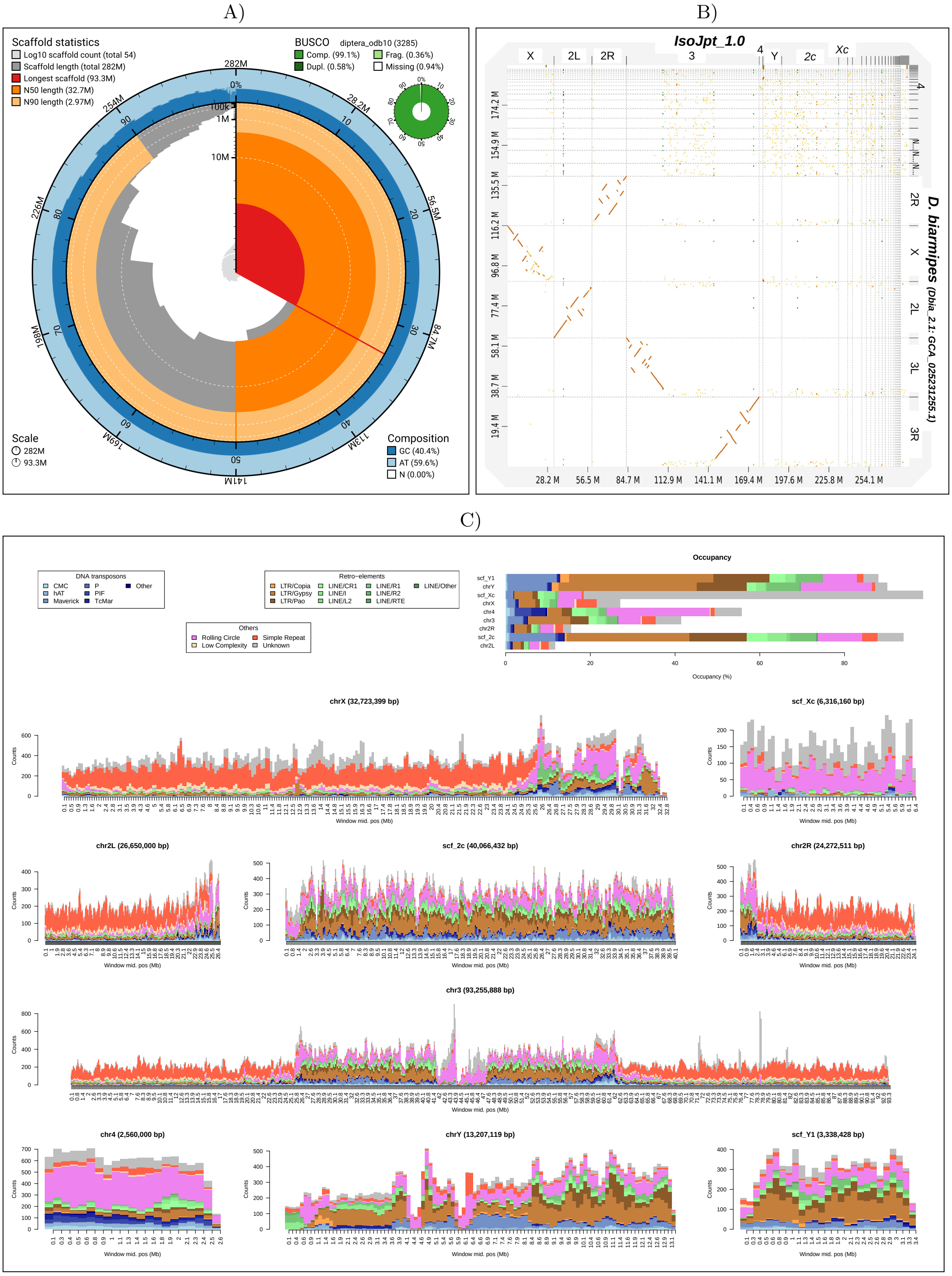


Figure S1: **Statistics of the novel *dsu_isojap1.0* male chromosome-level assembly of the *Drosophila suzukii* Japanese isofemale line.** A) Snailplot built using the `BlobToolKit v4.3.5` viewer (Challis *et al.*, 2020) based on the results from `BUSCO v5.4.7` for the `diptera_odb10` dataset that includes 3,285 genes (Manni *et al.*, 2021). B) Dotplot generated with `dgenies v1.5.0` (Cabanettes and Klopp, 2018) (default options) comparing the *dsu_isojap1.0* with the `Dbia2.1` reference assembly (GenBank ID: GCF_025231255.1) of the closely related species *Drosophila biarmipes*. C) Distribution of the number of copies of different families of repetitive elements over the assembly chromosomes per non overlapping 250 kb windows. The upper-right gives the overall occupancy per chromosome (in % of their length). The LTR order was found predominant (20.9%), followed by LINEs (8.32%), DNA transposons (6.66%) and Rolling Circles (RC, 6.59%). Repetitive sequences occupy almost all chrY (89.8%), scf_Y1 (87.6%), scf_2c (93.7%) and scf_Xc (98.5%) scaffolds and a high proportion of chr4 (53.7%). Conversely, the 2L (11.2%), 2R (14.8%) and chrX (25.7%) displayed the lowest proportion, the full chr3 scaffold an intermediary value of 40.4%.

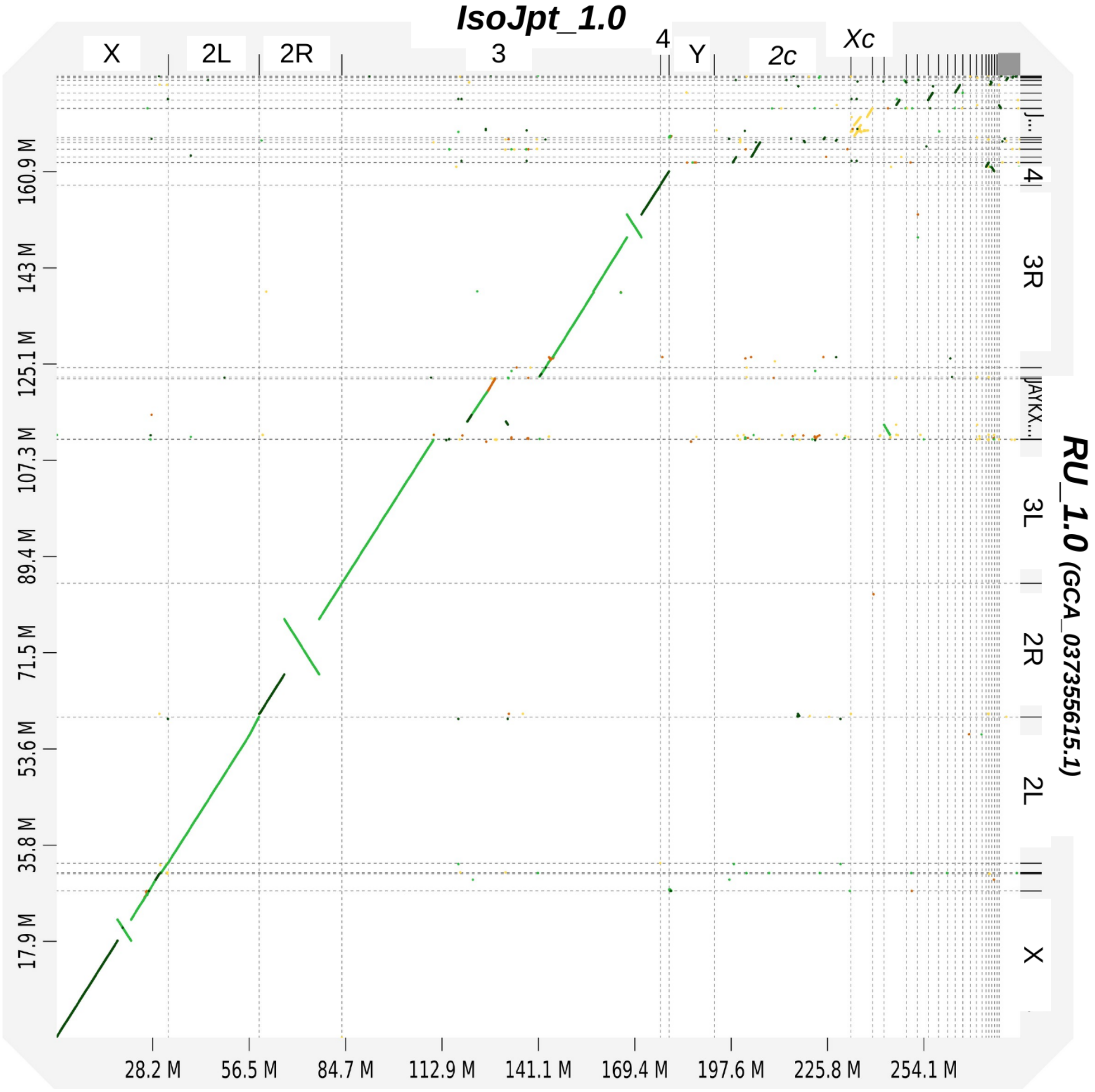


Figure S2: Comparison of the new *dsu_isojap1.0* reference assembly with the `RU_1.0`. The `RU_1.0` chromosome level assembly released in March 2024 on the NCBI repository (see Table 1) remains partial as it lacks ca. 100 Mb of mostly TE-rich and gene poor regions and was obtained from an US isofemale line. The dotplot was generated with `dgenies v1.5.0` (Cabanettes and Klopp, 2018) run with default options. Note that the `RU_1.0` lacks ca. 100 Mb of mostly TE-rich and gene poor regions and was obtained from an US isofemale line. The two assemblies are in very good agreement, although three large-scale inversions could be identified on the X (from position 17.94 to 21.88 Mb), the 2R (from position 7.413 to 17.59 Mb) and 3R (from position 83.51 to 87.75 Mb) chromosomes. More precisely, 40 blocks of sequence similarity $> 1Mb$ totaling 171.5 Mb (i.e. 96.1% of the `RU_1.0` assembly) could be found with only 1.33% of nucleotide divergence.

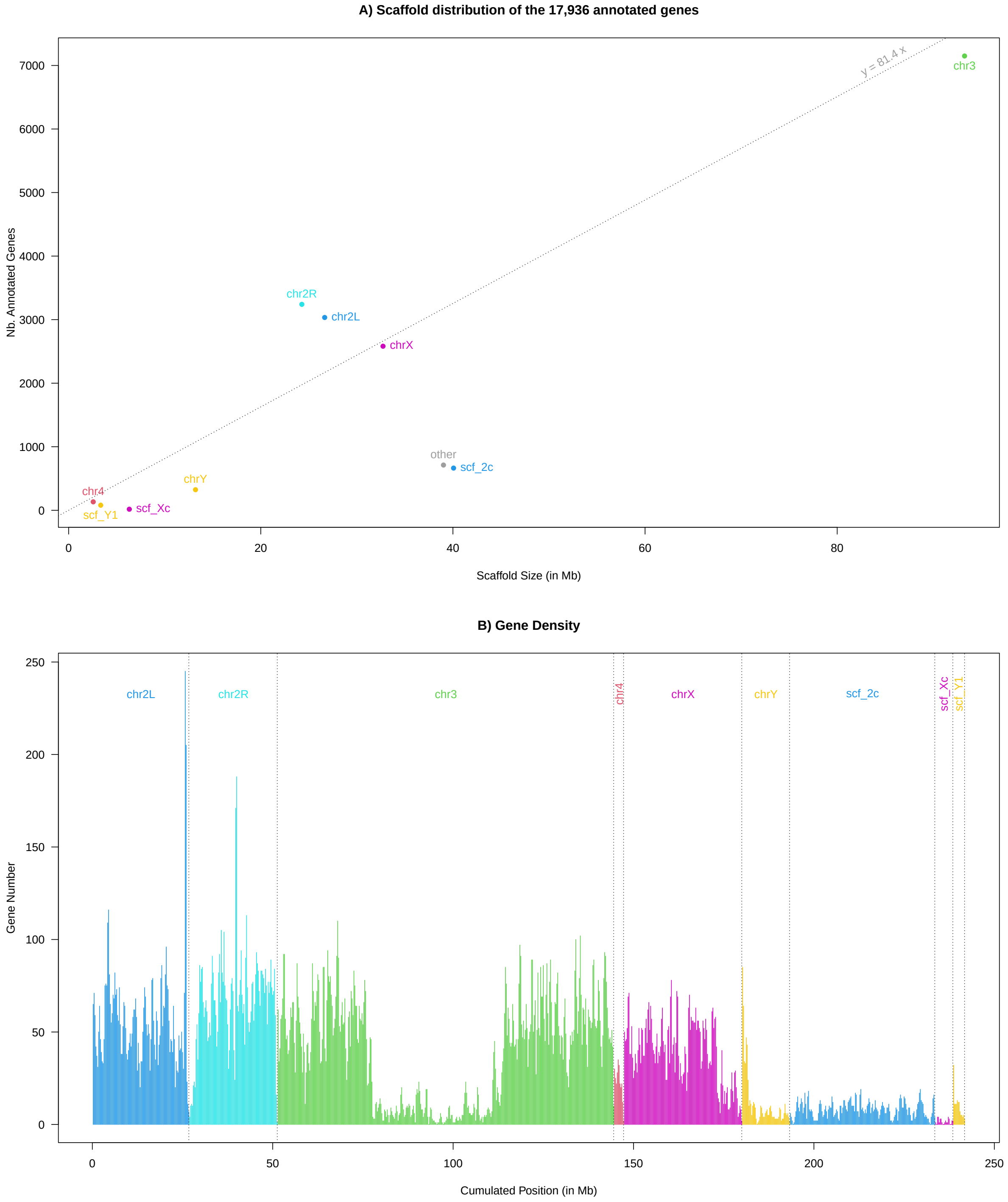


Figure S3: Summary of gene annotation of the new *dsu_isojap1.0* reference assembly obtained with the NCBI Eukaryotic Genome Annotation Pipeline (detailed in `https://www.ncbi.nlm.nih.gov/refseq/annotation_euk/Drosophila_suzukii/GCF_043229965.1-RS_2025_01/`). A) Number of annotated genes per scaffold as a function of their length. The dashed line represents the linear regression model fitted without an intercept on chromosomes 2L, 2R, 3, 4, X and Y, leading to an estimated gene density of 81.4 genes per Mb. B) Number of genes per sliding 500 kb sliding windows over (gene density) over the main scaffolds.

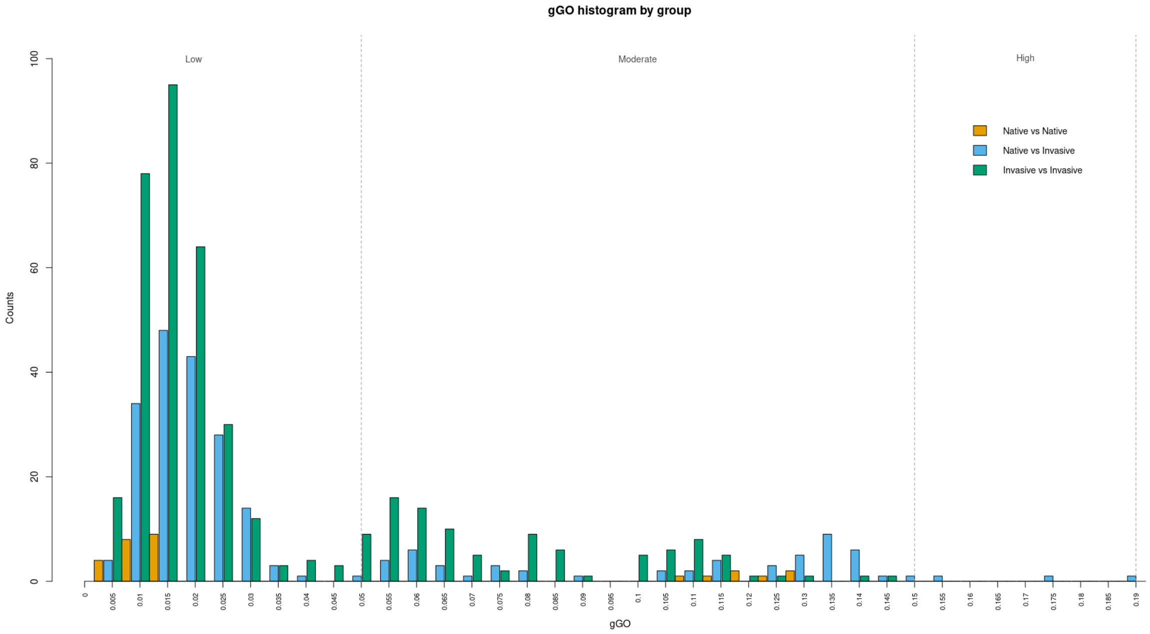


Figure S4: Histogram showing the distribution of gGO values between pairs of the 37 studied populations. Bars represent the number of population pairs falling into each gGO bin, grouped by comparison type: native vs native (orange), native vs invasive (blue), and invasive vs invasive (green). Vertical dashed lines indicate thresholds for interpreting offset magnitudes: Low (gGO $\leq$ 0.05), Moderate ($0.05 <$ gGO $\leq 0.15$), and High (gGO $> 0.15$)

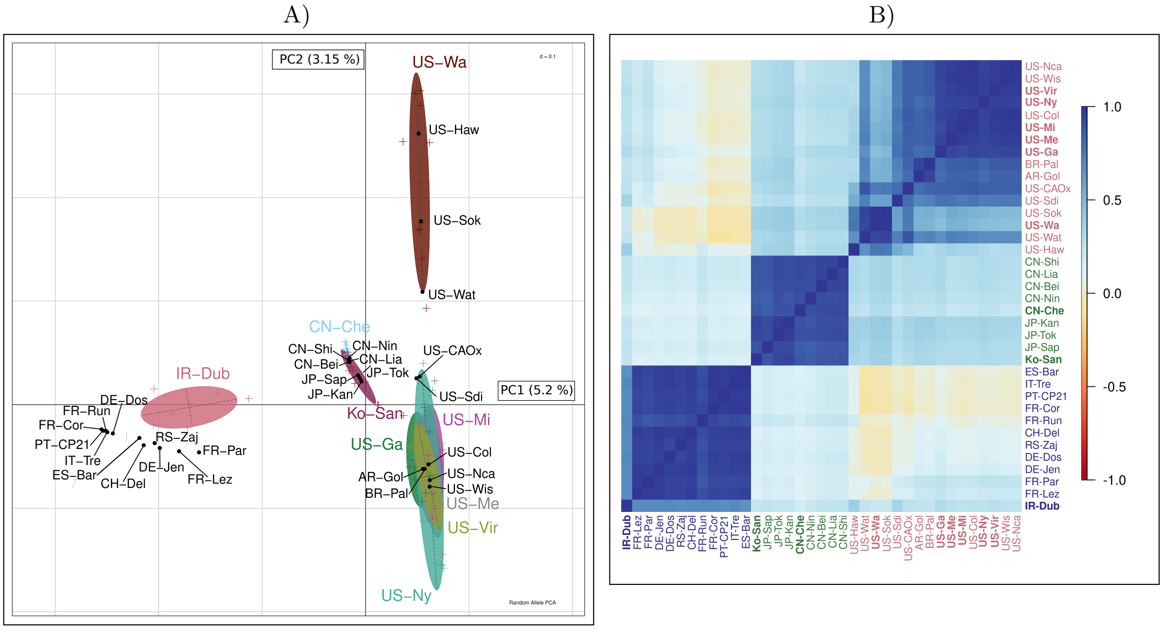


Figure S5: Structuring of genetic diversity across the 37 *D. suzukii* populations studied on X-chromosome variants. A) Random allele PCA. Pool-sequenced populations are represented by black dots, while individual sequencing samples are indicated by colored crosses. Colored ellipses group individual samples from the same population. B) Correlation plot of the scaled covariance matrix $\Omega$ estimated by BayPass for X-chromosome, which incorporates neutral population structure. Individual sequencing sample names are in bold. This figure shows minor differences with the figure performed for autosomal variants (Figure 2B and C). For the random allele PCA, the main notable difference is that, surprisingly, the South American populations (AR-Gol and BR-Pal) no longer clustered most closely with the two Southern Californian populations. Instead, they grouped with other U.S. populations. Also, and as expected from higher overall differentiation on the X chromosome, the percentage of variance explained by the first two PC was higher (Figure S6). Compared to the autosomal results, the estimated $\Omega$ matrix for X-chromosome showed less clear differentiation between Chinese and Japanese population groups. Another surprising result was that US-Wa and US-Sok appeared more closely related to US-CAOx.

A) B)

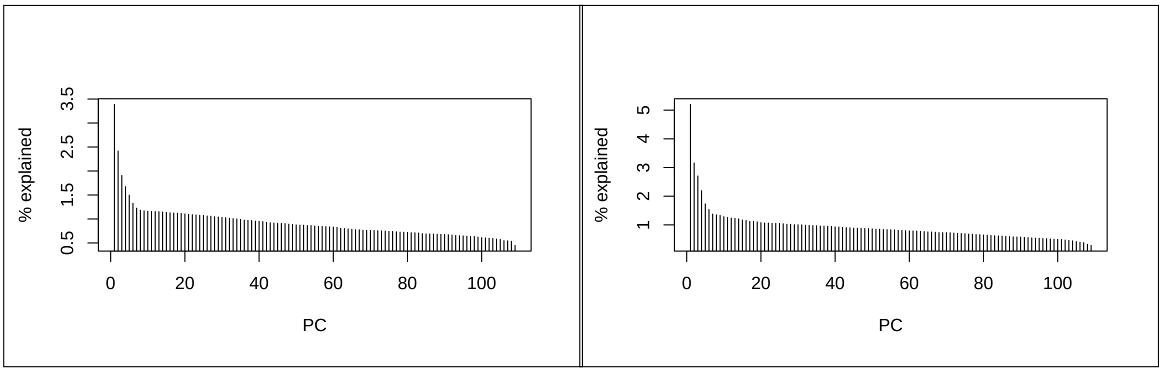


Figure S6: Percentage of variance explained along each Random Allele PCA axis. A) for autosomes and B) for X-chromosome.

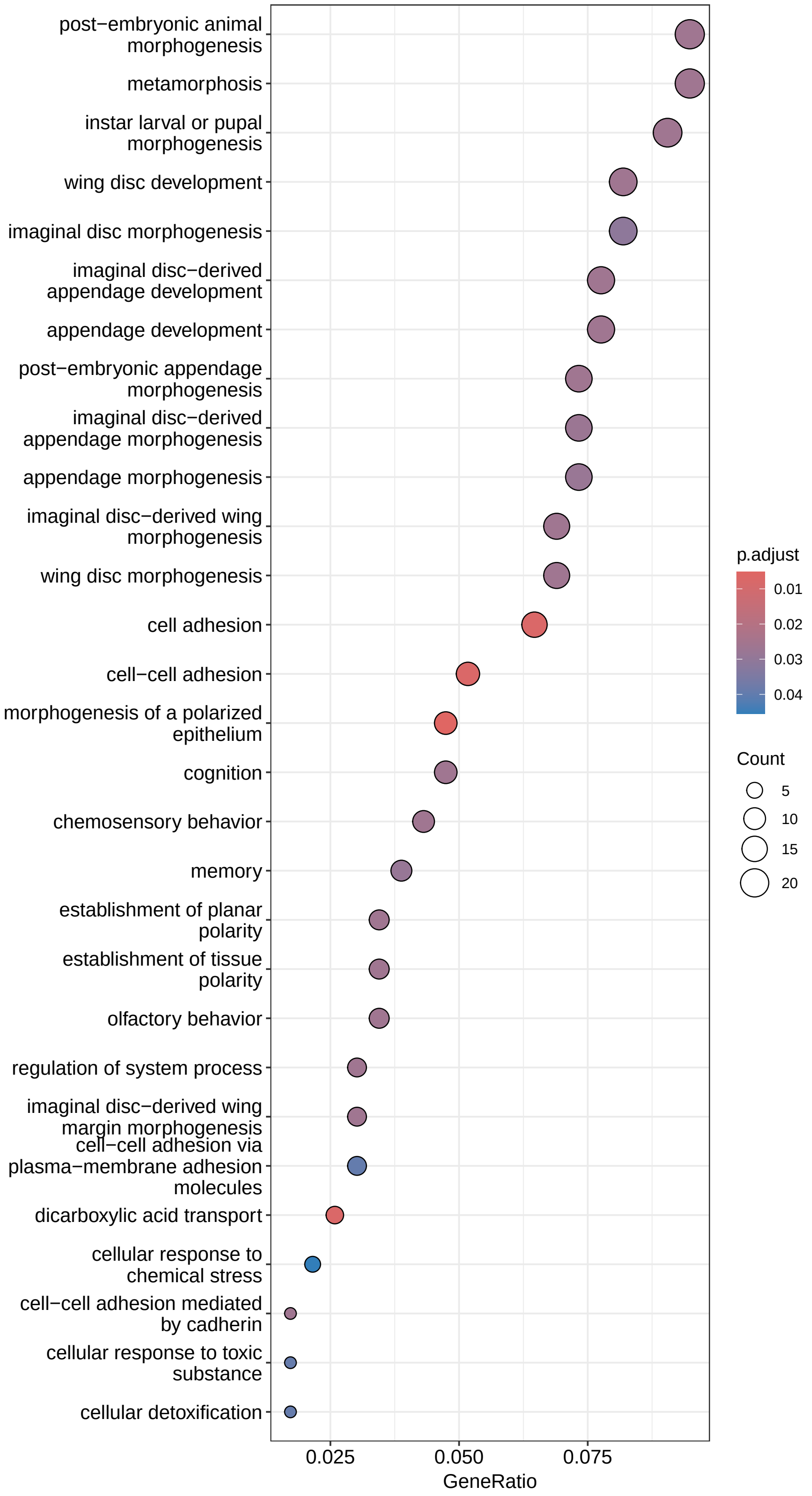


Figure S7: Bubble plot depicting the Gene Ontology (GO) enrichment analysis for Biological Process of the 322 candidate genes. The chart highlights the 29 GO terms with significant enrichment. Circle color represents the adjusted p-value for significant enrichment, while circle size corresponds to the number of genes associated with each GO term. The Gene Ratio reflects the proportion of enriched genes relative to the total number of candidate genes.

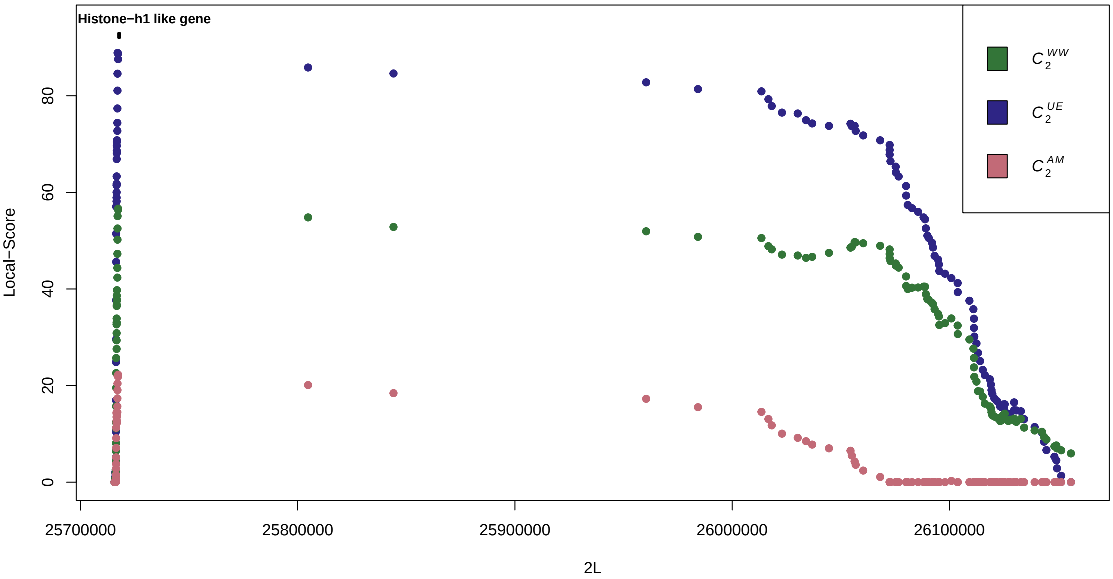


Figure S8: Manhattan plot of the region of *Histone-h1 like* gene (located on chromosome 2L), associated to the most significant peak amongst the three contrast. The position of the gene is indicated in black.

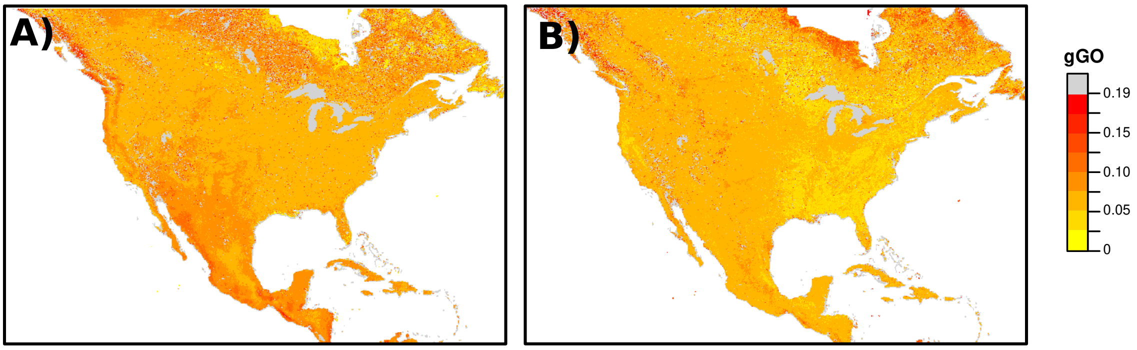


Figure S9: Comparison of the gGO estimates over North American invaded areas when taking A) US-Haw or B) CN-Nin populations as source reference. See Main text for details regarding the choice of source reference when invasion involved several source populations. Gray pixels represent outlying gGO values (as defined in M&M).

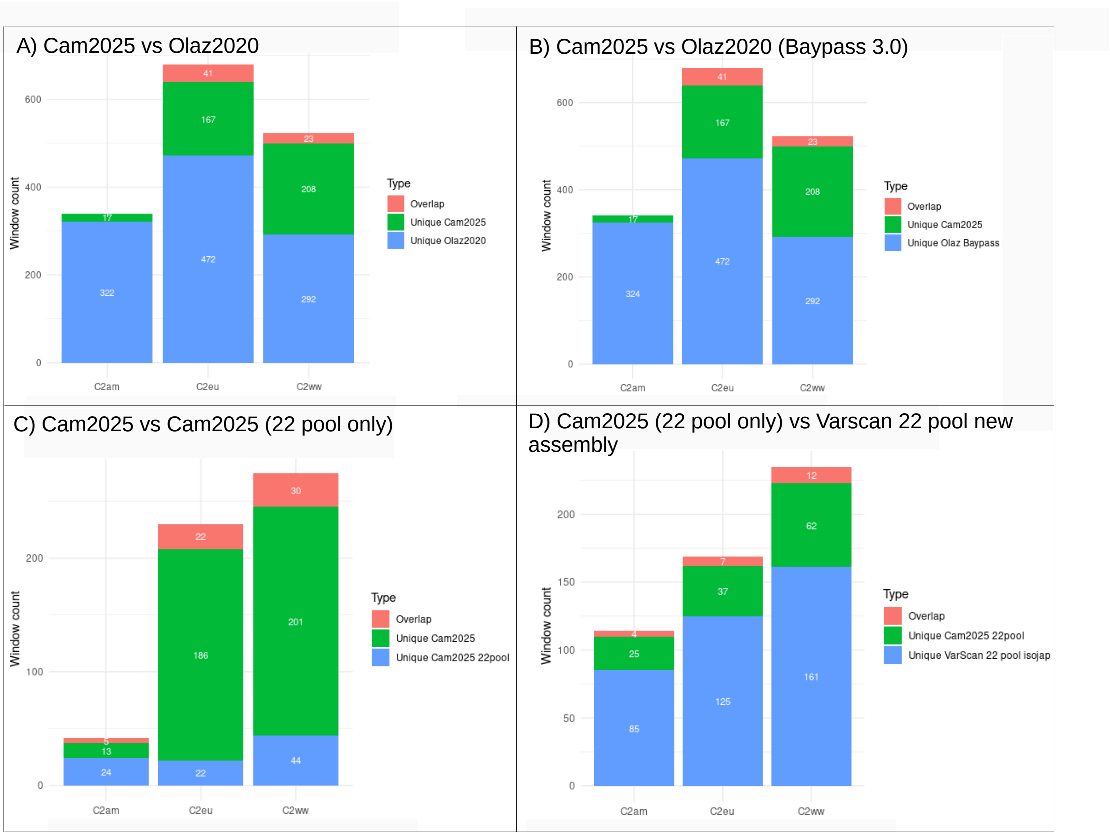


Figure S10: Comparison of significant genomic windows across different datasets and analytical frameworks. Each panel compares the genomic windows identified as significant by the $C_2$ statistics ($C_2^{AM}$, $C_2^{EU}$, $C_2^{WW}$) under varying datasets or analysis parameters. (A) Comparison of the local score windows detected in the present study (Cam2025) and those detected with the original $C_2$ estimates obtained by Olazcuaga *et al.* (2020). The corresponding original dataset (referred to as Olaz20, hereafter) consisting of Pool-Seq data for 22 population samples (all in common with the present study but DE-Jen sample which has been here replaced by a new Pool-Seq sample in Cam2025, see the main text and Table S1) which were mapped on the previous WT3-2.0 assembly (Paris *et al.*, 2020); with variant calling performed using `VarScan` (v2.3.4; Koboldt *et al.*, 2012) and analyzed with BayPass v2.1. Note that variant positions were transferred to the new *dsu_isojap1.0* assembly using LiftOver, available from the UCSC repository: `https://hgdownload.soe.ucsc.edu/downloads.html`. While the windows overlap remains modest, some shared windows are observed — suggesting that despite substantial differences in data structure, SNP calling, and analysis pipeline, the local score method is still able to detect similar $C_2$ signals. (B) Comparison between Cam2025 and Olaz2020 analyzed with BayPass v3.0 instead of v2.1 (with all other parameters unchanged). The overlap remains almost identical to Panel A (as the underlying $C_2$ estimates), indicating that the BayPass version does not explain the differences in detected $C_2$ windows. (C) Comparison between Cam2025 (all 37 populations) and a subset using only the 22 Pool-Seq populations used by Olazcuaga *et al.* (2020). The overlap here is similar to the one in panel A, with 57 overlapping windows against 64 in panel A. This suggests that the reduced number of populations largely explains the limited overlap with the (Olazcuaga *et al.*, 2020) study. (D) Comparison between the 22 Pool-Seq populations from Cam2025 and the same samples analyzed with SNPs called using `VarScan` (v2.4.6; Koboldt *et al.*, 2012) (rather than `FreeBayes`) to isolate the effect of SNP caller. The relatively low overlap points to the important impact of SNP calling methods. Unlike `FreeBayes`, which implements a probabilistic haplotype caller based on a Bayesian statistical framework—thus obviating the need for realignment around indels or base quality score recalibration—`VarScan` uses a heuristic-based approach that is more susceptible to sequencing and mapping errors, potentially introducing noise that can dilute true signal and reduce the consistency of detection with the $C_2$ statistic.

### Supplementary Notes

#### N1 Data and methods to construct the new chromosome-level *dsu_isojap1.0* assembly of the *D. suzukii* genome

To generate a novel high-quality chromosome-level *de novo* assembly of the *Drosophila suzukii* genome, we relied on PacBio HiFi long reads in conjunction with HiC sequencing data for scaffolding. For HiFi sequencing, we first extracted high molecular weight (HMW) DNA using the QIAGEN Genomic-tips 500/G kit (Qiagen, MD, USA) from a pool of 30 flash frozen males belonging to an isofemale line of Japanese origin established from females captured on Hachijo Island in 1978 (Ohashi *et al.*, 1991). We used the QIAGEN Genomic-tips 500/G kit (Qiagen, MD, USA) for DNA extraction following the tissue protocol using whole bodies of adults that were flash frozen in liquid nitrogen and then pulverized with a mortar and pestle. After 3h of lysis and one centrifugation step, DNA was immobilized in the column, washed several times, eluted, and finally desalted and concentrated by Isopropyl alcohol precipitation. A final wash in 70% ethanol was performed before resuspending DNA in EB buffer. Quantity and quality analysis of DNA was performed using NanoDrop and Qubit (Thermo Fisher Scientific, MA, USA). DNA integrity was also assessed using the Agilent FP-1002 Genomic DNA 165 kb on the Femto Pulse system (Agilent, CA, USA). Then a PacBio HiFi library was constructed using the SMRTbell Template Prep kit 2.0 (Pacific Biosciences, Menlo Park, CA, USA) according to PacBio recommendations (SMRTbell Express Template Prep kit 2.0 - PN: 100-938-900). Briefly, HMW DNA was first purified with 1X Agencourt AMPure XP beads (Beckman Coulter, Inc., CA USA) and sheared with Megaruptor 3 (Diagenode, Liège, BELGIUM) at an average size of 20 kb. After end repair, A-tailing and ligation of the SMRTbell adapter, the library was selected on BluePippin System (Sage Science, MA,USA) for a range size of 10-50kb. The size and concentration of the library were assessed using Agilent FP-1002 Genomic DNA 165 kb on the Femto Pulse system and the Qubit dsDNA HS reagents assay kit. Sequencing primer v5 and Sequel II DNA Polymerase 2.2 were annealed and bound, respectively, to the SMRTbell library. The library was loaded onto one 8M SMRT cell at an on-plate concentration of 90pM. Sequencing was performed on the Sequel II system on the Gentyane Genomic platform (INRAE Clermont-Ferrand, France) with the Sequel II Sequencing kit 3.0, a run movie time of 30 hours with an Adaptive Loading target (P1 + P2) at 0.75. After filtering and correcting the SMRTcell output, we obtained 743,798 reads totaling 14.7 Gb, with an N50 of 20,851 bp and a maximal length of 50.197 kb as assessed with LongQC v1.2.1 (Fukasawa et al., 2020).

A whole-genome primary assembly was then built from the HiFi reads using `hifiasm` (v0.19.8; Cheng *et al.*, 2021) with the options `--primary` and `--n-hap 8` to account for residual genetic diversity among the pooled individuals included in the HiFi library. Based on preliminary tests, this latter value was found to reduce overall assembly sizes and duplicate rates estimated with BUSCO (Manni *et al.*, 2021, see below). The resulting primary assembly consisted of 167 contigs totaling 305.9 Mb (N50=6.18 Mb), which were further filtered by removing one mitochondrial contig (poorly assembled) and 25 contaminant contigs (i.e., non *D. suzukii*) identified following Gautier (2023) by querying a database constructed from NCBI non-redundant nucleotides (nt) released in February 2020 using the program `Kraken2` (v2.1.2; Wood *et al.*, 2019). In particular, this included a 3.33 Mb contig assigned to *Acetobacter persici*, a 1.35 Mb contig assigned to *Lactobacillus fructivorans*, and a 93.0 kb contig assigned to *Acetobacter pasteurianus.* Finally, we completed the curation of the contig assembly by performing a self-alignment using `minimap2` (v2.20-r1061; Li, 2021) with the options `-xasm5 -DP --dual=no` to identify residual haplotigs. Note that all contigs < 100 kb were discarded as most were found to be haplotigs or junk DNA. The final curated contig assembly consisted of 75 contigs totaling 282,289,391 bp (N50=10.6 Mb and N90=1.79Mb) and included 99.06% (98.48% complete and single) of the 3,285 BUSCO genes of the `diptera_odb10` dataset (v5.4.7; Manni *et al.*, 2021). Following Gautier *et al.* (2018), we were able to identify eight X-linked contigs (totaling 39.14 Mb) and three Y–linked contigs (totaling 16.74 Mb) based on the ratio of the relative mean read coverage of the two pools `RS-Zaj` and `DE-Jen` consisting exclusively of males and females, respectively (see below and Table S1). More precisely, the two corresponding Pool-Seq data set were filtered with `fastp` (v0.23.2; Chen *et al.*, 2018) and mapped to the contig assembly with `bwa-mem2` (v2.2.1; Vasimuddin *et al.*, 2019), both run with default options, and the coverages for each contig were estimated from the resulting *bam* files using `mosdepth` (v0.3.6; Pedersen and Quinlan, 2017) run with option `-Q 20`.

To further scaffold the contig assembly at the chromosome level, we incorporated HiC data, which provides long-range chromatin interaction information. The HiC library was prepared from a pool of three flash-frozen males using the Arima-HiC kit according to the manufacturer's instructions and paired end (2x150bp) sequenced on an Illumina NovaSeq 6000 on the Get Plage platform (INRAE Toulouse, France). The obtained raw paired-end reads were filtered with `fastp` (v0.23.2; Chen *et al.*, 2018) with default options, resulting in a total of 573,783,218 read pairs with an estimated 18.25% of duplicate reads. The filtered paired-end reads were then mapped to the contig assembly using the `HiCstuff` pipeline program (v3.2.1; Baudry et al., 2020; Matthey-Doret et al., 2020) with the options `--mapping=iterative`, `--read-len=150`, `--duplicates`, `--distance-law`, `--filter` and `-e "DpnII,HinfI"`. A total of 148,588,798 valid pairs were successfully identified and used to scaffold the assembly with `yahs` (v1.2; Zhou et al., 2022) and default options, except `{file-type PA5` to read the input *pair* file.

The completeness of the assembly was evaluated with `BUSCO` on the `diptera_odb10` dataset (v5.4.7; Manni *et al.*, 2021). In addition, as mentioned in the main text (see also Table S3), a total of 3,197 of the 3,285 BUSCO genes were found complete and in single copy in both the *dsu_isojap1.0* and the

*Dmel6* assembly for the *D. melanogaster* genome (Hoskins *et al.*, 2015). Moreover, the overlap in BUSCO gene content between the syntenic chromosome arms of the two species was almost complete with only a few discrepancies, aligning with the very close but not perfect conservation of Muller elements among *Drosophila* species (Bhutkar *et al.*, 2008). More specifically, an overlap of 99.4%, 99.9% and 100% was found for chromosomes X, 3 and 4 respectively. The overlap was a bit lower for chromosome arms 2L and 2R (95.8% and 96.0%) respectively, suggesting a less complete sequence. However, an additional large scaffold (40.07 Mb) containing only 19 BUSCO genes, 8 mapping to the 2L and 11 to the 2R *D. melanogaster* chromosome arms, likely corresponded to the (extended) centromeric regions of chromosome 2 (as confirmed by their repeat content, see below and Figure S1) and was thus named *2c*. Similarly, the X scaffold (32.72 Mb in length) actually included seven of the 8 contigs assigned to the X chromosome (see above), the remaining eighth (6.316 Mb) was renamed *scf_Xc* as it was probably more centromeric (see below and Figure S1C). In contrast, none of the three contigs (total 16.74 Mb) assigned to the Y chromosome could be scaffolded. The largest (13.21 Mb) was then named Y, while the other two were manually joined into an 3.338 Mb Y-linked scaffold (*scf_Y1*) based on alignment of the Occludin-Related Y (*Ory*) *D. melanogaster* gene. Note that no BUSCO genes in common with the *Dmel6* assembly mapped to the Y or `scf_Xc` contigs (Table S3).

We annotated repetitive elements using `RepeatMasker` (v4.1.4; Smit *et al.*, 2023) run with options `-s -no_is -cutoff 200 -frag 20000`, the `wublast` search engine (option `-e wublast`) and the curated and annotated TE database developed by (Mérel *et al.*, 2021) for the *D. suzukii* genome. The count distribution of the different TE orders over the scaffolds highlighted distinct chromosomal regions (Figure S1C). In particular, the center of chr3, one of the two extremities of 2L, X and 2R, and the entire chromosomes 4, Y and scf_Y1, scf_2c (93.7%) and scf_Xc scaffolds lacked simple repeats and were enriched in Long Terminal Repeat retrotransposons and Rolling Circle transposons. This suggests that such TE signature specifies centromeric regions in the *D. suzukii* genome, as shown by Chang *et al.* (2019) in *D. melanogaster*, although different TE superfamilies are involved. The assembly *dsu_isojap1.0* was finally submitted to the NCBI repository for gene annotation with the NCBI Eukaryotic Genome Annotation Pipeline (Thibaud-Nissen *et al.*, 2013). A total of 17,936 genes (including 15,749 coding and 2,219 noncoding) and 266 nontranscribed pseudogenes were identified. Most annotated genes were mapped to the identified autosomal scaffolds, with 79.3% located on chromosomes 2L, 2R, 3, 4 and scf_2c (Figure S3A). Similarly, 14.5% of genes were mapped to the X-linked scaffold (chrX) and scf_Xc. A total of 404 genes (2.25%) were annotated on the Y-linked scaffolds (chrY and scf_Y1), while 712 genes (3.97%) were assigned to the remaining small scaffolds. Furthermore, along the main chromosomes, the gene density was lower in the centromeric regions (e.g., middle of chr3), which were identified based on their TE content, as well as in the other scaffolds, including scf_2c and scf_Xc (Figure S3B).

#### N2 *D. suzukii* sample curation

Following Gautier (2023), we removed all samples with a high level of contaminating sequences, except for the CN-Nin sample (collected in Ningbo, China). This latter sample indeed remained highly valuable as it represented a native population closely related to one of the contributing sources of the America invasion identified by Fraimout *et al.* (2017). Although this sample exhibited 15% *D. subpulchrella* contamination (i.e., 7 *D. subpulchrella* out of the 50 pooled individuals), we do not expect any strong bias introduced into our global analyses, since the *D. subpulchrella* specific variants may remain at very low frequency in the total analyzed data sets. Accordingly, the CN-Nin sample was found to be closely related to other samples from China in Figures 2B or 2C. We considered only two (JP-Kan and US-CAOx) of the seven Pool-Seq samples from Feng *et al.* (2024), the others being either already sequenced samples (n=3) or closely related geographically (n=2) to those from Olazcuaga *et al.* (2020). Finally, we selected a subset of the 235 individuals sequenced by Lewald *et al.* (2021) based on both geographic and preliminary genetic analyses to limit redundancy in the representation of US populations and to ensure a minimum sample size per population (from n=7 to n=10 individuals). Note that the samples representing the US states of Georgia (US-Ga, n=9), New York (US-Ny, n=10), and Washington (US-Wa, n=10) each combined individuals collected simultaneously from several neighboring sampling sites located at a maximum of 27 km, 45 km, and 15 km apart, respectively (Table S2).

#### N3 Variant calling

To perform variant calling, FreeBayes (v1.3.6; Garrison and Marth, 2012) was used with options -K (to output all alleles that pass input filters, regardless of genotyping outcome, assuming a pooled sequencing model); -C 1 -F 0.01 (at least one count and a fraction of 1% of the counts of the alternate allele to evaluate the position); -G 5 (at least 5 counts supporting an alternate allele in all sample to retain the allele); -E -1 (to disable clumping of contiguous variants into complex alleles); --limit-coverage 500 (to downsample per-sample coverage to 500 reads if greater than this coverage); -n 4 (to evaluate only the best 4 alleles, ranked by sum of supporting quality scores), -m 30 -q 20 (minimum mapping and base qualities set to 30 and 20, respectively). For X-linked contigs, we also specified the ploidy of the 5 male individuals with the *-A* option. Variant calling was performed in parallel on a computer grid for the 1,179 nonoverlapping 250 kb windows covering the whole genome assembly. The resulting vcf files were reordered and concatenated per chromosome type using the view and concat programs of bcftools (v1.20; Danecek et al., 2021), and filtered using a custom awk script to keep only bi-allelic variants (including the reference allele) with QUAL>20, NS>100, 100<DP<5000, SAF>0, SAR>0, RPR>1, RPL>1, EPP>0, SRP>0, and a global MAF computed from the overall read counts $> 0.01$.

### N4 New updates to the BayPass model to analyze Genotype Likelihoods and hybrid data sets

#### N4.1 Modeling Genotype Likelihoods

Let each genotype be coded by its number of alternate allele $l$, i.e., for a diploid individual $l = 0$ if homozygote for the reference allele, $l = 1$ if heterozygote, or $l = 2$ if homozygote for the alternate allele. For haploid individuals (e.g., males at X–linked SNPs), only the two values $l = 0$ or $l = 1$ are observable. Further, let $GL_{ijkl} = \mathrm{P}\left(y_{ijk}^{(ind)} \mid g_{ijk} = l\right)$ denote the likelihood, computed by the genotype caller, of the observed Ind-Seq data for the $k$th individual from population $j$ at locus $i$ when its genotypes $g_{ijk} = l$. Following the same notations as in Gautier (2015) and assuming Hardy-Weinberg equilibrium, the (conditional) probability of the observed data may then simply be defined as a function of the (reference) allele frequency $\alpha_{ij}$ at a bi-allelic locus $i$ in population $j$ and individual ploidy $p_{ijk}$ ($p_{ijk}$=1 or 2 if individual $k$ from population $j$ is haploid or diploid at SNP $i$, respectively), as $\mathrm{P}\left(y_{ijk}^{(ind)} \mid \alpha_{ij}\right) = \sum\limits_{l=0}^{p_{ijk}} \binom{p_{ijk}}{l} \alpha_{ij}^{p_{ijk}-l} \left(1-\alpha_{ij}\right)^{l} GL_{ijkl}$. Finally, assuming that individuals are randomly sampled (i.e., conditional independence), the conditional probability of the observed data $Y_{ij}^{(ind)} = \left\{y_{ijk}^{(ind)}\right\}$ at SNP $i$ for the $n_j$ individuals from population $j$ is:

$$f\left(Y_{ij}^{(ind)} \mid \alpha_{ij}\right) = \prod_{k=1}^{n_j} \mathrm{P}\left(y_{ijk}^{(ind)} \mid \alpha_{ij}\right) = \prod_{k=1}^{n_j} \left(\sum_{l=0}^{p_{ijk}} \binom{p_{ijk}}{l} \alpha_{ij}^{p_{ijk}-l} \left(1-\alpha_{ij}\right)^{l} GL_{ijkl}\right) \tag{N1}$$

This likelihood may then easily be incorporated at the first level of the Bayesian hierarchical models described in Gautier (2015).

#### N4.2 Description of the MCMC algorithm used to sample of the $\alpha_{ij}^{\star}$'s

The sampling of $\alpha_{ij}$'s from the full posterior distribution was implemented within the BayPass Markov Chain Monte Carlo (MCMC) algorithm using a random walk Metropolis update similar to the alternate algorithm 1 described in Supplementary File S1 of Gautier (2015). Briefly, the $\alpha_{ij}^{\star}$'s are iteratively updated in each population, one locus at a time, by sampling from their full conditional distribution which has the following general form (using the same notations as in Gautier, 2015):

$$f\left(\alpha_{ij}^{\star} \mid .\right) \propto f\left(\widetilde{\alpha}_{ij}^{\star} \mid \widetilde{\boldsymbol{\alpha}}_{i,-j}^{\star}, \boldsymbol{\Lambda}\right) f\left(Y_{ij}^{(ind)} \mid \alpha_{ij}\right)$$

where:

- $\widetilde{\boldsymbol{\alpha}}_{\boldsymbol{i}}^{\star} = \left\{\frac{\alpha_{ij}^{\star} - \pi_i - \delta_i \beta_i \boldsymbol{Z}_j}{\sqrt{\pi(1-\pi_i)}}\right\}_{(1..J)}$ that is $\widetilde{\boldsymbol{\alpha}}_{\boldsymbol{i}}^{\star} \mid \boldsymbol{\Lambda} \sim \mathrm{N}_J\left(\mathbf{I}_{\mathrm{J}}; \boldsymbol{\Omega} = \boldsymbol{\Lambda}^{-1}\right)$
- $f\left(Y_{ij}^{(ind)} \mid \alpha_{ij}\right)$ is as defined in Equation N1 above
- $\alpha_{ij} = \max\left(0; \min\left(1; \alpha_{ij}^{\star}\right)\right)$

From the properties of the multivariate Gaussian distribution: $\widetilde{\alpha}^{\star}_{ij} \mid \widetilde{\boldsymbol{\alpha}}^{\star}_{\boldsymbol{i,-j}}, \boldsymbol{\Lambda} \sim N\left(\mu_{(\alpha),ij}, \sigma^2_{(\alpha),ij}\right)$, where $\mu_{(\alpha),ij} = \mathbf{C}\widetilde{\boldsymbol{\alpha}}_{\boldsymbol{i,-j}}$ and $\sigma^2_{(\alpha),ij} = \omega_{jj} - \mathbf{C}\boldsymbol{\Omega}_{\boldsymbol{kj}}$. Here, $\mathbf{C} = \boldsymbol{\Omega}_{\boldsymbol{jk}}\boldsymbol{\Omega}^{-1}_{\boldsymbol{kk}}$ represents the matrix of regression coefficients, and $\boldsymbol{\Omega}_{\boldsymbol{kj}}$, $\boldsymbol{\Omega}_{\boldsymbol{kk}}$, and $\boldsymbol{\Omega}_{\boldsymbol{jj}}$ are blocks of the matrix $\boldsymbol{\Omega}$. For instance, for $j = 1$:

$$\boldsymbol{\Omega} = \begin{pmatrix} \omega_{11} & \boldsymbol{\Omega_{12}} \\ \boldsymbol{\Omega_{21}} & \boldsymbol{\Omega_{22}} \end{pmatrix}$$

As a consequence:

$$f\left(\alpha^{\star}_{ij} \mid .\right) \propto e^{-\frac{1}{2\sigma^2_{(\alpha),ij}}\left(\alpha^{\star}_{ij} - \mu_{(\alpha),ij}\right)^2} \times f\left(Y^{(ind)}_{ij} \mid \alpha_{ij}\right)$$

Because this full conditional is not of usual form, a (random-walk) Metropolis update is implemented. A candidate $\widetilde{\alpha}^{\star,(c)}_{ij}$ is sampled from the following uniform distribution $\text{Unif}\left(\widetilde{\alpha}^{\star}_{ij} - \delta^{(\alpha)}_{ij}; \widetilde{\alpha}^{\star}_{ij} + \delta^{(\alpha)}_{ij}\right)$ where $\delta^{(\alpha)}_{ij}$ is adjusted for each $\alpha^{\star}_{ij}$ during pilot runs (to obtain acceptance rates ranging between $\tau_{\min} = 0.25$ and $\tau_{\max} = 0.4$, by default, see the BayPass manual for details). As this proposal distribution is symmetric, the candidate value $\widetilde{\alpha}^{\star,(c)}_{ij}$ is accepted with probability $min\left(1, \psi^{(\alpha)}_{ij} = \frac{f\left(\alpha^{\star,(c)}_{ij} \mid .\right)}{f\left(\alpha^{\star}_{ij} \mid .\right)}\right)$ according to the Metropolis rule.

#### N4.3 Analyzing hybrid data sets

Capitalizing on the versatility of Bayesian hierarchical modeling, the new version 3.0 of BayPass was modified to allow joint analysis of allele count and / or read count (e.g. Pool-Seq) and / or GL data. More precisely, let $\boldsymbol{Y} = \left\{\left\{Y^{(ac)}_{ij}\right\}_{j\in\left[1,J^{(ac)}\right]}; \left\{Y^{(rc)}_{ij'}\right\}_{j'\in\left[1,J^{(rc)}\right]}; \left\{Y^{(ind)}_{ij''}\right\}_{j''\in\left[1,J^{(ind)}\right]}\right\}_{i\in[1,nsnp]}$ a genomic data set consisting of:

- Sample allele counts $Y^{(ac)}$ for $J^{(ac)}$ sampled populations, typically coded in BayPass as a matrix with $n_{snp}$ rows and $2 \times J^{(ac)}$ columns containing for each SNP and each population a pair of entries corresponding to the observed counts for the reference (i.e., sums of the individual genotypes coded as above) and alternate alleles, respectively.

- Sample read counts (e.g., from Pool-Seq experiment) $Y^{(rc)}$ for $J^{(rc)}$ sampled populations typically coded in BayPass as a matrix with $n_{snp}$ rows and $2 \times J^{(rc)}$ columns containing for each SNP and each population a pair of entries corresponding to the observed read counts for the reference and alternate alleles, respectively.

- Individuals' GLs (e.g., from Ind-Seq experiment) $Y^{(ind)}$ available for $J^{(ind)}$ sampled populations typically coded in BayPass as a (character) matrix with $n_{snp}$ rows and $n_{ind}$ columns (e.g., $n_{ind} = 15 \times J^{(ind)}$ if 15 individuals per sampled populations were sequenced) containing the GL triplet (or doublet if haploid) for each SNP and each individual.

Assuming conditional independence of markers (i.e., not accounting for LD), and given the underlying population allele frequencies $\boldsymbol{\alpha} = \left\{\{\alpha_{ij}\}_j; \{\alpha_{ij'}\}_{j'}; \{\alpha_{ij''}\}_{j''}\right\}_i$, the full likelihood of the combined data set $\boldsymbol{Y}$ becomes:

$$f\left(\boldsymbol{Y} \mid \boldsymbol{\alpha}\right) = \prod_{i=1}^{nsnp} \left( \prod_{j=1}^{J^{(ac)}} f\left(Y_{ij}^{(ac)} \mid \alpha_{ij}\right) \prod_{j'=1}^{J^{(rc)}} f\left(Y_{ij'}^{(rc)} \mid \alpha_{ij'}\right) \prod_{j''=1}^{J^{(ind)}} f\left(Y_{ij''}^{(ind)} \mid \alpha_{ij''}\right) \right)$$

, where $f\left(Y_{ij}^{(ac)} \mid \alpha_{ij}\right)$ and $f\left(Y_{ij}^{(ac)} \mid \alpha_{ij}\right)$ are the likelihoods previously defined in Gautier (2015) for allele count and read count data, respectively, and $f\left(Y_{ij''}^{(ind)} \mid \alpha_{ij''}\right)$ is the likelihood based on GL's defined above. Then, the sampling of all $\alpha_{ij}$' s from the complete posterior distribution is performed using MCMC as described in Gautier (2015) for the $\alpha_{ij}$' s associated with the count or read count data, and as described above for the $\alpha_{ij}$' s associated with the GL data (see the BayPass manual for additional details).

#### N5 Evaluation of the significance of the overlap of candidate genomic windows

To evaluate the expected (approximate) number of genomic windows that overlap between two independent tests, we considered all possible pairs $(R_1, R_2)$, where $R_i$ is a region detected by test number $i$, whose size in bp is denoted $l_i$. The two regions overlap when $R_2$ starts within $R_1$, which occurs with probability $l_1/L$ where $L$ is the total genome size, or less than $l_2$ bp ahead of $R_1$, which occurs with probability $l_2/L$. The total number of expected overlaps was thus computed as the sum of $(l_1 + l_2)/L$ over all possible pairs. To avoid under-estimating this number due to the existence of large parts of the genome associated with a limited detection power, only chromosomes arms where signals had been actually detected were accounted when computing $L$.

### N6 Environmental variables for Genomic Offset computation

The 29 environmental variables used in the GEA to estimate regression coefficients and compute gGO can be grouped in 3 data type :

- Bioclimatic variables: Included 19 standard variables, along with the mean monthly climate moisture (highlighting humidity's role in fitness-related traits in *D. suzukii* Winkler *et al.* (2020)), and the mean monthly near-surface wind speed (potential main driver of adaptation in *D. melanogaster* European populations (Bogaerts-Márquez et al., 2021)). All data (averaged over 1981–2010 to represent the environment of each sampling location by capturing long-term trends while minimizing interannual variability) were extracted from the CHELSA database (Karger et al., 2017) using the R packages `dismo` (Hijmans et al., 2023) and `raster` (Hijmans, 2022).

- Land-use variables: based on the ESRI Land Cover 2020 dataset (Karra et al., 2021), we included percentage cover of six classes (crops, built areas, trees, water, bareground, rangeland). Other classes (ice/snow, clouds, flooded vegetation) were discarded due to lack of variation across sites.

- Agricultural production variables: Given the species' dependency on fruit availability (whether cultivated or wild), we aimed to incorporate host fruit presence into our GEA. However, due to the extreme generalism of the species (Poyet *et al.*, 2015; Olazcuaga et al., 2023), and the lack of fine-scale, host-specific data at the global level, we used global agricultural production as a proxy. Specifically, we extracted data from the MIRCA-OS database (Kebede et al., 2025) on annual harvested area (in hectares) for irrigated and rainfed crops in the "Other perennials" category for the year 2015. This group includes many crops that are known host plants for *D. suzukii* (e.g., cranberries, berries, blueberries, and strawberries).

### N7 Details about noteworthy candidate genes

Among the top 45 genes identified through our genome scan approach, several are known to be involved in biologically relevant functions,, including stress resistance and detoxification. For instance, in *D. melanogaster*, *CG30015* is likely involved in heavy metal susceptibility (Zhou et al., 2017), *Olf413* has been identified as a candidate for the toxicological response to monoterpene pesticides (Sabio et al., 2024), and *Prosalpha2* likely plays a role in a genetic resistance network to pesticides (Zhang and Zhang, 2019). Additionally, both *ChChd2* and *ClpX* may contribute to the protection against oxidative stress (Meng et al., 2017; Hill et al., 2018). Together, these genes may support adaptive responses to environmental stressors and anthropogenic pressures, including exposure to pesticides, potentially facilitating the establishment of *D. suzukii* in agricultural landscapes.

Several genes related to dispersal and movement were also among the top 45 candidates, such as i) *CG34215* found associated with flight performance in *D. melanogaster* (Spierer et al., 2021); ii) *Oamb* which influences locomotion, climbing ability, and exploratory behaviors (El-Kholy et al., 2021); iii) *Hmgcr* involved in sexual dimorphism of locomotor behavior (Belgacem and Martin, 2007); and iv) *Ptr* involved in neuromuscular function during larval development and likely adult lifespan regulation (Parada and Prieto, 2024). Likewise, some of the top candidate genes involved in chemosensation and feeding behavior, such as i) *Cheb42c* which is implicated in pheromone perception (Xu et al., 2002); ii) *Gr57a*, a gustatory receptor found associated with variations in olfactory perception in *D. melanogaster* (Arya et al., 2015); iii) *Obp56h*, which is linked to mating and feeding behavior (Shorter et al., 2016; Delclos et al., 2024; Swarup et al., 2014) and is differentially expressed between summer and winter morphs in *D. suzukii*, suggesting a role in seasonal feeding adjustments (Schwanitz et al., 2022); and iv) *Ppk11* that influences salt taste perception (Liu et al., 2003).

We also identified candidate genes related to immune response, such as i) *CG34215* found to be differentially expressed in *D. suzukii* following infection by the entomopathogenic nematode *Steinernema carpocapsae* (Garriga et al., 2024); and ii) *Psq*, identified as a potential target of *Wolbachia*-induced cytoplasmic incompatibility in *D. melanogaster* (Zheng et al., 2019). Finally, the *CG30015* and *Oamb* candidate genes have been found involved in social behaviors associated with aggression in *D. melanogaster* (Edwards et al., 2006; Luo et al., 2014).

It is worth stressing that most of the genes mentioned above have been found in candidate regions where there are not any or very few other genes. Therefore, there is good statistical evidence that these genes have been the drivers of the selection signal that has been detected in these regions.

### Supplementary Material References


Arya, G. H., Magwire, M. M., Huang, W., Serrano-Negron, Y. L., Mackay, T. F., and Anholt, R. R. (2015). The genetic basis for variation in olfactory behavior in drosophila melanogaster. *Chemical senses*, 40(4):233–243.

Baudry, L., Guiglielmoni, N., Marie-Nelly, H., Cormier, A., Marbouty, M., Avia, K., Mie, Y. L., Godfroy, O., Sterck, L., Cock, J. M., Zimmer, C., Coelho, S. M., and Koszul, R. (2020). instaGRAAL: chromosome-level quality scaffolding of genomes using a proximity ligation-based scaffolder. *Genome Biol*, 21(1):148.

Belgacem, Y. H. and Martin, J.-R. (2007). Hmgcr in the corpus allatum controls sexual dimorphism of locomotor activity and body size via the insulin pathway in drosophila. *PLoS One*, 2(1):e187.

Bhutkar, A., Schaeffer, S. W., Russo, S. M., Xu, M., Smith, T. F., and Gelbart, W. M. (2008). Chromosomal rearrangement inferred from comparisons of 12 Drosophila genomes. *Genetics*, 179(3):1657–1680.

Bogaerts-Márquez, M., Guirao-Rico, S., Gautier, M., and González, J. (2021). Temperature, rainfall and wind variables underlie environmental adaptation in natural populations of Drosophila melanogaster. *Molecular Ecology*, 30(4):938–954.

Cabanettes, F. and Klopp, C. (2018). D-GENIES : Dot plot large GENomes in an interactive, efficient and simple way. *PeerJ Preprints*, 6:e26567v1.

Challis, R., Richards, E., Rajan, J., Cochrane, G., and Blaxter, M. (2020). BlobToolKit – Interactive Quality Assessment of Genome Assemblies. *G3*, 10(4):1361–1374.

Chang, C.-H., Chavan, A., Palladino, J., Wei, X., Martins, N. M. C., Santinello, B., Chen, C.-C., Erceg, J., Beliveau, B. J., Wu, C.-T., Larracuente, A. M., and Mellone, B. G. (2019). Islands of retroelements are major components of drosophila centromeres. *PLOS Biology*, 17(5):1–40.

Chen, S., Zhou, Y., Chen, Y., and Gu, J. (2018). fastp: an ultra-fast all-in-one fastq preprocessor. *Bioinformatics*, 34(17):i884–i890.

Cheng, H., Concepcion, G. T., Feng, X., Zhang, H., and Li, H. (2021). Haplotype-resolved de novo assembly using phased assembly graphs with hifiasm. *Nature Methods*, 18(2):170–175.

Danecek, P., Bonfield, J. K., Liddle, J., Marshall, J., Ohan, V., Pollard, M. O., Whitwham, A., Keane, T., McCarthy, S. A., Davies, R. M., and Li, H. (2021). Twelve years of SAMtools and BCFtools. *GigaScience*, 10(2):giab008.

Delclos, P. J., Adhikari, K., Mai, A. B., Hassan, O., Oderhowho, A. A., Sriskantharajah, V., Trinh, T., and Meisel, R. (2024). Trans regulation of an odorant binding protein by a proto-y chromosome affects male courtship in house fly. *Elife*, 13:e90349.

Edwards, A. C., Rollmann, S. M., Morgan, T. J., and Mackay, T. F. C. (2006). Quantitative genomics of aggressive behavior in drosophila melanogaster. *PLoS genetics*, 2(9):e154.

El-Kholy, S. E., Afifi, B., El-Husseiny, I., and Seif, A. (2021). Octopamine signaling via oamb is essential for a well-orchestrated climbing performance of adult drosophila melanogaster.

Feng, S., DeGrey, S. P., Guédot, C., Schoville, S. D., and Pool, J. E. (2024). Genomic Diversity Illuminates the Environmental Adaptation of Drosophila suzukii. *Genome Biology and Evolution*, 16(9):evae195.

Fraimout, A., Debat, V., Fellous, S., Hufbauer, R. A., Foucaud, J., Pudlo, P., Marin, J.-M., Price, D. K., Cattel, J., Chen, X., Deprá, M., François Duyck, P., Guedot, C., Kenis, M., Kimura, M. T., Loeb, G., Loiseau, A., Martinez-Sañudo, I., Pascual, M., Polihronakis Richmond, M., Shearer, P., Singh, N., Tamura, K., Xuéreb, A., Zhang, J., and Estoup, A. (2017). Deciphering the Routes of invasion of Drosophila suzukii by Means of ABC Random Forest. *Molecular Biology and Evolution*, 34(4):980–996.

Fukasawa, Y., Ermini, L., Wang, H., Carty, K., and Cheung, M.-S. (2020). Longqc: a quality control tool for third generation sequencing long read data. *G3: Genes, Genomes, Genetics*, 10(4):1193–1196.

Garriga, A., Toubarro, D., Morton, A., Simões, N., and García-del Pino, F. (2024). Analysis of the immune transcriptome of the invasive pest spotted wing drosophila infected by steinernema carpocapsae. *Bulletin of Entomological Research*, 114(5):622–630.

Garrison, E. and Marth, G. (2012). Haplotype-based variant detection from short-read sequencing. *arXiv*, 1207.3907.

Gautier, M. (2015). Genome-Wide Scan for Adaptive Divergence and Association with Population-Specific Covariates. *Genetics*, 201(4):1555–1579.

Gautier, M. (2023). Efficient *k-mer* based curation of raw sequence data: application in *Drosophila suzukii*. *Peer Community Journal*, 3:e79.

Gautier, M., Yamaguchi, J., Foucaud, J., Loiseau, A., Ausset, A., Facon, B., Gschloessl, B., Lagnel, J., Loire, E., Parrinello, H., Severac, D., Lopez-Roques, C., Donnadieu, C., Manno, M., Berges, H., Gharbi, K., Lawson-Handley, L., Zang, L.-S., Vogel, H., Estoup, A., and Prud'homme, B. (2018). The Genomic Basis of Color Pattern Polymorphism in the Harlequin Ladybird. *Current Biology*, 28(20):3296–3302.e7.

Hijmans, R. J. (2022). *raster: Geographic Data Analysis and Modeling*. R package version 3.5-15.

Hijmans, R. J., Phillips, S., Leathwick, J., and Elith, J. (2023). *dismo: Species Distribution Modeling*. R package version 1.3-14.

Hill, V. M., O'Connor, R. M., Sissoko, G. B., Irobunda, I. S., Leong, S., Canman, J. C., Stavropoulos, N., and Shirasu-Hiza, M. (2018). A bidirectional relationship between sleep and oxidative stress in drosophila. *PLoS biology*, 16(7):e2005206.

Hoskins, R. A., Carlson, J. W., Wan, K. H., Park, S., Mendez, I., Galle, S. E., Booth, B. W., Pfeiffer, B. D., George, R. A., Svirskas, R., Krzywinski, M., Schein, J., Accardo, M. C., Damia, E., Messina, G., Méndez-Lago, M., de Pablos, B., Demakova, O. V., Andreyeva, E. N., Boldyreva, L. V., Marra, M., Carvalho, A. B., Dimitri, P., Villasante, A., Zhimulev, I. F., Rubin, G. M., Karpen, G. H., and Celniker, S. E. (2015). The release 6 reference sequence of the drosophila melanogaster genome. *Genome Research*, 25(3):445–58.

Karger, D. N., Conrad, O., Böhner, J., Kawohl, T., Kreft, H., Soria-Auza, R. W., Zimmermann, N. E., Linder, H. P., and Kessler, M. (2017). Climatologies at high resolution for the earth's land surface areas. *Scientific Data*, 4(1):170122.

Karra, K., Kontgis, C., Statman-Weil, Z., Mazzariello, J. C., Mathis, M., and Brumby, S. P. (2021). Global land use/land cover with Sentinel 2 and deep learning. *2021 IEEE International Geoscience and Remote Sensing Symposium IGARSS*, pages 4704–4707.

Kebede, E. A., Oluoch, K. O., Siebert, S., Mehta, P., Hartman, S., Jägermeyr, J., Ray, D., Ali, T., Brauman, K. A., Deng, Q., et al. (2025). A global open-source dataset of monthly irrigated and rainfed cropped areas (mirca-os) for the 21st century. *Scientific Data*, 12(1):208.

Koboldt, D. C., Zhang, Q., Larson, D. E., Shen, D., McLellan, M. D., Lin, L., Miller, C. A., Mardis, E. R., Ding, L., and Wilson, R. K. (2012). Varscan 2: somatic mutation and copy number alteration discovery in cancer by exome sequencing. *Genome Research*, 22(3):568–76.

Lewald, K. M., Abrieux, A., Wilson, D. A., Lee, Y., Conner, W. R., Andreazza, F., Beers, E. H., Burrack, H. J., Daane, K. M., Diepenbrock, L., Drummond, F. A., Fanning, P. D., Gaffney, M. T., Hesler, S. P., Ioriatti, C., Isaacs, R., Little, B. A., Loeb, G. M., Miller, B., Nava, D. E., Rendon, D., Sial, A. A., Bezerra da Silva, C. S., Stockton, D. G., Van Timmeren, S., Wallingford, A., Walton, V. M., Wang, X., Zhao, B., Zalom, F. G., and Chiu, J. C. (2021). Population genomics of Drosophila suzukii reveal longitudinal population structure and signals of migrations in and out of the continental United States. *G3*, 11(12):jkab343.

Li, H. (2021). New strategies to improve minimap2 alignment accuracy. *Bioinformatics*, 37(23):4572–4574.

Liu, L., Leonard, A. S., Motto, D. G., Feller, M. A., Price, M. P., Johnson, W. A., and Welsh, M. J. (2003). Contribution of drosophila deg/enac genes to salt taste. *Neuron*, 39(1):133–146.

Luo, J., Lushchak, O. V., Goergen, P., Williams, M. J., and Nässel, D. R. (2014). Drosophila insulin-producing cells are differentially modulated by serotonin and octopamine receptors and affect social behavior. *PloS one*, 9(6):e99732.

Manni, M., Berkeley, M. R., Seppey, M., Simão, F. A., and Zdobnov, E. M. (2021). BUSCO Update: Novel and Streamlined Workflows along with Broader and Deeper Phylogenetic Coverage for Scoring of Eukaryotic, Prokaryotic, and Viral Genomes. *Molecular Biology and Evolution*, 38(10):4647–4654.

Matthey-Doret, C., Baudry, L., Bignaud, A., Cournac, A., Remi-Montagne, Guiglielmoni, N., Foutel-Rodier, T., and Scolari, V. F. (2020). hicstuff: Simple library/pipeline to generate and handle hi-c data.

Meng, H., Yamashita, C., Shiba-Fukushima, K., Inoshita, T., Funayama, M., Sato, S., Hatta, T., Natsume, T., Umitsu, M., Takagi, J., et al. (2017). Loss of parkinson's disease-associated protein chchd2 affects mitochondrial crista structure and destabilizes cytochrome c. *Nature communications*, 8(1):15500.

Mérel, V., Gibert, P., Buch, I., Rodriguez Rada, V., Estoup, A., Gautier, M., Fablet, M., Boulesteix, M., and Vieira, C. (2021). The Worldwide Invasion of Drosophila suzukii Is Accompanied by a Large Increase of Transposable Element Load and a Small Number of Putatively Adaptive Insertions. *Molecular Biology and Evolution*, 38(10):4252–4267.

Ohashi, Y. Y., Haino-Fukushima, K., and Fuyama, Y. (1991). Purification and charachterization of an ovulation stimulating substance from the male accessory glands of drosophila suzukii. *Insect Biochemistry*, 21(4):413–419.

Olazcuaga, L., Baltenweck, R., Leménager, N., Maia-Grondard, A., Claudel, P., Hugueney, P., and Foucaud, J. (2023). Metabolic consequences of various fruit-based diets in a generalist insect species. *Elife*, 12:e84370.

Olazcuaga, L., Loiseau, A., Parrinello, H., Paris, M., Fraimout, A., Guedot, C., Diepenbrock, L. M., Kenis, M., Zhang, J., Chen, X., Borowiec, N., Facon, B., Vogt, H., Price, D. K., Vogel, H., Prud'homme, B., Estoup, A., and Gautier, M. (2020). A Whole-Genome Scan for Association with Invasion Success in the Fruit Fly Drosophila suzukii Using Contrasts of Allele Frequencies Corrected for Population Structure. *Molecular Biology and Evolution*, 37(8):2369–2385.

Parada, C. and Prieto, D. (2024). Survival, movement, and lifespan: Decoding the roles of patched-related (ptr) in drosophila melanogaster. *bioRxiv*, pages 2024–05.

Paris, M., Boyer, R., Jaenichen, R., Wolf, J., Karageorgi, M., Green, J., Cagnon, M., Parinello, H., Estoup, A., Gautier, M., Gompel, N., and Prud'homme, B. (2020). Near-chromosome level genome assembly of the fruit pest Drosophila suzukii using long-read sequencing. *Scientific Reports*, 10(1):11227.

Pedersen, B. S. and Quinlan, A. R. (2017). Mosdepth: quick coverage calculation for genomes and exomes. *Bioinformatics*, 34(5):867–868.

Poyet, M., Le Roux, V., Gibert, P., Meirland, A., Prevost, G., Eslin, P., and Chabrerie, O. (2015). The wide potential trophic niche of the asiatic fruit fly drosophila suzukii: the key of its invasion success in temperate europe? *PloS one*, 10(11):e0142785.

Sabio, M. C., Alzogaray, R., and Fanara, J. J. (2024). Genetic architecture of the toxicological response to eucalyptol and citronellal in drosophila melanogaster. *Pesticide Biochemistry and Physiology*, 202:105938.

Sario, S., Marques, J. P., Farelo, L., Afonso, S., Santos, C., and Melo-Ferreira, J. (2024). Dissecting the invasion history of spotted-wing drosophila (drosophila suzukii) in portugal using genomic data. *BMC Genomics*, 25(1):813.

Schwanitz, T. W., Polashock, J. J., Stockton, D. G., Rodriguez-Saona, C., Sotomayor, D., Loeb, G., and Hawkings, C. (2022). Molecular and behavioral studies reveal differences in olfaction between winter and summer morphs of drosophila suzukii. *PeerJ*, 10:e13825.

Shorter, J. R., Dembeck, L. M., Everett, L. J., Morozova, T. V., Arya, G. H., Turlapati, L., St. Armour, G. E., Schal, C., Mackay, T. F., and Anholt, R. R. (2016). Obp56h modulates mating behavior in drosophila melanogaster. *G3: Genes, Genomes, Genetics*, 6(10):3335–3342.

Smit, A., Hubley, R., and Green, P. (2023). *RepeatMasker Open-4.0.* Institute for Systems Biology.

Spierer, A. N., Mossman, J. A., Smith, S. P., Crawford, L., Ramachandran, S., and Rand, D. M. (2021). Natural variation in the regulation of neurodevelopmental genes modifies flight performance in drosophila. *PLoS Genetics*, 17(3):e1008887.

Swarup, S., Morozova, T. V., Sridhar, S., Nokes, M., and Anholt, R. R. (2014). Modulation of feeding behavior by odorant-binding proteins in drosophila melanogaster. *Chemical senses*, 39(2):125–132.

Thibaud-Nissen, F., Souvorov, A., Murphy, T., DiCuccio, M., and Kitts, P. (2013). Eukaryotic genome annotation pipeline. In *The NCBI Handbook*. National Center for Biotechnology Information (US). https://www.ncbi.nlm.nih.gov/books/NBK169439/.

Vasimuddin, M., Misra, S., Li, H., and Aluru, S. (2019). Efficient architecture-aware acceleration of bwa-mem for multicore systems. In *2019 IEEE International Parallel and Distributed Processing Symposium (IPDPS)*, pages 314–324.

Winkler, A., Jung, J., Kleinhenz, B., and Racca, P. (2020). A review on temperature and humidity effects on Drosophila suzukii population dynamics. *Agricultural and Forest Entomology*, 22(3):179–192.

Wood, D. E., Lu, J., and Langmead, B. (2019). Improved metagenomic analysis with kraken 2. *Genome Biology*, 20(1):257.

Xu, A., Park, S.-K., D'Mello, S., Kim, E., Wang, Q., and Pikielny, C. (2002). Novel genes expressed in subsets of chemosensory sensilla on the front legs of male drosophila melanogaster. *Cell and tissue research*, 307:381–392.

Zhang, G. and Zhang, W. (2019). Protein–protein interaction network analysis of insecticide resistance molecular mechanism in drosophila melanogaster. *Archives of Insect Biochemistry and Physiology*, 100(1):e21523.

Zheng, Y., Shen, W., Bi, J., Chen, M.-Y., Wang, R.-F., Ai, H., and Wang, Y.-F. (2019). Small rna analysis provides new insights into cytoplasmic incompatibility in drosophila melanogaster induced by wolbachia. *Journal of Insect Physiology*, 118:103938.

Zhou, C., McCarthy, S. A., and Durbin, R. (2022). YaHS: yet another Hi-C scaffolding tool. *Bioinformatics*, 39(1):btac808.

Zhou, S., Luoma, S. E., St. Armour, G. E., Thakkar, E., Mackay, T. F., and Anholt, R. R. (2017). A drosophila model for toxicogenomics: Genetic variation in susceptibility to heavy metal exposure. *PLoS genetics*, 13(7):e1006907.
